# Evaluation of surface-based hippocampal registration using ground-truth subfield definitions

**DOI:** 10.1101/2023.03.30.534978

**Authors:** Jordan DeKraker, Nicola Palomero-Gallagher, Olga Kedo, Neda Ladbon-Bernasconi, Sascha E.A. Muenzing, Markus Axer, Katrin Amunts, Ali R. Khan, Boris Bernhardt, Alan C. Evans

## Abstract

The hippocampus is an archicortical structure, consisting of subfields with unique circuits. Understanding its microstructure, as proxied by these subfields, can improve our mechanistic understanding of learning and memory and has clinical potential for several neurological disorders. One prominent issue is how to parcellate, register, or retrieve homologous points between two hippocampi with grossly different morphologies. Here, we present a surface-based registration method that solves this issue in a contrast-agnostic, topology-preserving manner. Specifically, the entire hippocampus is first analytically unfolded, and then samples are registered in 2D unfolded space based on thickness, curvature, and gyrification. We demonstrate this method in seven 3D histology samples and show superior alignment with respect to subfields using this method over more conventional registration approaches.

**Highlights:**

- Hippocampal subfields contain microcircuits that are critical for memory and vulnerable to neurological disease.
- Hippocampi have variable folding patterns between individuals, making them hard to register or parcellate.
- We present a surface-based hippocampal registration method that is analogous to neocortical inflation to a sphere and registration.
- Testing in seven detailed 3D histology samples revealed successful registration with respect to hippocampal subfields, and outperformed more conventional methods.
- This method provides groundwork for detailed multimodal hippocampal mapping across subjects and datasets in the future.

**Data availability:** The methodological advancements described here are made easily accessible in the latest version of open source software HippUnfold^1^. Code used in the development and testing of these methods, as well as preprocessed images, manual segmentations, and results, are openly available^2^.

## Introduction

The hippocampus is part of the archicortex that, like the neocortex, can be further parcellated into subfields according to its cytoarchitecture (Ding & Van Hoesen, 2015; Duvernoy et al., 2013; Palomero-Gallagher et al., 2020). The study of hippocampal subfields is promising for both basic science, since their microcircuits are thought to be fundamental to memory processes (Milner et al., 1968; O’Keefe & Nadel, 1978; Riphagen et al., 2020), and for the pathogenesis of several brain disorders given their vulnerabilities to many conditions, notably epilepsy (Bernhardt et al., 2015, 2016; Blumcke et al., 2013; Thom, 2014), Alzheimer’s disease (Braak & Del Tredici, 2014; Flores et al., 2015), and Schizophrenia (Haukvik et al., 2018; Roeske et al., 2020). However, subfield parcellation is challenging. This relates to variability across segmentation protocols both at the level of histology and imaging (Wisse et al., 2017; Yushkevich, Amaral, et al., 2015). Moreover, the hippocampus has a complex shape that varies between individuals (Chang et al., 2018; DeKraker et al., 2021; Ding & Van Hoesen, 2015; Palomero-Gallagher et al., 2020). This topic has received widespread attention, leading to the development of an international harmonization effort that focuses on extracting geometric regularities from reference histology slices that can be applied to MRI, mostly using coronal slices (Olsen et al., 2019; Wisse et al., 2017; Yushkevich, Amaral, et al., 2015). Building on this discussion, we assert that the most basic geometric consistency of the hippocampus is a 3D folded surface and so subfield parcellation schemes should be applied using surface-based registration, similar to state-of-the-art neocortical parcellation.

Surface-based registration, either in the hippocampus or neocortex, aims to account for inter-individual differences in folding or gyrification patterns during registration (DeKraker et al., 2021; Fischl et al., 2008; Im et al., 2008; Lyttelton et al., 2007; MacDonald et al., 2000; Van Essen et al., 2000); Robbins, 2003). Briefly, this consists of first generating a 3D model of the structure of interest and representing it as a surface, typically a mid-thickness surface. In the neocortex, this surface can be “inflated” or remapped, *e*.*g*. for viewing on the surface of a sphere. Registration can then be performed between two spheres based on some feature map by rotating until the maximum overlap achieved (Kim et al., 2015; Klein et al., 2010; Lyttelton et al., 2007). Posing registration problems on a sphere helps account for inter-individual variability in gyral and sulcal patterning, which can vary drastically between individuals (Bartley et al., 1997; Goualher et al., 1999; Régis et al., 2005). For example, conventional 3D volumetric registration may take one gyrus and stretch it across two gyri from another individual, especially in areas where the number or shape of gyri varies between individuals. In surface-based registration, homologous points are not constrained to fall in similar absolute positions but rather similar topological positions (*e*.*g*., one gyral peak could be homologous to a point halfway down the depth of an adjacent sulcus from another individual). Since the major gyri and sulci of the brain are typically invariant across individuals (Le Guen et al., 2018), gyral and sulcal patterning can be used as one feature to inform registration, often after smoothing to remove smaller secondary or tertiary gyri and sulci which tend to be more variable (Tardif et al., 2015), while other features more indicative of cortical architecture, such as thickness or intracortical myelin, can be used as well (Glasser et al., 2016; Lyttelton et al., 2007; Van Essen et al., 2012).

Here, we present such a surface-based registration method specifically for hippocampal surfaces. Rather than inflation to a sphere, we rely on previous work which maps the hippocampus to a flat rectangle to preserve its topology (DeKraker et al., 2022). This also allows for the use of existing 2D image-based registration tools without reformulation for use on a surface mesh but, in effect, the same advantages and constraints as typical surface-based registration are preserved. Evaluation of this method is performed using ground-truth (*i*.*e*., histologically-derived) subfield segmentations from seven samples that were sliced, imaged microscopically, and then digitally reconstructed into a 3D block with various histology contrasts. We benchmark this new method against unfolding alone, which provides some intrinsic alignment between subjects (DeKraker et al., 2018) but which we believe can be further improved with the present methods, and against more conventional 3D volumetric registration approaches. The method has been openly published at https://github.com/khanlab/hippunfold/tree/unfold_reg.

## Materials and Methods

### Data

Seven 3D reconstructed hippocampal histology samples were examined in this study from three different datasets including four donor brains:

1. BigBrain: 2 hemispheres (donor 1), Merker stain (Merker, 1983), downsampled from 20 to 40μm isotropic voxel resolution (Amunts et al., 2013),
2. 3D Polarized Light Imaging (3D-PLI): 1 hemisphere (donor 2), 48x48x60μm resolution, contrast driven by birefringence properties of myelin sheaths surrounding axons (Axer et al., 2011), and
3. AHEAD: 4 hemispheres (donors 3 and 4), blockface imaging and multiple stains, 150x150x200μm resolution (Alkemade et al., 2022).

Details of these dataset acquisitions and processing can be found in their respective references. The 3D-PLI sample was examined here only in terms of transmittance rather than directional data. The transmittance has already been demonstrated in a few brain sections to provide valuable information to segment hippocampal subregions (Zeineh et al., 2017).

### Manual segmentation

HippUnfold (DeKraker et al., 2022) achieves automatic segmentation and unfolding of *in-vivo* hippocampal MRI data, typically T1w or T2w images, which do not resemble the contrasts seen in the present histology datasets and thus manual segmentation had to be used. This unfolding method requires a detailed hippocampal gray matter mask, as well as labeling of termini at the anterior, posterior, and medial hippocampal edges, dentate gyrus (DG), and laminar strata radiatum and lacunosum-moleculare (SRLM). Here, we consider SRLM to be a “mixed” label since it can include components of the subicular complex, CA fields, DG, as well as blood vessels and CSF within the hippocampal sulcus. Thus, it is used to differentiate the upper and lower surfaces of the remaining hippocampal cortex, contiguous with the “pial” and “white” surfaces of the neocortex, respectively, and SRLM volume is excluded from further analyses.

In BigBrain, these labels were available from previous work (DeKraker et al., 2020). Rater J.D. performed manual segmentation in all other samples using ITK-SNAP (Yushkevich et al., 2006). In the 3D-PLI sample, the absolute anterior and posterior of the hippocampus were cut off during tissue preparation. Thus, the required labels were extrapolated manually over the missing regions in order to recover a fully 3D hippocampal shape that is amenable to unfolding (see **Figure 1A**). In the AHEAD brain samples, manual segmentation was performed on blockface images.

**Figure 1.**
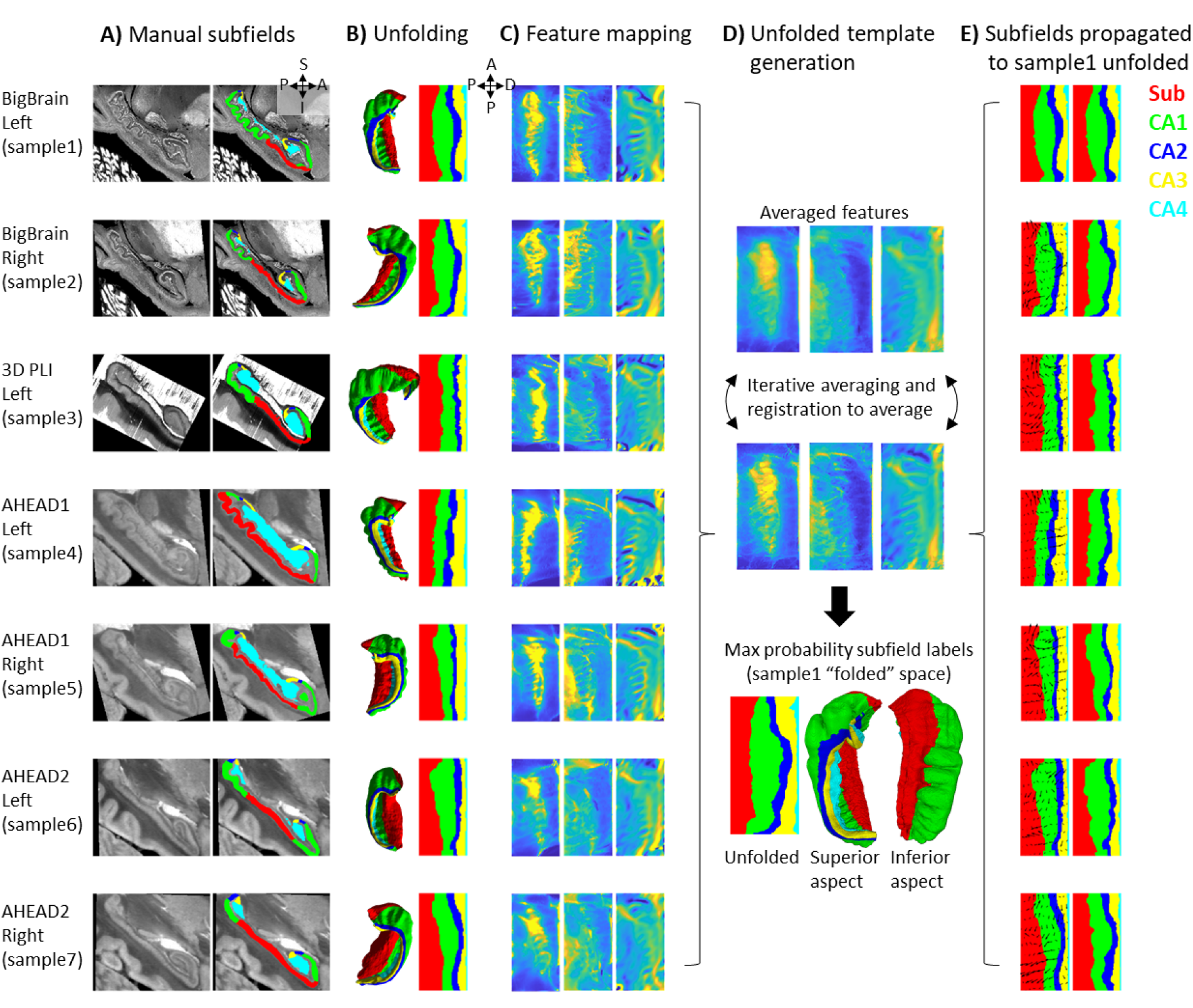
Surface-based subfield alignment pipeline. **A)** Each 3D dataset manually parcellated into subfields. **B)** Subfields mapped to a common unfolded space using HippUnfold. **C)** Morphometric features mapped to unfolded space (from left to right: gyrification, thickness, curvature). **D)** Multimodal features iteratively aligned in unfolded space using 2D registration. **E)** Subfield labels propagated through unfolded 2D registrations to sample1. Closeups of the segmentations for the individual datasets can be found in **Supp. Figure 2**.

For all datasets, hippocampal gray matter was subsequently manually parcellated by rater J.D. into subfield labels: Subicular complex (Sub) and Cornu Ammonis (CA) fields 1-4 based on the available histological features and according to the criteria outlined by (Ding & Van Hoesen, 2015; Duvernoy et al., 2013; Palomero-Gallagher et al., 2020). Since these sources differ slightly in their boundary criteria, and no prior reference perfectly matches the present samples, subjective judgment was used to draw boundaries after considering all three prior works. The “prosubiculum” label used by Ding & Van Hoesen and Palomero-Gallagher et al. was included as part of the subicular complex. See Supplementary Materials 2: ground-truth segmentation for more details. The DG was also labeled but was grouped together with CA4 since HippUnfold’s current method for unfolding of the DG relies on heuristics in MRI that would not be appropriate in this work (*i*.*e*., it uses a template prior to estimate DG topology at the cost of smoothing labels). These manual labels were defined based upon cytoarchitectonic features at the highest level of resolution available and were deemed “ground-truth” subfield definitions. It is important to note that BigBrain is stained for cell bodies, while 3D-PLI transmittance contrast is driven by cell bodies and nerve fibers (both introducing light extinction effects; Menzel et al., 2020) and thus contains very different microstructural information. Thus, both cell body and fiber distribution patterns were consulted during subfield definition. In the AHEAD dataset, multiple imaging modalities were available, albeit with imperfect registration to the blockface images, larger interslice gaps, some missing data, and limited resolution. These additional contrasts were overlaid over blockface images (where available and appropriate due to the above limitations) to better inform subfield segmentation.

### Unfolded registration

HippUnfold (DeKraker et al., 2022) was used to map each dataset to a standardized unfolded space (see **Figure 1B**). This unfolded space consists of a triangulated mesh with uneven face sizes, so as to preserve a constant spacing between points in the folded hippocampus. The current work, however, defined this tessellation as a regular mesh grid in unfolded space consisting of 256×128 points across the anterior-posterior (A-P) and proximal-distal (P-D) (relative to the neocortex) axes of the unfolded hippocampus, respectively. This regular grid in unfolded space means that surface points (henceforth, vertices) can effectively be treated as pixels of a flat 2D image without the need to interpolate missing pixel values. However, it should also be noted that, as a consequence, native or “folded” images are sampled more densely in some areas than others, particularly in the anterior and posterior extremes, which can lead to noisier unfolded data in these regions.

HippUnfold also calculates morphological features, namely thickness, gyrification index, and curvature in each subject’s native space (**Figure 1C**). These features are desirable for inter-subject registration since *i)* they are associated with subfield boundaries (DeKraker et al., 2018, 2022; Yushkevich, Pluta, et al., 2015), *ii)* they don’t require cytoarchitectonic information to measure, and *iii)* they are agnostic to imaging contrast differences. Registration performed in unfolded hippocampal space is analogous to registration of neocortical surfaces that have been inflated to a sphere, since both methods preserve topology rather than absolute position. However, one difference is that registration on a sphere allows one point to “wrap” around the meridian of a sphere whereas in a rectangular unfolded space, the proximal edge (*i*.*e*., closest to the neocortex) does not “wrap” to the distal (*i*.*e*., closest to the dentate gyrus) edge of the P-D axis, and the same applies to the anterior-posterior, A-P, edges. This is in agreement with the true geometry of the hippocampus though, which has the topology of a rectangle (*i*.*e*., four true edge termini) rather than a sphere (*i*.*e*., zero true termini).

The 2D registration was performed using all three of the above morphological features with equal weighting, using ANTs multicontrast SyN deformable registration (Avants et al., 2011) (**Figure 1D**). Rather than registering all samples’ feature maps to one sample, we instead used an iterative template building method (Avants et al., 2010), which first averages images, registers each image to the average, and then repeats the registration to the newly generated average. This process is repeated 4 times, with each iteration sharpening the averaged template and improving registration precision. We concatenated the transforms from each sample to the template with the inverse transform from the template to sample1 (BigBrain left hemisphere) and applied it to subfield labels to evaluate their overlap with that sample’s ground-truth subfield definitions (**Figure 1E**). Sample1 was chosen because it had the highest resolution and, therefore, provided the greatest cytoarchitectonic detail for identification of subfield boundaries, while also being the more common hemisphere in this study (*i*.*e*., 4 left; 3 right). In principle, any sample could have been chosen, and one had to be chosen to test overlap in any one native (or folded) space.

### Control condition: volumetric registration

The proposed pipeline was compared to a conventional 3D volumetric registration approach: ANTs template generation (Avants et al., 2011) under ideal tissue contrast conditions (*i*.*e*., based on binarized gray matter labels). This is outlined in **Figure 2** and detailed below. First, all hippocampal gray matter labels were binarized, right hippocampi were flipped, and binary masks were rigidly registered to sample1 using Greedy’s moment-based initialization (Yushkevich et al., 2016) (2 moments), which can handle images initially in different spaces relative to the origin (which was the case in some samples here) (**Figure 2B**). Images were then resampled to 100μm isovoxel resolution to reduce compute time and memory requirements, which were prohibitively high at native resolution. ANTs template generation was then used as before (**Figure 2C**), with the following differences: registrations were all 3D and unimodal.

**Figure 2.**
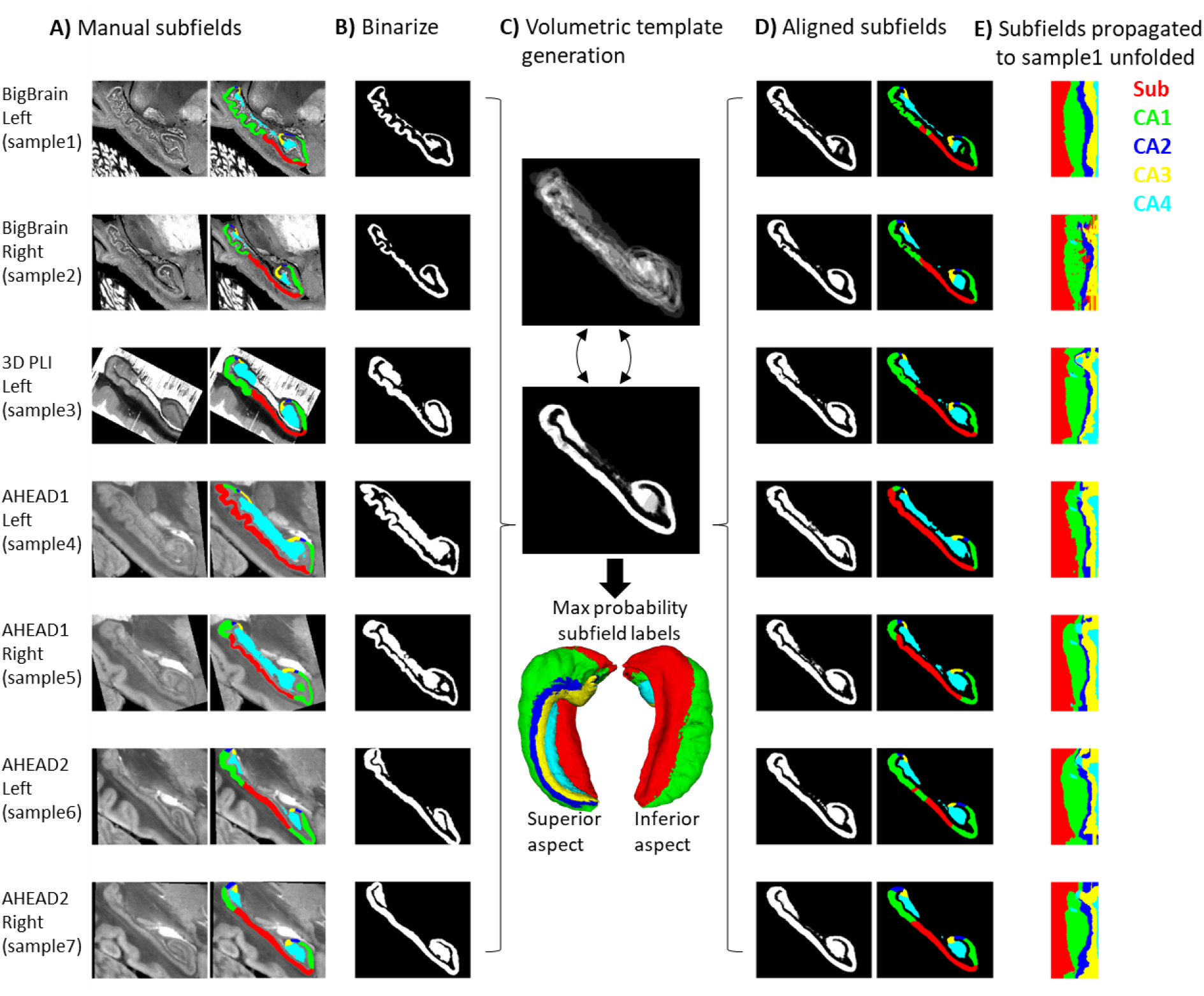
Control condition using volumetric registration. **A)** Subfield segmentation as in **Figure 1A. B)** Binarized and rigidly aligned hippocampal gray matter masks (left hemispheres flipped). **C)** Iterative alignment using ANTs template building in 3D. **D)** Each sample’s subfields propagated to template space. **E)** Each sample’s subfields propagated to sample1 unfolded space.

Using binarized images likely overestimates a fully automated registration method for subfield parcellation, since it presents idealized tissue contrast conditions (*i*.*e*., perfect contrast between grey and white matter). This has been chosen as a practical and robust approach, avoiding local minima commonly encountered by image-intensity based registration procedures in the mesiotemporal region (Qiu & Miller, 2008). In principle, cross-modal registration can be performed using metrics like mutual information or cross-correlation, but these metrics still do not fully compensate for differences in contrasts, luminance, or intensity levels, and may have many local minima solutions making it less tractable. Thus, the control condition used here could be thought of as a best-case volumetric registration.

Registrations from each sample to the template and the inverse transform from the template to sample1 were concatenated and applied to subfield labels (**Figure 2D**). For easier comparison with **Figure 1**, each sample’s subfields were sampled along sample1 mid-thickness surface and projected to unfolded space (**Figure 2E**). In some cases, the mid-thickness surface fell outside of the propagated subfield labels and returned a background value of 0. The missing values were imputed by nearest-neighbour interpolation in unfolded space, providing additional correction for misregistration.

### Evaluation metrics

The Dice overlap metric (Dice, 1945), which can also be considered an overlap fraction ranging from 0-1, was calculated for all subjects’ subfields registered to sample1. This was repeated in 2D unfolded and in 3D native spaces, since some parts of unfolded space expanded or contracted more than others when projected to native space, which can over- or under-emphasize subfield differences. For example, sample3 was unfolded and then registered to the unfolded average, making up two transformations. These were then concatenated with the inverse transformation of unfolded sample1 to the same unfolded average, and the inverse transformation of native sample1 to unfolded space. This concatenated transformation was used to project labels from sample3 native space directly to sample1 native space, which should ideally lead to near-perfect subfield alignment in sample1 native space. Dice overlap between sample1 and sample3 registered to sample1 was then calculated in sample1 native space. A secondary metric, border distances, was also calculated in sample1 native space. This was calculated by computing distance from a given border in sample1 to all voxels, followed by concatenating these distances at the location of each propagated sample’s corresponding subfield borders, providing a minimum direct 3D distance between borders in real-world units. These distances were calculated from all subfield borders (*i*.*e*., Sub-CA1, CA1-CA2, CA2-CA3, and CA3-CA4).

In addition to registration in 2D unfolded space and 3D native space, overlap metrics were calculated for unfolding without registration. That is, borders from each sample were projected to unfolded space and projected directly to sample1 space, which replicated the current subfield segmentation behaviour in HippUnfold (v1.2.0).

To evaluate the contribution of each morphometric feature or combination of features (thickness, gyrification, and curvature) to registration, unfolded space registrations were also performed using all combinations of these features. These were then evaluated using Dice scores in sample1 native space.

## Results

### Qualitative alignment

**Figure 1A** shows equivalent sagittal slices from each sample after rigid alignment, in which considerable subfield variability can be seen between samples. This is due in part to the out-of-plane issue discussed elsewhere (DeKraker et al., 2021). Viewing a fully 3D model (**Figure 1B**) can help in identifying differences between samples due to true morphological variability rather than field-of-view differences. Projecting these labels to unfolded space (**Figure 1B**) preserved sample-specific differences in the size of each subfield but removed variability due to different folding and gyrification configurations between samples, which already brought subfields into close alignment and was the current basis for subfield segmentation in HippUnfold.

Following registration in unfolded space (**Figure 1E**), subfields are even more closely aligned (for example samples 3 and 5 are no longer dominated by the Sub label). In addition, there are no topological breaks or reordering of subfields in **Figure 1**. In **Figure 2E**, following conventional volumetric alignment, there are cases where CA1 shows isolated islands (*e*.*g*. sample2), and where CA1 borders CA3 directly instead of first passing through CA2 (*e*.*g*. sample3). This does not match the original subfield segmentations from any sample or the literature which states that subfields should be contiguous and consistently ordered (Duvernoy et al., 2013). These issues arise in 3D registration due to breaks in topology: for example, hippocampal gray matter may become stretched across the SRLM and vestigial hippocampal sulcus, or across adjacent gyri. When subfield labels are propagated across a sulcus, they can become discontinuous with respect to the mid-thickness surface topology of a given sample. This type of 2D topology is conserved in a surface-based or unfolded hippocampal space registration.

### Quantitative alignment

Better Dice overlap on average and for every individual subfield was observed using unfolded registration over unfolding and refolding into a different sample’s folded configuration alone (**Figure 3B**). Both methods also outperformed the control condition using conventional ANTs 3D volumetric registration. These results were tested using one-tailed, paired-samples t-tests, pairing subfields and subjects, which revealed significant differences between each method and in both unfolded and native spaces. The same pattern was also observed when evaluated according to nearest corresponding subfield border distances in native space (**Figure 3C**). One example of each of these registration methods (sample3 to sample1) is shown in **Figure 1A**.

**Figure 3.**
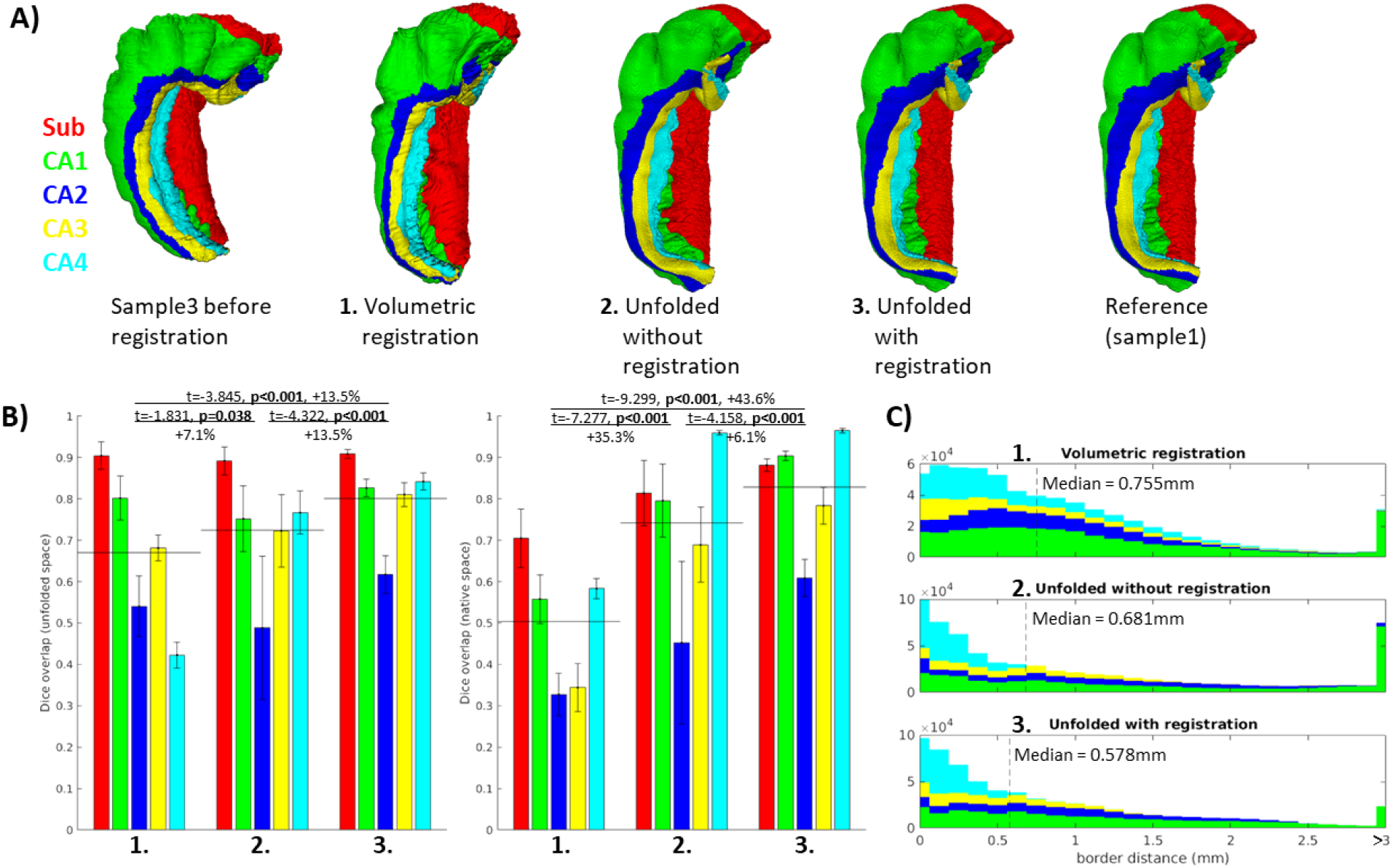
Evaluation of aligned subfield definitions. **A)** Qualitative example of subfields from the third sample projected to the first sample’s native “folded” space using conventional 3D volumetric alignment, unfolding to account for inter-individual differences in folding shape, and unfolding followed by registration in unfolded space. **B)** Dice overlap achieved. Each measure was calculated in unfolded space (left) and again in the first sample’s (BigBrain left hemisphere) native folded space (right). Black lines indicate the mean across all subfields. **C)** Distances between all aligned subfield borders using the three methods described above. Dashed lines indicate the median distance.

### Contribution of unfolded morphological features

By using different combinations of unfolded features, we could determine which is most informative about unfolded registration and subfield boundary alignment (**Figure 4**). In this regard, curvature was the most informative individual feature while thickness and curvature was the most informative combination of two features, despite the fact that thickness was the least informative individual feature (similar to no unfolded registration). Therefore, while each is informative, they may contain overlapping information and their combination isn’t as complementary as curvature and thickness together. Regardless, combining all three features still showed the best performance. Future work should explore the use of additional features, such as intracortical myelin or laminar distributions of neurons, to inform registration. This was not examined here since it was not available in all datasets owing to differing contrasts.

**Figure 4.**
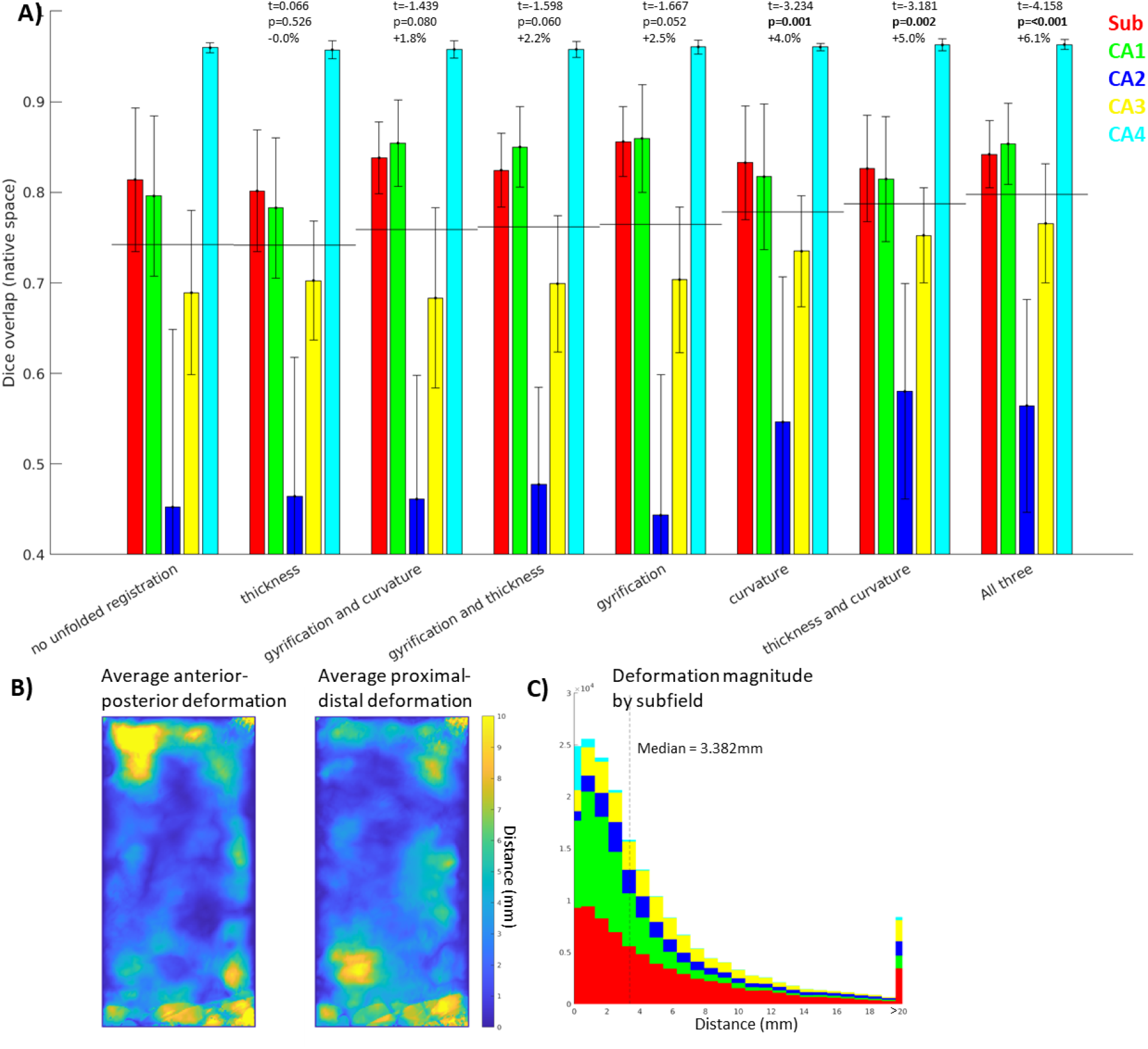
Contribution of each morphological feature to unfolded registration performance. A) Unfolded space registration was repeated for all combinations of unfolded morphological features and evaluated by Dice overlap in native space. Combinations are ordered by their Dice scores averaged across the 5 subfields. p-values are relative to no unfolded registration, using one-tailed paired-samples t-tests as above. B,C) Evaluation of which hippocampal vertices (B) and subfields (C) were most deformed in unfolded registration.

**Figure 4B and C** show the extent of deformations in unfolded space following ANTs registration with default parameters. The median deformation magnitude across all vertices was 3.382mm, but more constrained or liberal deformations could be achieved by adjusting the ANTs elasticity and fluidity parameters (proportional to the resulting deformation field’s maximum and smoothness, respectively). More liberal deformations could lead to more precise subfield alignment, however, this also runs the risk of falling into local minima during optimization, or of distorting tissue beyond a reasonable distance. Thus, default parameters were used to maintain robustness at the cost of potential gains in precision.

## Discussion

The current work presented a novel surface-based registration of hippocampal cortex between histology samples, and our evaluations demonstrated that it outperformed even an idealized version of conventional volumetric registration. This enabled mapping of multiple features at a sub-millimetric scale that would otherwise be impossible. For example, maps of hippocampal cyto- or myeloarchitectonic features can be constructed from multiple samples using different stains or imaging methods that would otherwise preclude one another due to tissue destruction (*e*.*g*., once stained, a slice cannot be easily imaged with other stains). This can be useful both in studying hippocampal subfields and in subfield-agnostic mapping and other data driven methods (*e*.*g*., Borne et al., 2023; Paquola et al., 2020; Patel et al., 2020; Przeździk et al., 2019; Vogel et al., 2020; Vos de Wael et al., 2018). We hope that this work will provide an avenue towards mapping of hippocampal data across many modalities, scales, and different fields in future work.

Surface-based hippocampal registration can be used in subfield parcellation in MRI, histology, or other imaging methods by registration to the unfolded maximum probability subfield atlas provided here (Figure 1D). Note that the current work differs from other subfield segmentation protocols, even those which employ surfaces and unfolding, in that our method constrains registration (and therefore subfield segmentation) topologically by the unique folded shape of a given hippocampus. Other methods generally first employ subfield segmentation (either manually or using conventional volumetric registration) and then reduce volumes to surfaces or employ other flat mapping techniques of the hippocampus (Caldairou et al., 2016; Ekstrom et al., 2009; Pipitone et al., 2014; Yushkevich et al., 2016; Yushkevich, Pluta, et al., 2015; Zeineh et al., 2000, 2001). For example, SurfPatch (Caldairou et al., 2016) computes volumetric registration and then propagates surfaces rather than labelmaps, which avoids discretization errors. However, this method still does not guarantee correct topology across different hippocampal folding patterns (see discussion below on multi-atlas registration), and has not been demonstrated at the level of detail examined here.

Other surface-based registration methods such as spherical harmonics (SPHARM) (Brechbühler et al., 1995; Gerig et al., 2001) have been employed to find homologous vertices between irregular shapes including hippocampal samples (Styner et al., 2004). In principle, this is a similar pose of the registration problem as the methods employed here. However, SPHARM requires a spherical topology, which in (Styner et al., 2004) is mapped to the outer boundaries of the hippocampus rather than a midthickness surface, and so this method does not fully leverage the geometric topology constraints of the hippocampus. There has been an adaption of the SPHARM-PDM model to hippocampi, in which the spherical parameterization of the outer hull was propagated along a Laplacian field to the hippocampal midthickness surface (Kim et al., 2014), and this approach has since then been used and validated in the study of hippocampal organization in both health and disease (Bernhardt et al., 2016; Vos de Wael et al., 2018). Other vertex-wise or even point-cloud registration methods could be employed for hippocampal midthickness surfaces in future work. One final example is the recent FastSurfer implementation of Laplace Eigenfunctions for neocortical surface registration, which involves registration to a sphere (Henschel et al., 2020). This method does not require an inherently spherical topology and the only major conceptual difference between it and the present work is that we hold hippocampal termini or endpoints fixed, for additional regularization, whereas FastSurfer derives them from the surface mesh itself.

Ravikumar et al. (2021) recently performed flat mapping of the medial temporal lobe neocortex using a Laplace coordinate system as employed here, and showed sharpening of group-averaged images following deformable registration in unfolded space. This indirectly suggests better intersubject alignment. We perform a similar group-averaged sharpening analysis in Supplementary Materials 1: in-vivo demonstration. Though the gains in image sharpness observed here were modest, we note that current MRI resolution and automated segmentation methods allow for only imperfect hippocampal feature measures. We thus expect unfolded registration to grow in importance as MRI and segmentation methods continue to advance.

The feature most strongly driving surface-based registration in the present study was curvature. Many neocortical surface-based registration methods focus on gyral and sulcal patterning at various levels (e.g. strong alignment of primary sulci, with weaker weighting on secondary and tertiary sulci) (Miller et al., 2021). In the present study, hippocampal gyri are variable between samples and so could perhaps be thought of as similar to tertiary neocortical gyri, and this may help explain why gyrification was not the primary driving feature in aligning hippocampal subfields. The methods used to quantify gyrification are often related to curvature, but differ across studies. In the hippocampus, unlike in the neocortex, the mouth of sulci are wide and so sulcal depth, which is often used in defining neocortical gyrification index, is not straightforward to measure. Using HippUnfold, gyrification is defined by the extent of tissue distortion between folded and unfolded space, and individual gyri/sulci are hard to resolve in unfolded gyrification maps, but are readily visible in curvature maps. Thus, hippocampal curvature may be more informative simply due to higher measurement precision. Future work could also employ measures like quantitative T1 relaxometry or other proxies of intracortical myelin content (*e*.*g*. Tardif et al., 2015; Glasser et al., 2016; Paquola et al. 2018) for hippocampal alignment, but this is not possible in cross-modal work as in the various histology stains examined here.

One limitation of the evaluation performed on our surface-based registration is that our control condition using 3D volumetric registration did not employ a multi-atlas as in some other popular methods, including those discussed above (Caldairou et al., 2016; Yushkevich et al., 2016; Yushkevich, Pluta, et al., 2015). A multi-atlas applies registration of several references to a target image and then combines propagated labels from the references to the target (*e*.*g*. via maximum probability). We partially compensated for this issue by using an idealized volumetric registration with detailed binarized hippocampal gray matter masks rather than images, which generally contain more noise, blurring, and surrounding structures that are not necessarily informative about hippocampal shape. 3D registrations at the current resolution also have costly compute requirements that scale with resolution and the number of samples in the multi-atlas. In addition, a multi-atlas is not guaranteed to contain a sample with a similar folding configuration as the target sample and, even if it does, the combination of multiple registered samples may lead to errors or over-smoothing. By contrast, HippUnfold can be moderately compute intensive at high resolution, but it only needs to be performed once and then registration in unfolded space has trivially light compute requirements, even when using multiple contrasts as performed here. Nevertheless, we hope to eventually see the development of multi-atlas volumetric registration at a microscale, as well as work performing a systematic comparison with surface-based registration.

## Conclusions

We formulated a registration performed in a standardized “unfolded” hippocampal space, and showed that this method consistently improved inter-individual alignment with respect to subfields. This method is topologically-constrained and driven by contrast-agnostic feature maps, meaning that it can be performed across image modalities regardless of whether cytoarchitectonic features are directly accessible or not. Overall, this work constitutes a state-of-the-art registration method between hippocampi at a scale approaching the micron level.

## Acknowledgements

This work was supported by the Helmholtz International BigBrain Analytics and Learning Laboratory (HIBALL), funded by Healthy Brains, Healthy Lives (HBHL) and the Helmholtz Association. JD was furthermore supported by the National Science and Engineering Research Council of Canada (NSERC). BCB acknowledges research support from NSERC (Discovery-1304413), Canadian Institutes for Health Research (CIHR FDN-154298, PJT-174995), SickKids Foundation (NI17-039), BrainCanada, HBHL, and the Tier-2 Canada Research Chairs program. 3D-PLI related work was supported by the European Union’s Horizon 2020 Framework Programme for Research and Innovation (grant no. 945539: “Human Brain Project” SGA3 and by the computing time granted through JARA on the supercomputer JURECA at Forschungszentrum Jülich.

## Footnotes

1 https://github.com/khanlab/hippunfold/pull/228

2 https://zenodo.org/record/7757416

