## Supplementary Materials 1 for "Evaluation of surface-based hippocampal registration using ground-truth subfield definitions"

### Supplementary Materials 1: *in-vivo* MRI demonstration

In order to determine whether the gains from unfolded registration introduced in the current histology work translate to *in-vivo* MRI, we examined a set of 10 healthy participants scanned at 7 Tesla magnetic field strength. Each subject’s T1w image was run through the automated HippUnfold pipeline (DeKraker et al., 2022) with and without the inclusion of unfolded space registration to the group-averaged template derived in the present study. We reasoned that including unfolded registration should lead to better alignment and therefore sharper images after averaging across subjects and hemispheres, as demonstrated, for example by Ravikumar et al. (2021). Here, we quantify sharpness as the mean gradient magnitude (MGM) of the averaged image. Morphological features used in this unfolded space alignment, namely, gyrification, thickness, and curvature, showed a 10.9%, 0.6%, and 9.1% increase in MGM, respectively.

We also measured quantitative T1 relaxation times (qT1) along hippocampal midthickness surfaces. This generally showed differences across the proximal-distal, or subfield-related, axis of the hippocampus, with highest values being found in the CA1 region indicating a relatively low intracortical myelin content compared to the other subfields. Group-averaged qT1 MGM also increased, by 2.1%, with the inclusion of registration in unfolded space. This feature was not used to inform unfolded registration but still showed a small increase in sharpening with group averaging, which reinforces the validity of morphometry as a basis for registration across many domains.

It should be noted that the gains from unfolded registration are likely curtailed in *in-vivo* MRI compared to in histology due to decreased resolution and therefore reduced sensitivity of morphological measures. Even at 7 Tesla MRI field strength, it was noted that considerably less detail (namely, less small folds or gyrifications within the hippocampus) were visible both in the raw images and in the automatically generated image segmentations and surfaces. This leads to overall lower gyrification and curvature, and increased thickness measures. We thus note that the gains in intersubject alignment demonstrated here may not be as clear in more commonly used 3 Tesla MRI, but we hope that as acquisition methods continue to advance, the alignment methods demonstrated here will only grow in importance.

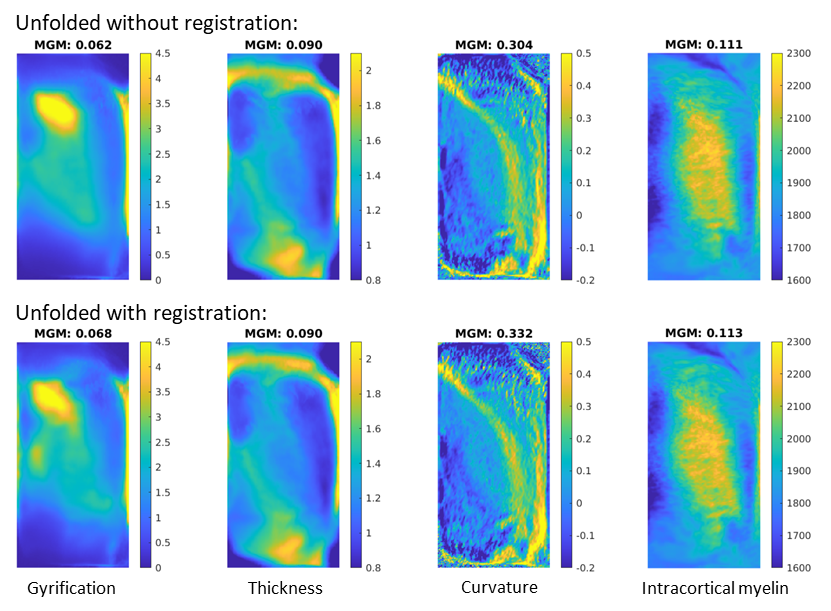

Figure 1. Image sharpness when averaging *in-vivo* MRI subjects’ hippocampal features with and without unfolded space registration. (MGM = mean gradient magnitude of the group-averaged image).

#### Methods:

All participants were recruited between May 2022 and April 2023 at the Montreal Neurological Institute (MNI). Healthy controls met the following inclusion criteria: (1) age between 18-65 years; (2) no neurological or psychiatric illness; (3) no MRI contraindication; (4) no drug/alcohol abuse problem; and (5) no history of brain injury and surgery. The Research Ethics Board (REB) at McGill University approved this study. 10 participants were examined here (mean age 26.7 years, S.D. 4.36).

Scans were acquired using a 7 Tesla Siemens MRI at the McConnell Brain Imaging Centre of the Montreal Neurological Institute. Each participant underwent multiple types of scans, including T1-weighted (T1w) structural MRI, diffusion-weighted imaging (DWI), resting-state functional MRI (rs-fMRI), and quantitative T1 (qT1) mapping. For the purposes of the present study, we will discuss only qT1 imaging.

qT1 relaxometry data was acquired using a 3D-MP2RAGE sequence, with the following parameters: 0.5mm isotropic voxels, 320 sagittal slices, TR=5170 ms, TE=2.44 ms, TI1=1000 ms, TI2=3200 ms, flip angle=4°, flip angle 2=4°, iPAT=3, bandwidth=210 Hz/px, echo spacing=7.8 ms, and partial Fourier=6/8. Both inversion images were combined for qT1 mapping to minimize sensitivity to B1 inhomogeneities and optimize intra- and inter-subject reliability (Haast et al., 2016; Marques et al., 2010).

HippUnfold was run on raw qT1 images. The inclusion of unfolded space registration was performed using the latest HippUnfold software release (v1.3.0, described fully at https://github.com/khanlab/hippunfold/pull/228) which includes the methods and reference histology data described in the current study.

Outliers (3 S.D. from the median) were clipped for each feature before averaging. Each group-averaged feature (gyrification, thickness, curvature, and qT1) was z-scored before calculating the image gradient and mean gradient magnitude (MGM). This makes the MGM ranges more similar for easier comparison across the different features.
