## Supplementary Materials 2 for "Evaluation of surface-based hippocampal registration using ground-truth subfield definitions"

### Supplementary Materials 2: ground-truth segmentation

As discussed in the Introduction section, the definition of hippocampal subfields in MRI differs systematically between labs (see Yushkevich, Amaral, et al., 2015), and this is also the case in “ground-truth” histological subfield definitions (Olsen et al., 2019). To our knowledge these differences have not been quantified systematically, but (Olsen et al., 2019) demonstrate considerable differences between raters even with a coronal slice where no out-of-plane sampling issues are present. The evaluation of unfolded hippocampal subfield registration detailed in the present study does not rely on harmonized subfield definitions, but rather on internal (or intra-rater) consistency and were thus all performed by J.D. However, we nevertheless sought to determine whether these definitions matched those of other neuroanatomists in the field in order to determine the value of the current ground-truth labelmaps as a reference material for future work. We have made labelmaps for all samples in native and unfolded spaces and the maximum probability unfolded subfield labels available online (see Data Availability Statement).

Expert histologist O.K. performed manual subfield annotations in a subset of 66 coronal slices from the BigBrain left and right hemispheres. O.K.’s annotations included labels “parasubiculum”, “subiculum proper”, “presubiculum”, and “prosubiculum” (Palomero-Gallagher et al., 2020); Amunts et al., 2021; doi: 10.25493/X4S6-E64) which were all combined into a single label to match the “subicular complex” label employed by rater J.D. Additionally, to focus on borders between subfields rather than differences in gray matter definition, the same mask of the entire hippocampus was applied to both labelmaps. This mask was defined as the voxels for which both J.D. and O.K. had labeled any subfield. Dice overlap was calculated across all labeled coronal slices as well as in unfolded space (**supp. Figure 1**), and for a full visualization of both raters’ labels on all slices after matching gray matter bounds, see **supp. Figure 2**.

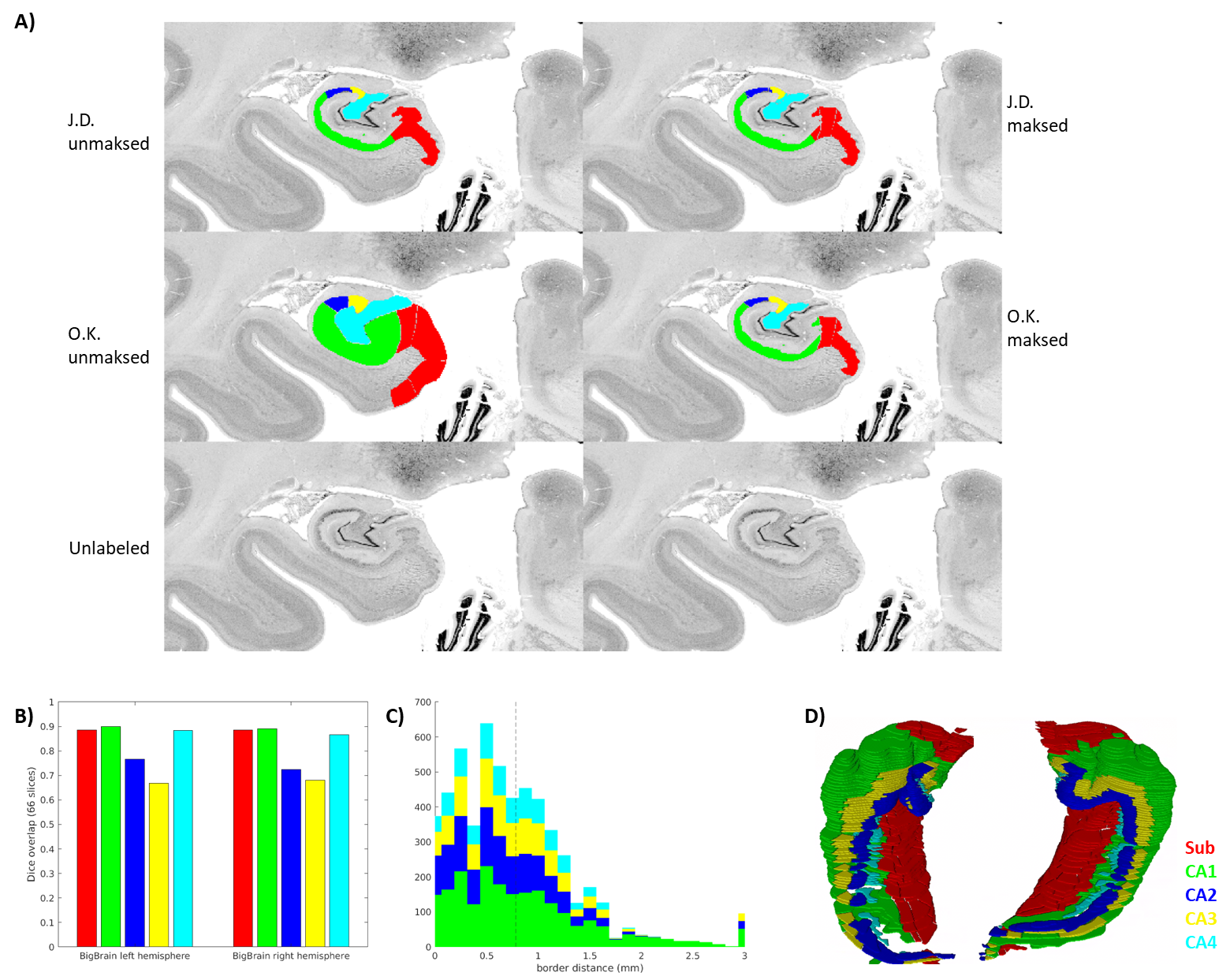

**Supp. Figure 1**. Inter-rater ground-truth subfield segmentation in the left and right BigBrain hippocampus (sample1 and sample2). **A)** Grey matter mask (defined as any unlabelled voxel from either rater) applied to one slice. **B)** and **C)** show inter-rater Dice and border distances, respectively, as in **Figure 3**. **D)** Stacked coronal slices from rater O.K.

The median distance between ground-truth raters was 0.781μm, and Dice scores in coronal slices were in the range considered very good (0.8-0.9) for larger subfields Sub, CA1, and CA4, and moderate (0.6-0.8) for smaller subfields CA2 and CA3. It is interesting to note that in CA3 and CA4, the proposed unfolded registration method actually outperformed ground-truth label reliability. Ground-truth reliability should represent the ceiling for automated registration performance. Two factors, thus, likely led to systematically higher Dice scores and lower border distances in the unfolded registration evaluation: 1) all labeling was performed by rater J.D. who likely had different but consistent criteria for identifying borders, and 2) unfolded registrations are bound by the same distal “endpoint”. This is defined within HippUnfold as being the DG or the true topological “terminus” of the archicortex (making up also a portion of the true terminus of the cortex altogether). This terminus point can, thus, act as an anchor for homologous points between samples, which is reflected most strongly in the evaluation in overlap of the subfields closest to that point: CA4 and CA3. HippUnfold leverages this topological endpoint in determining homology between samples, but it is important to note that this method relies on consistent identification of that endpoint (which is relatively clear in histology but can be challenging in MRI, see (DeKraker et al., 2022) for discussion).

**Supp. Figure 2**. Side-by-side comparison between raters (O.K. top; J.D. middle; unlabelled bottom) in a subset of 54 slices where both the left and right hemispheres were fully labeled. Slices are ordered from anterior-to-posterior (cont’d 9 pages).

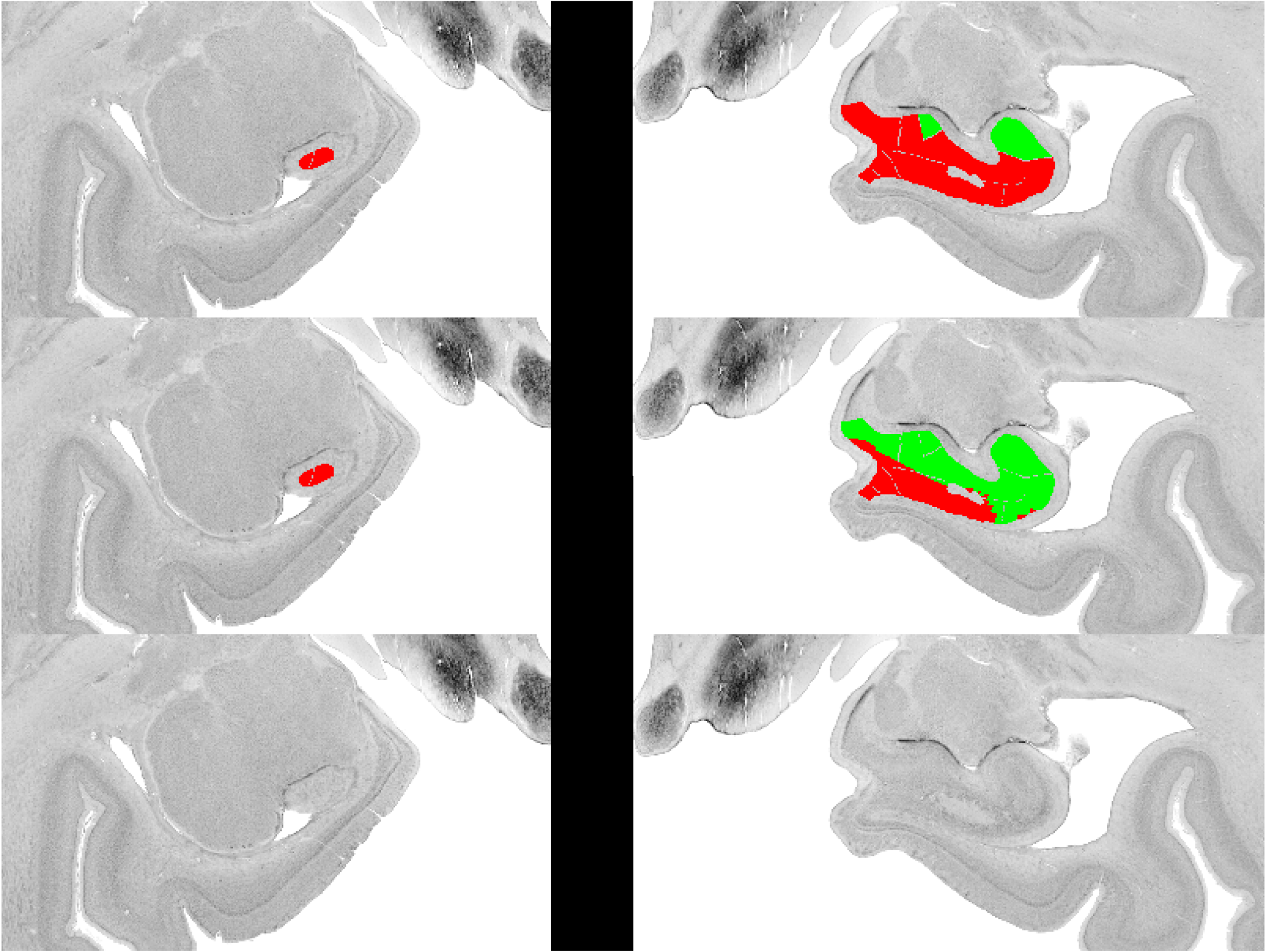

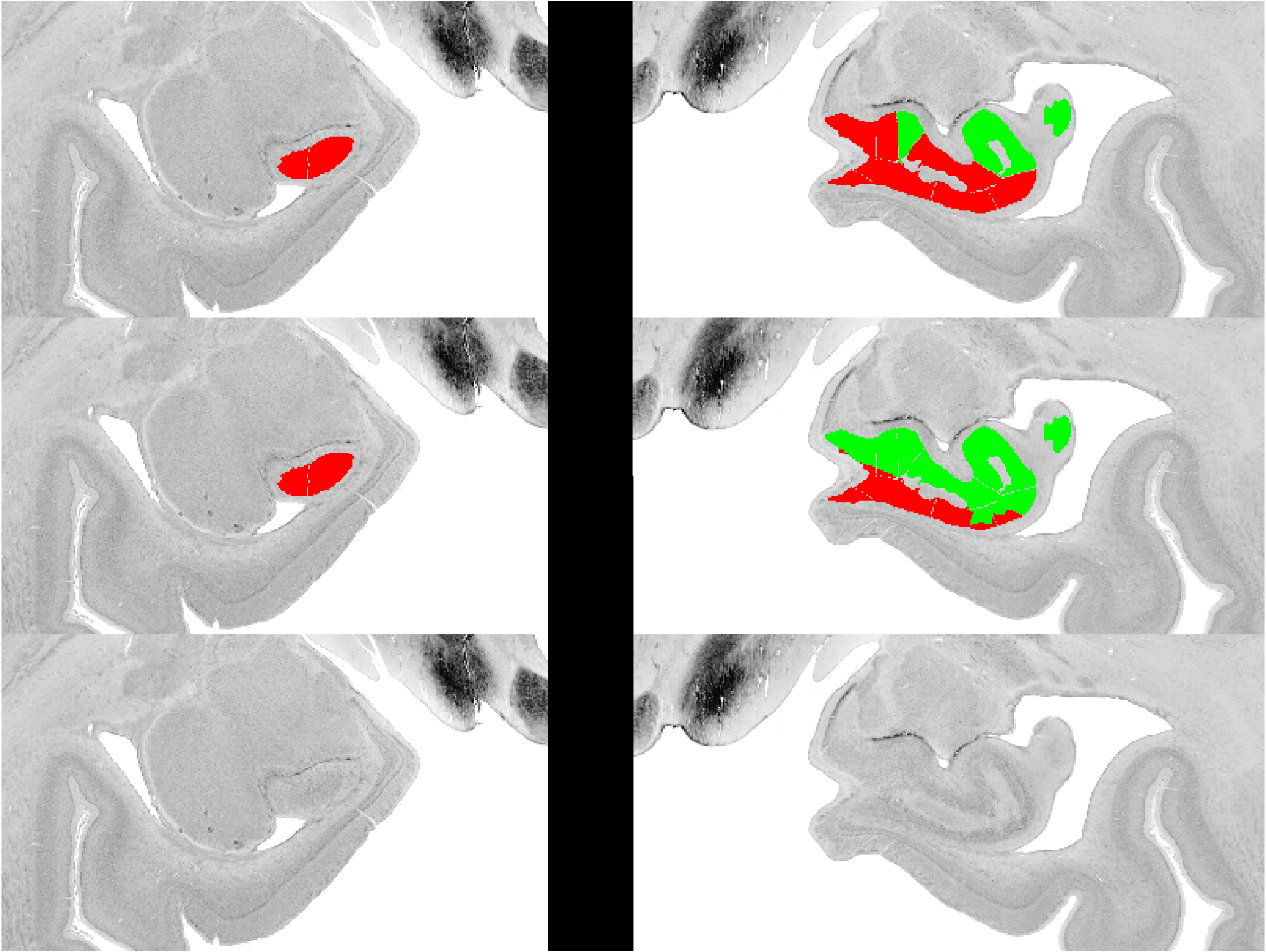

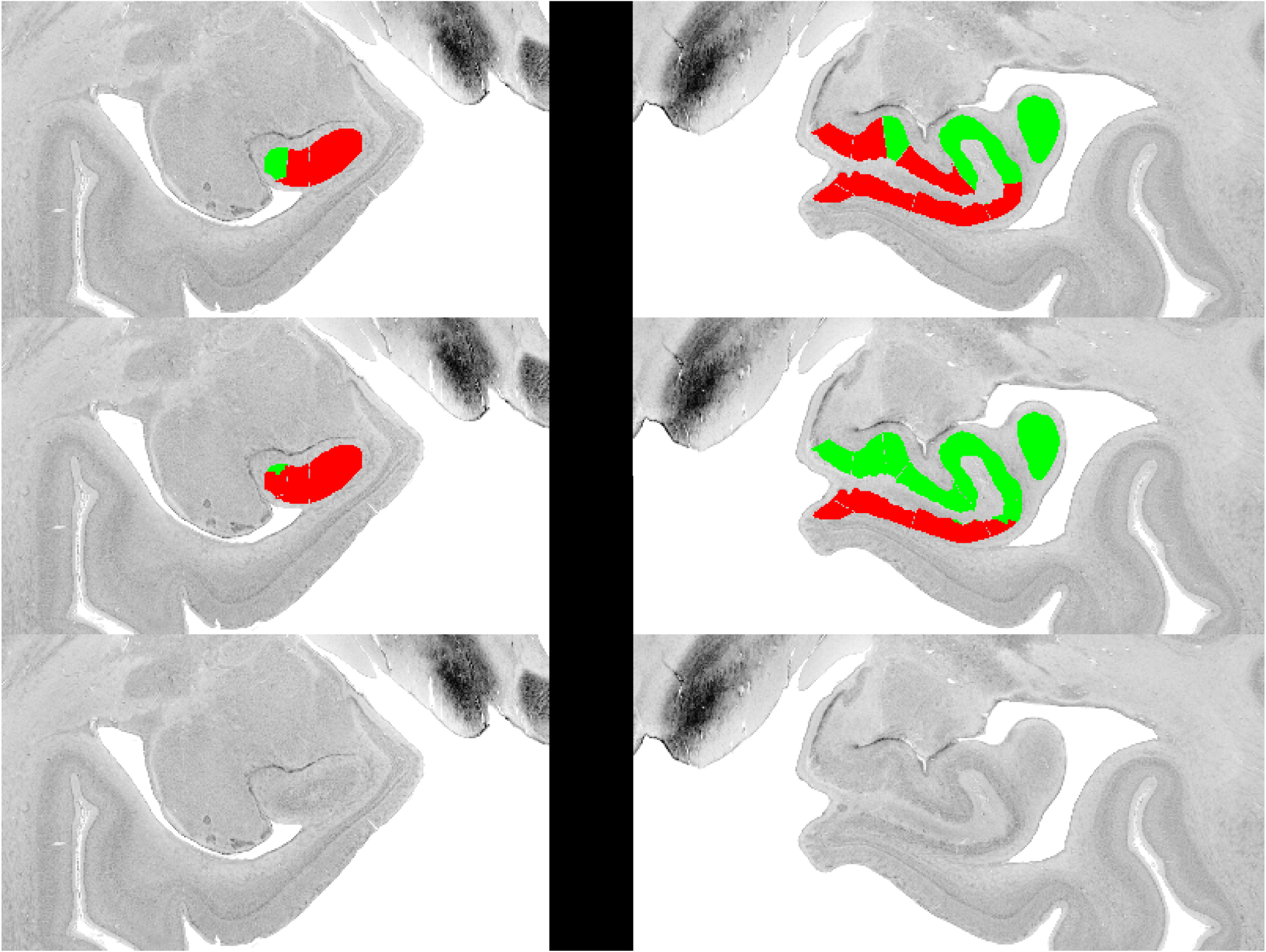

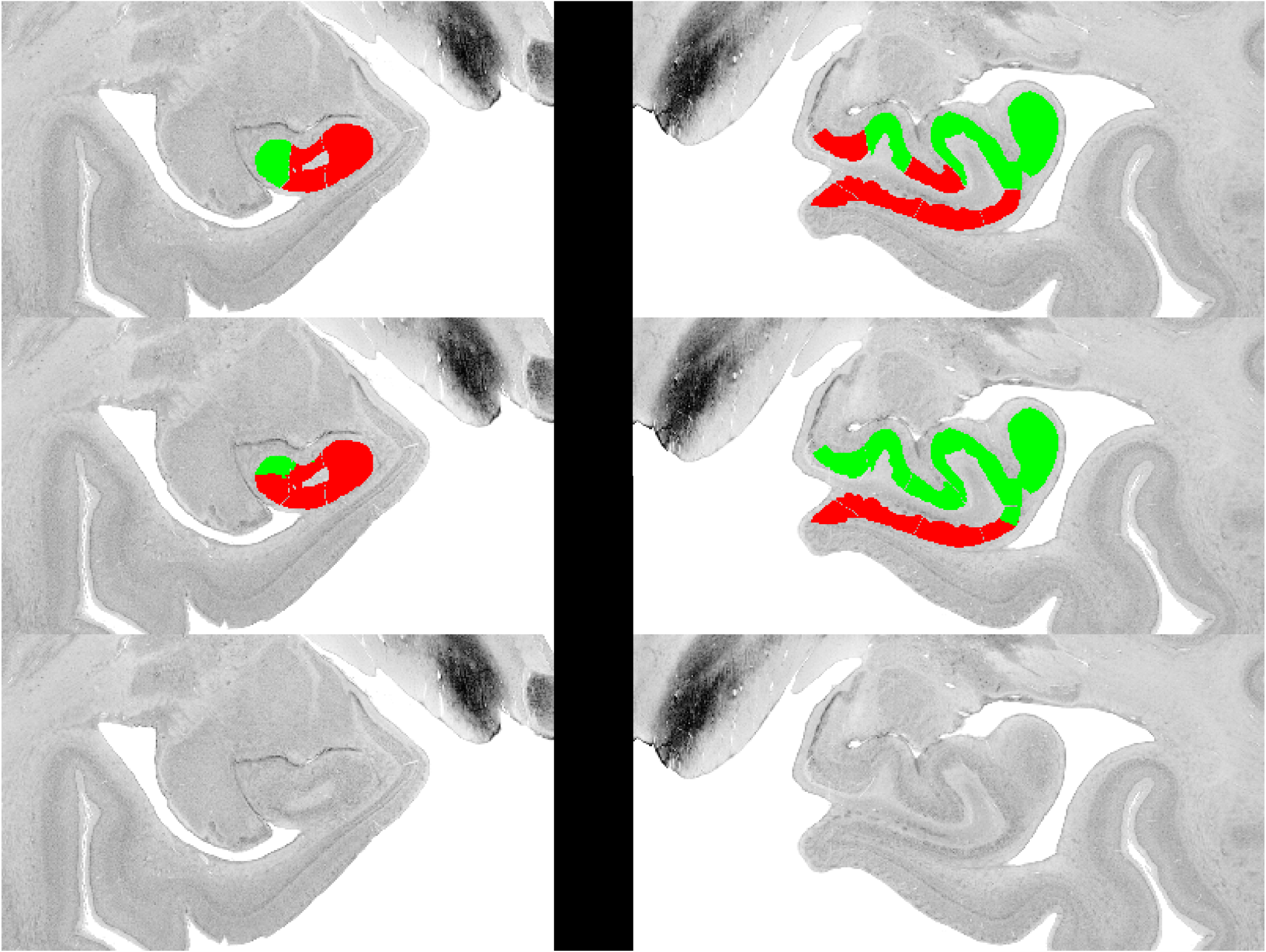

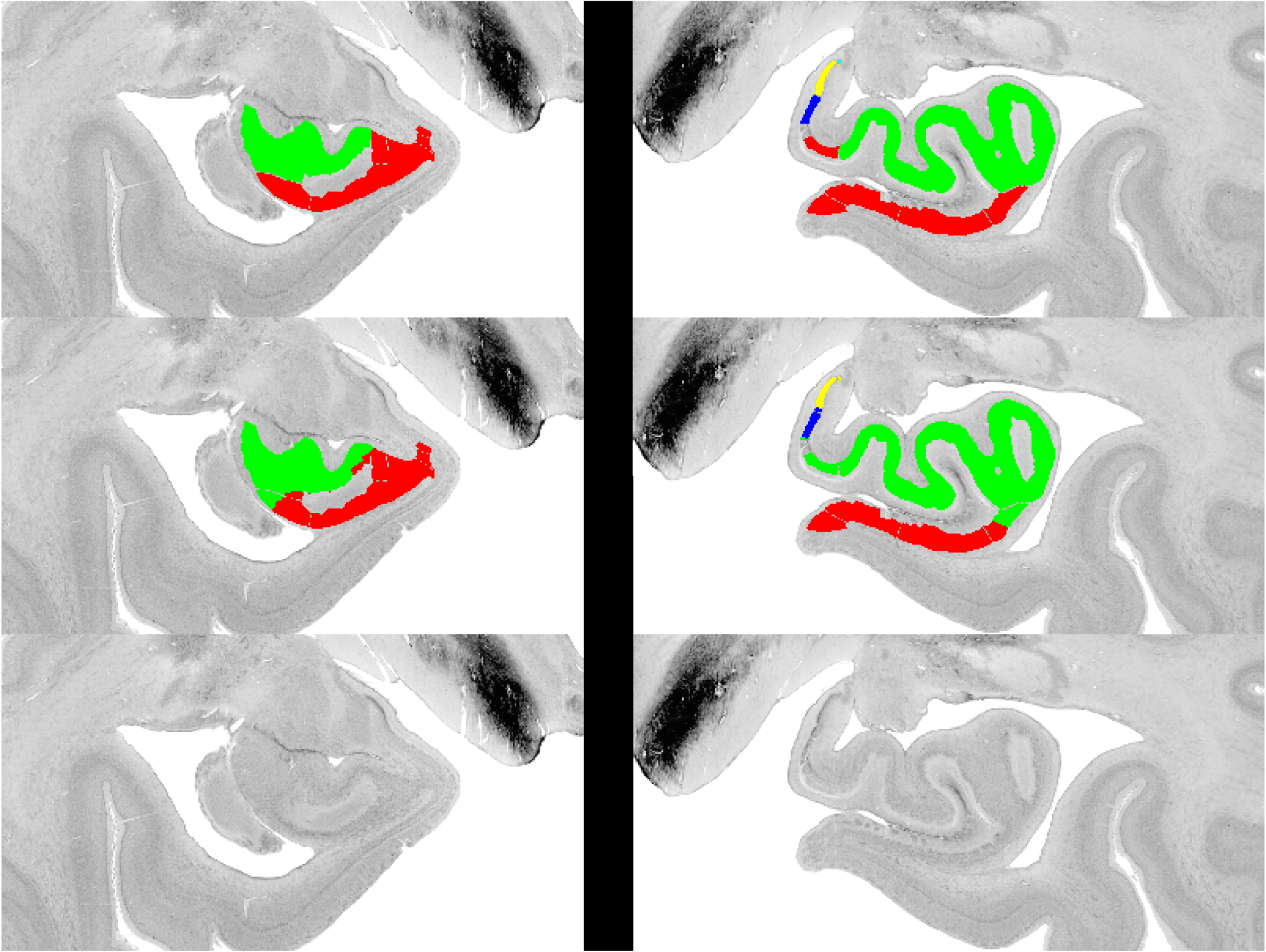

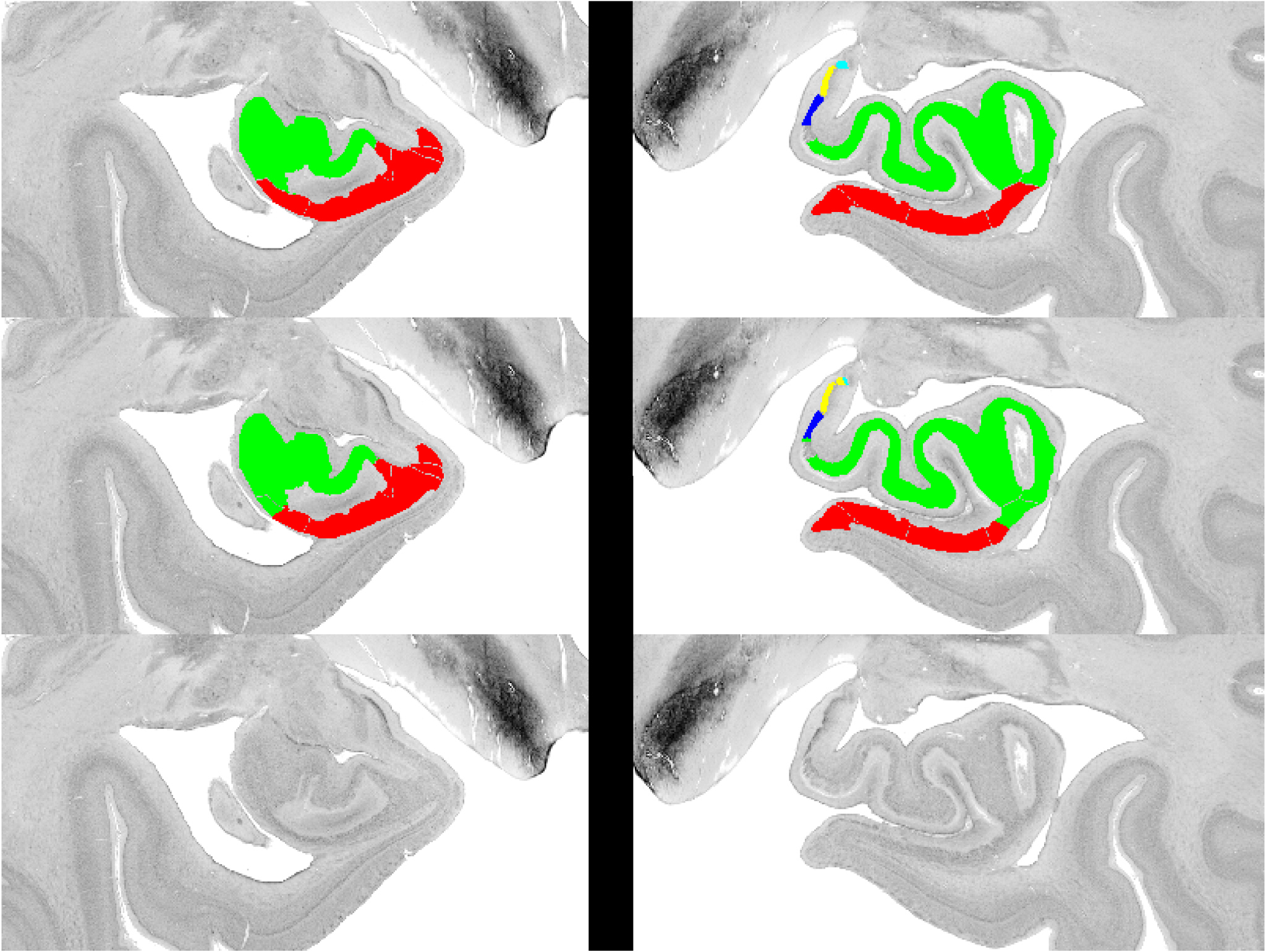

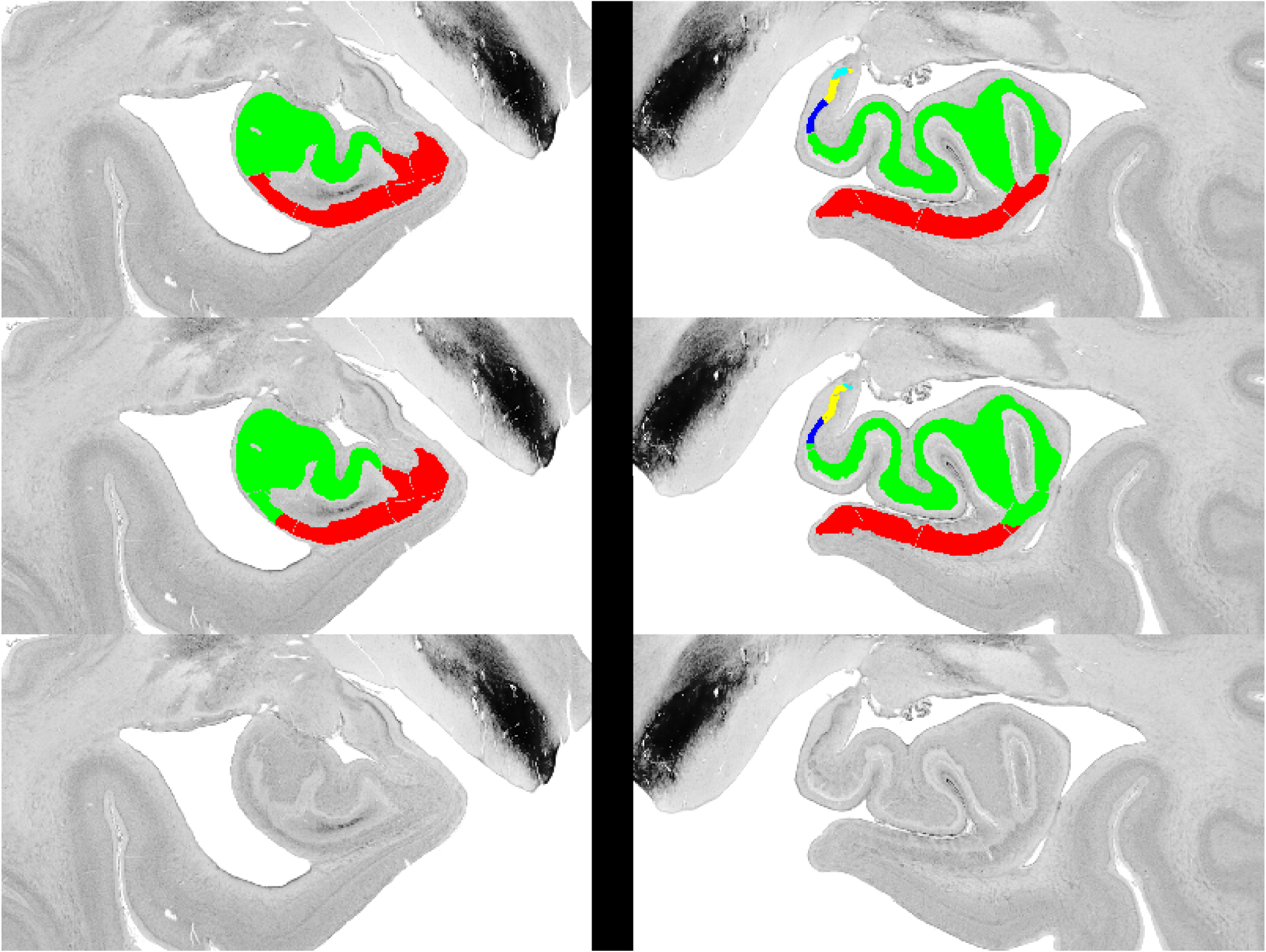

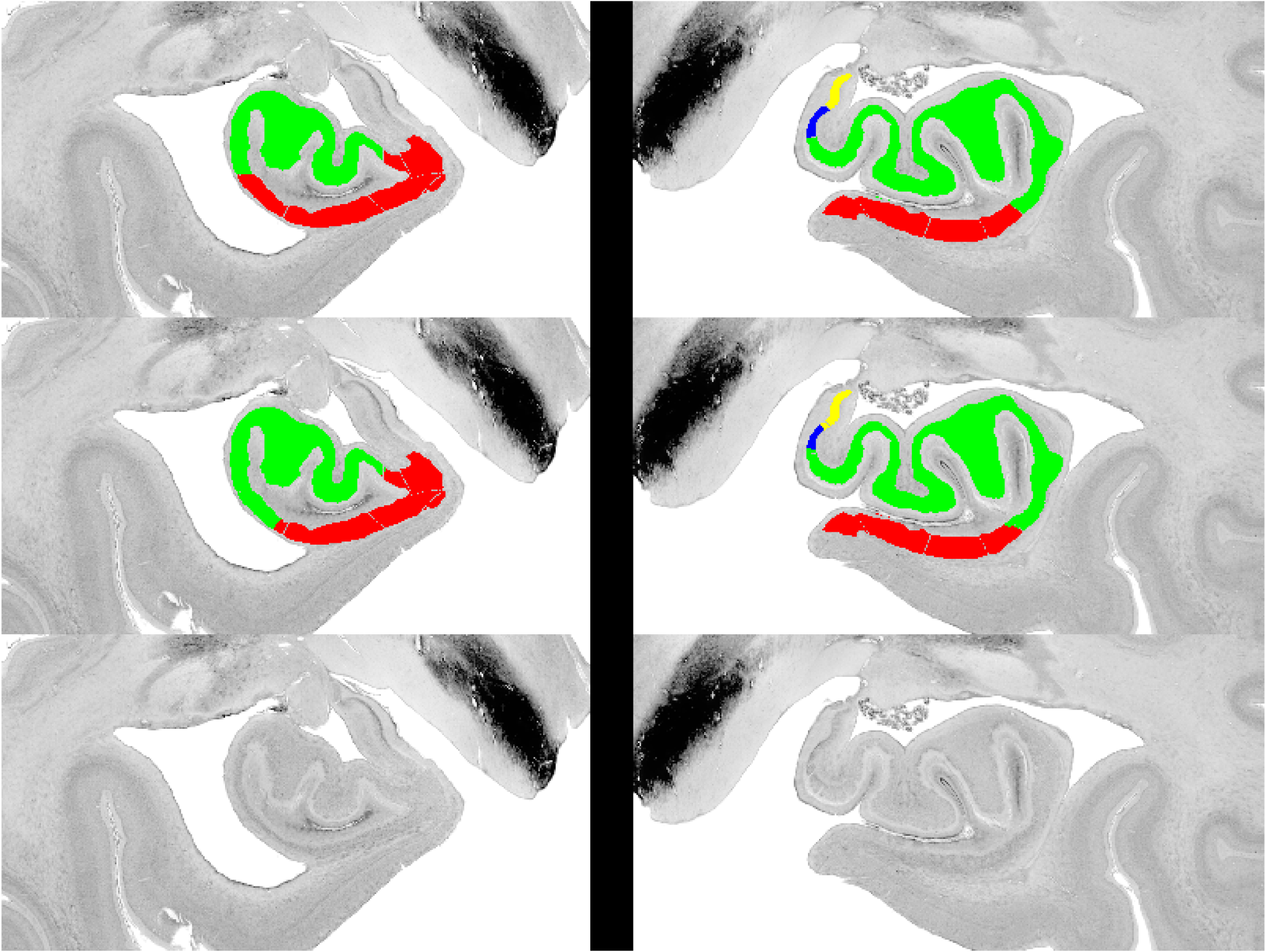

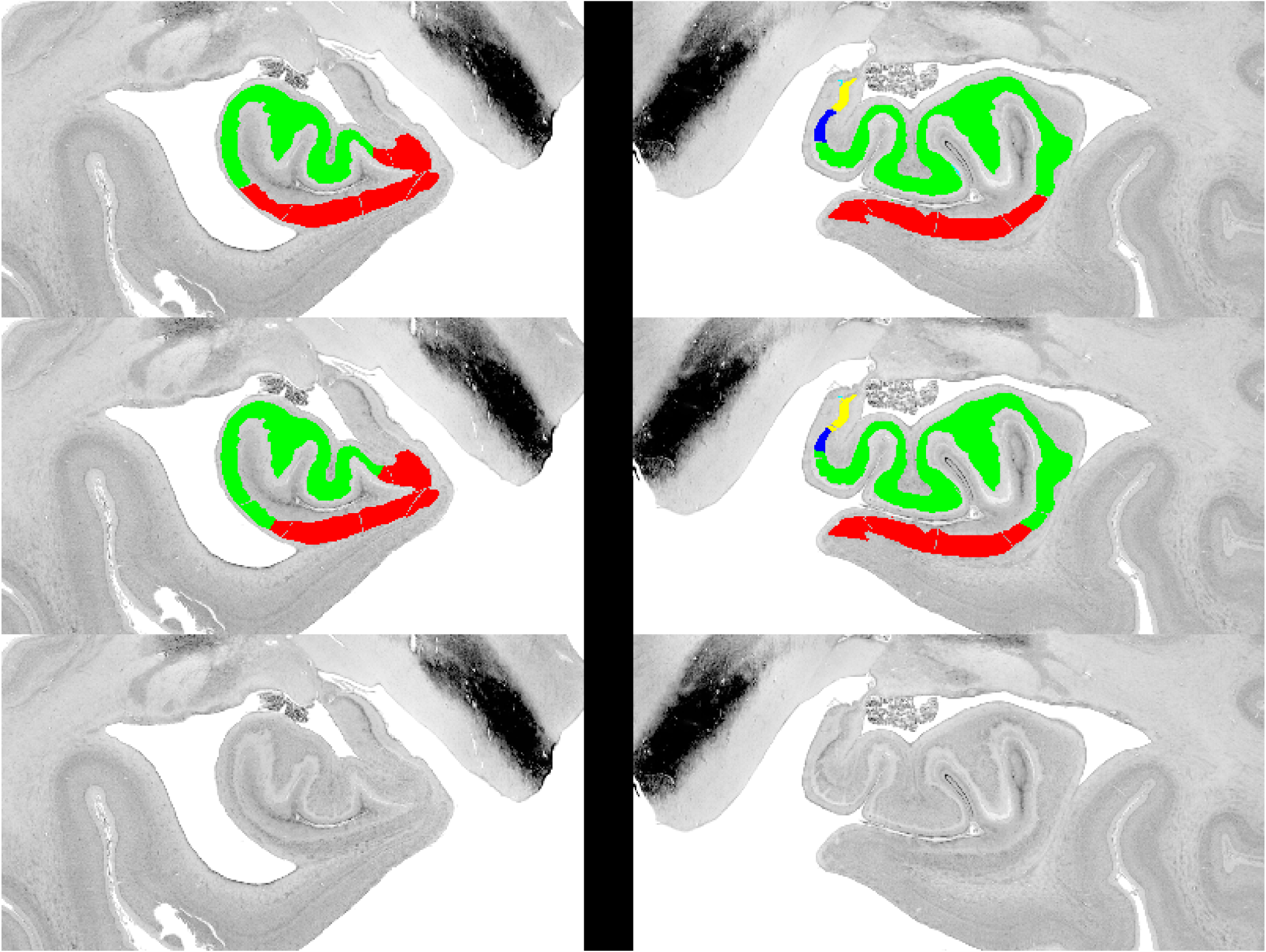

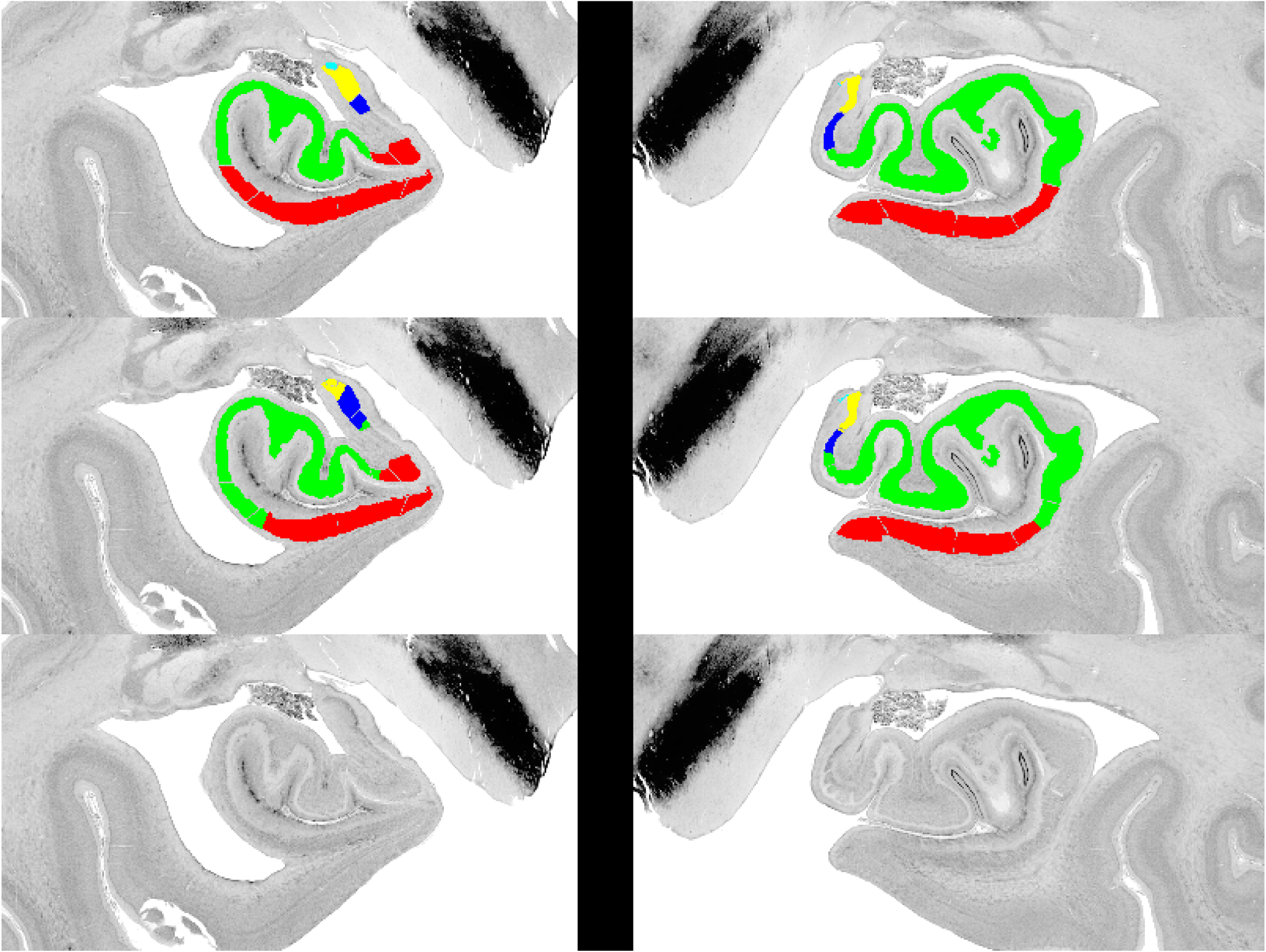

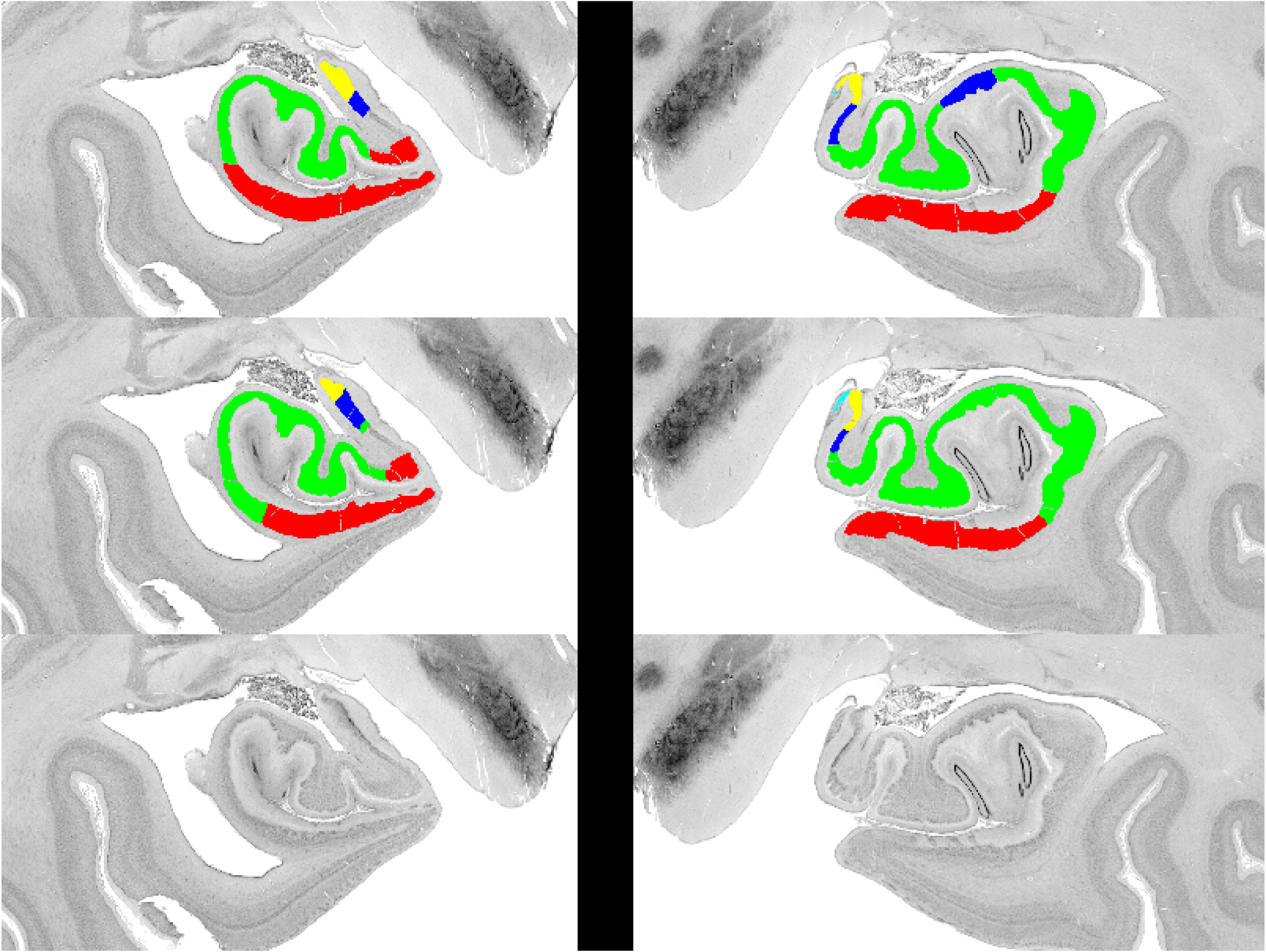

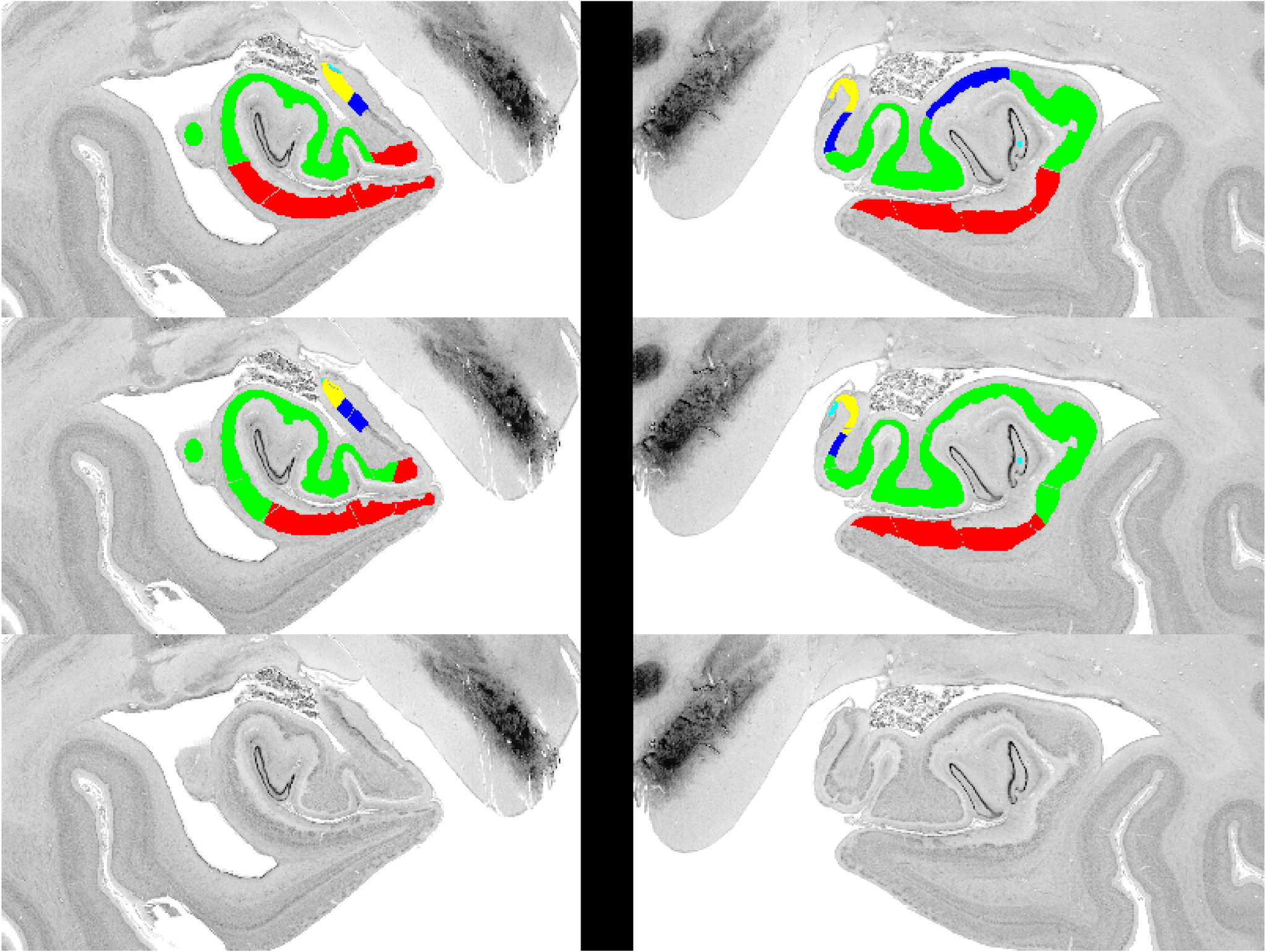

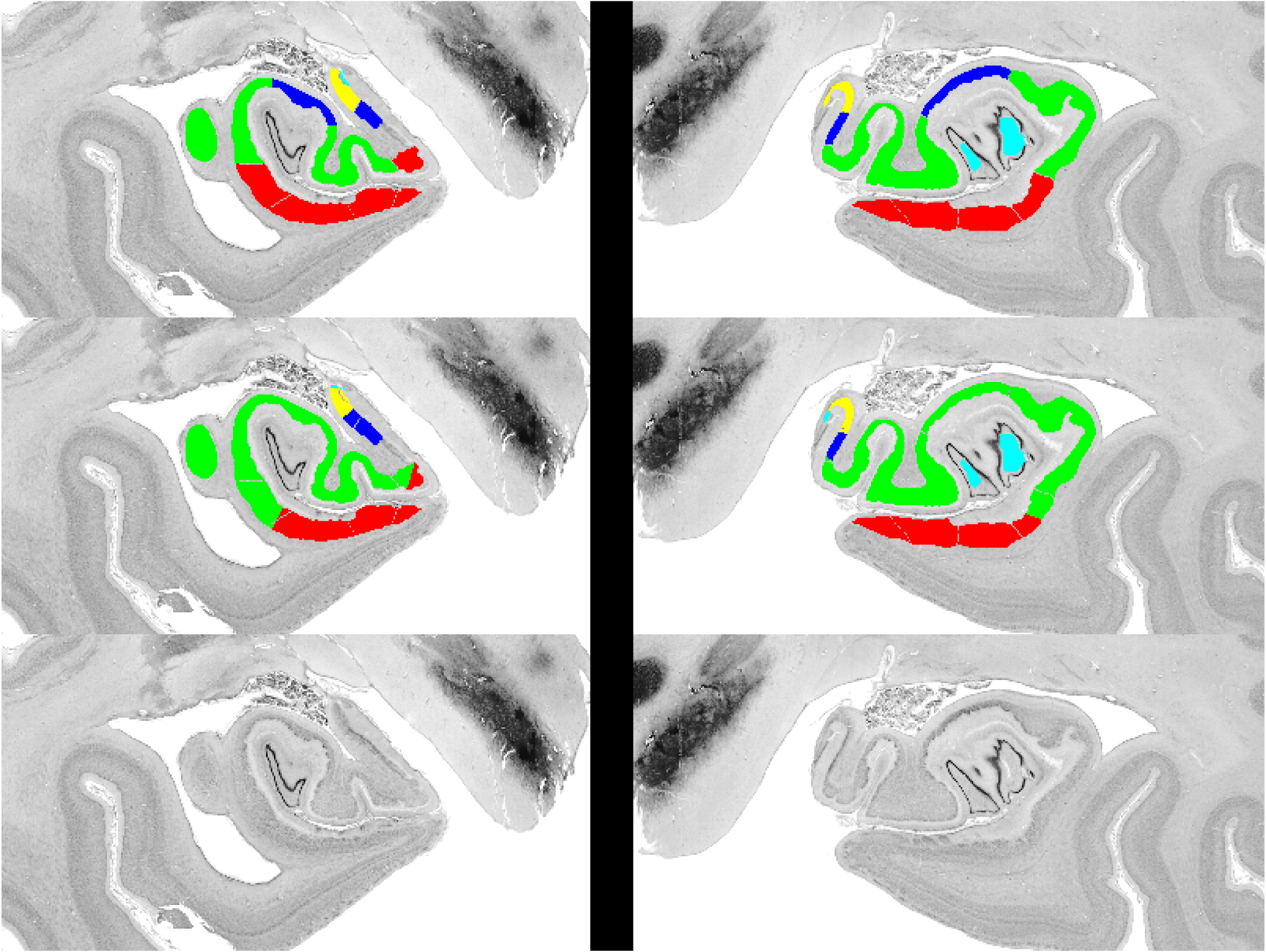

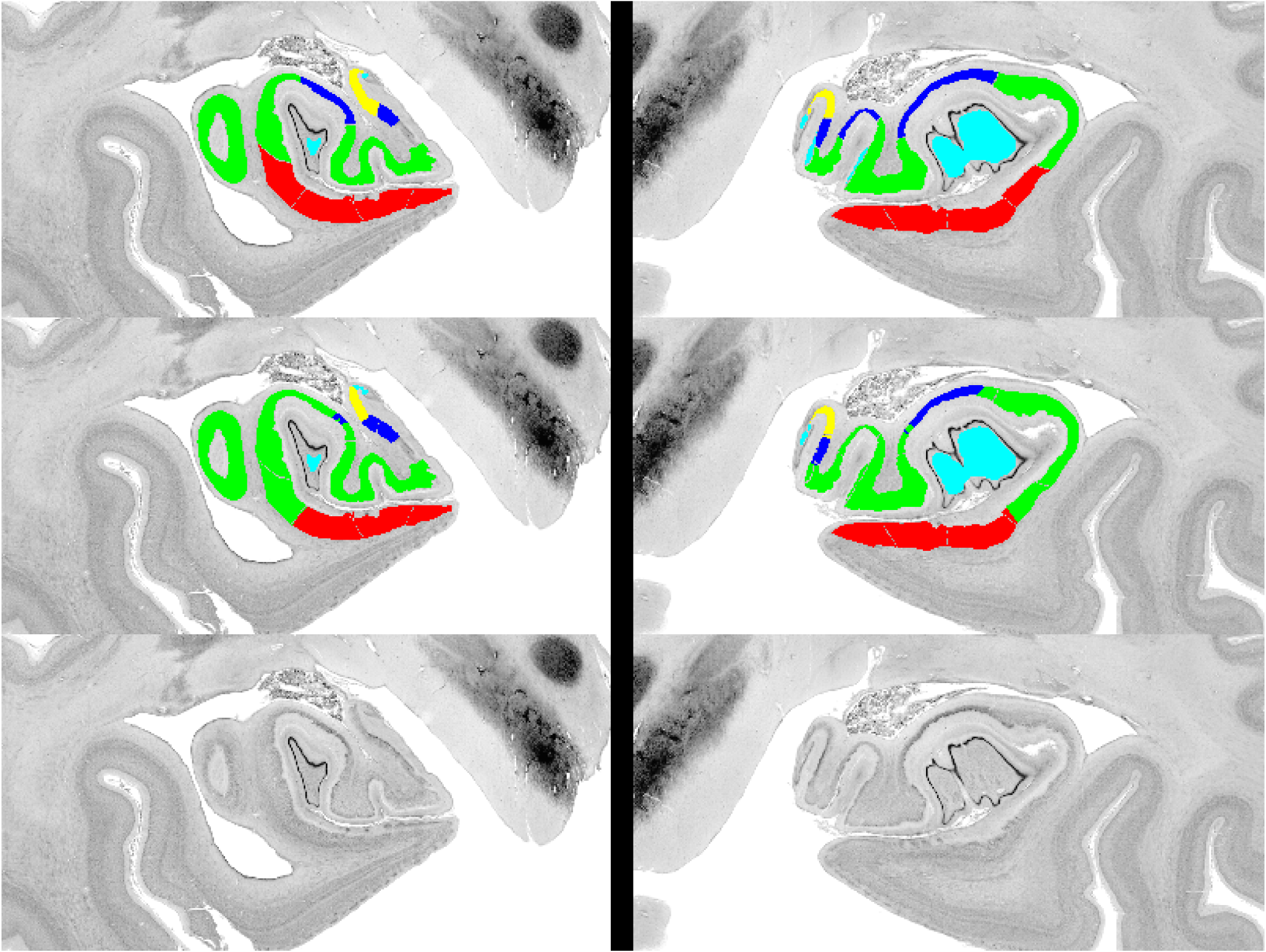

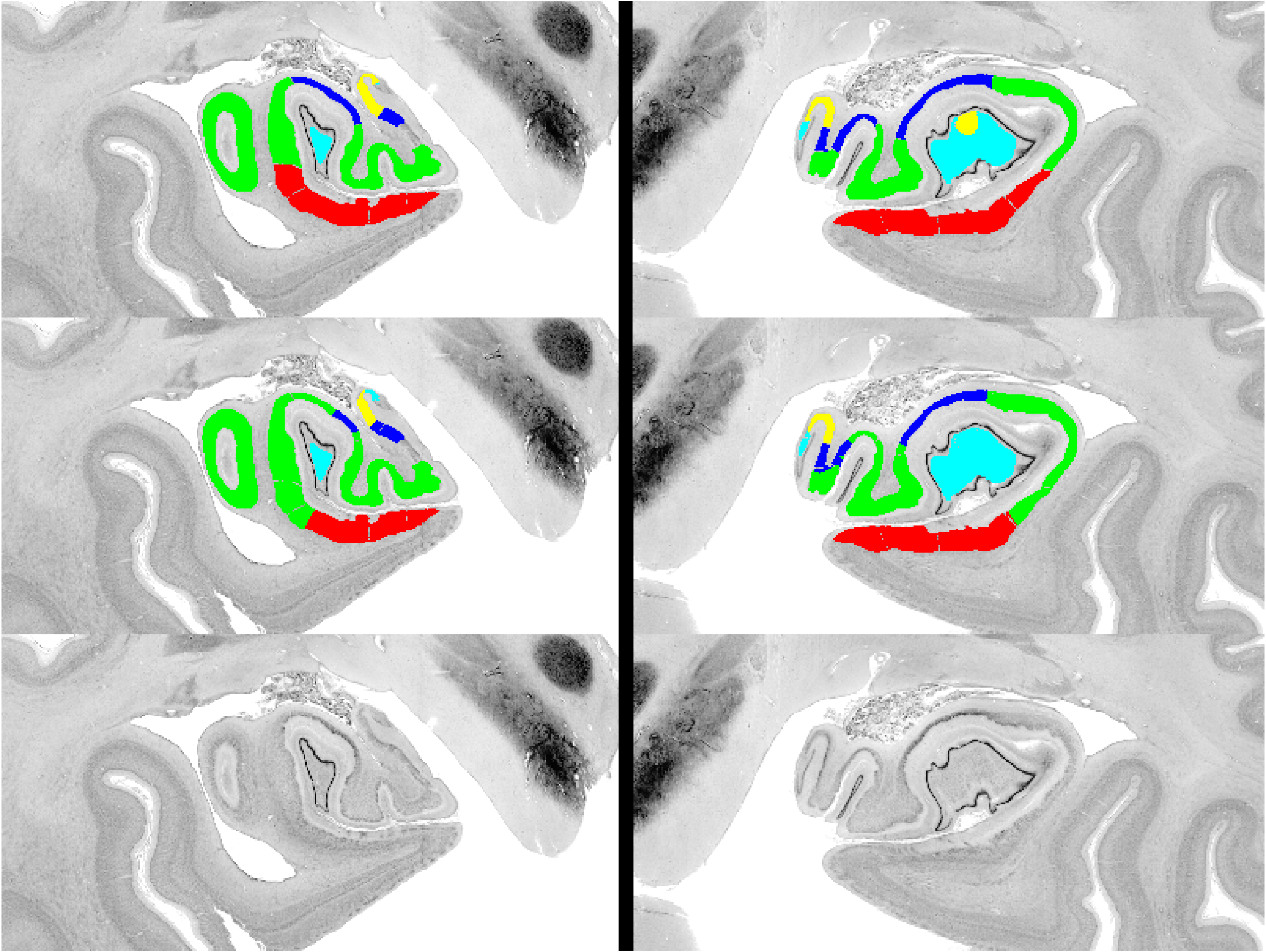

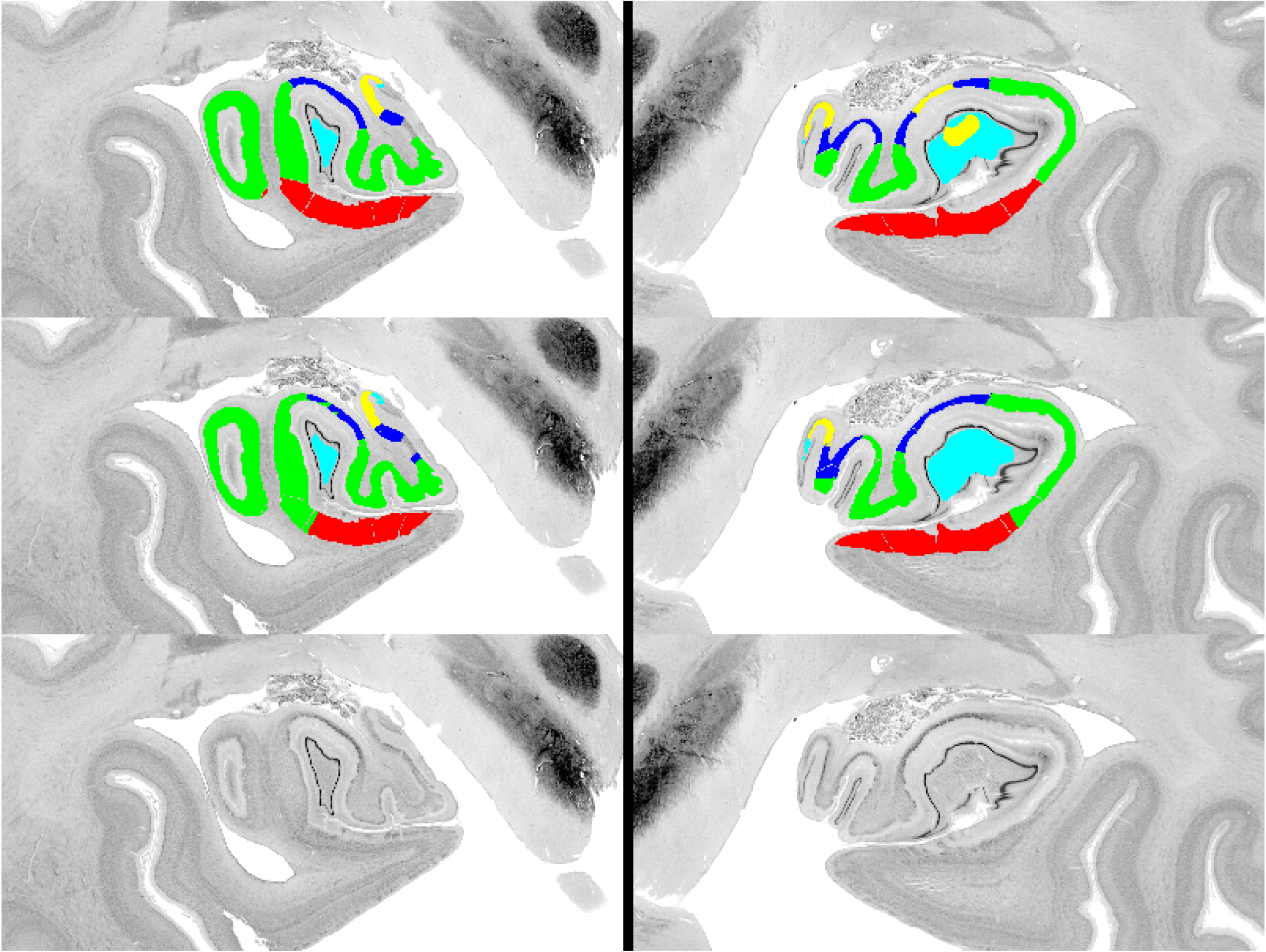

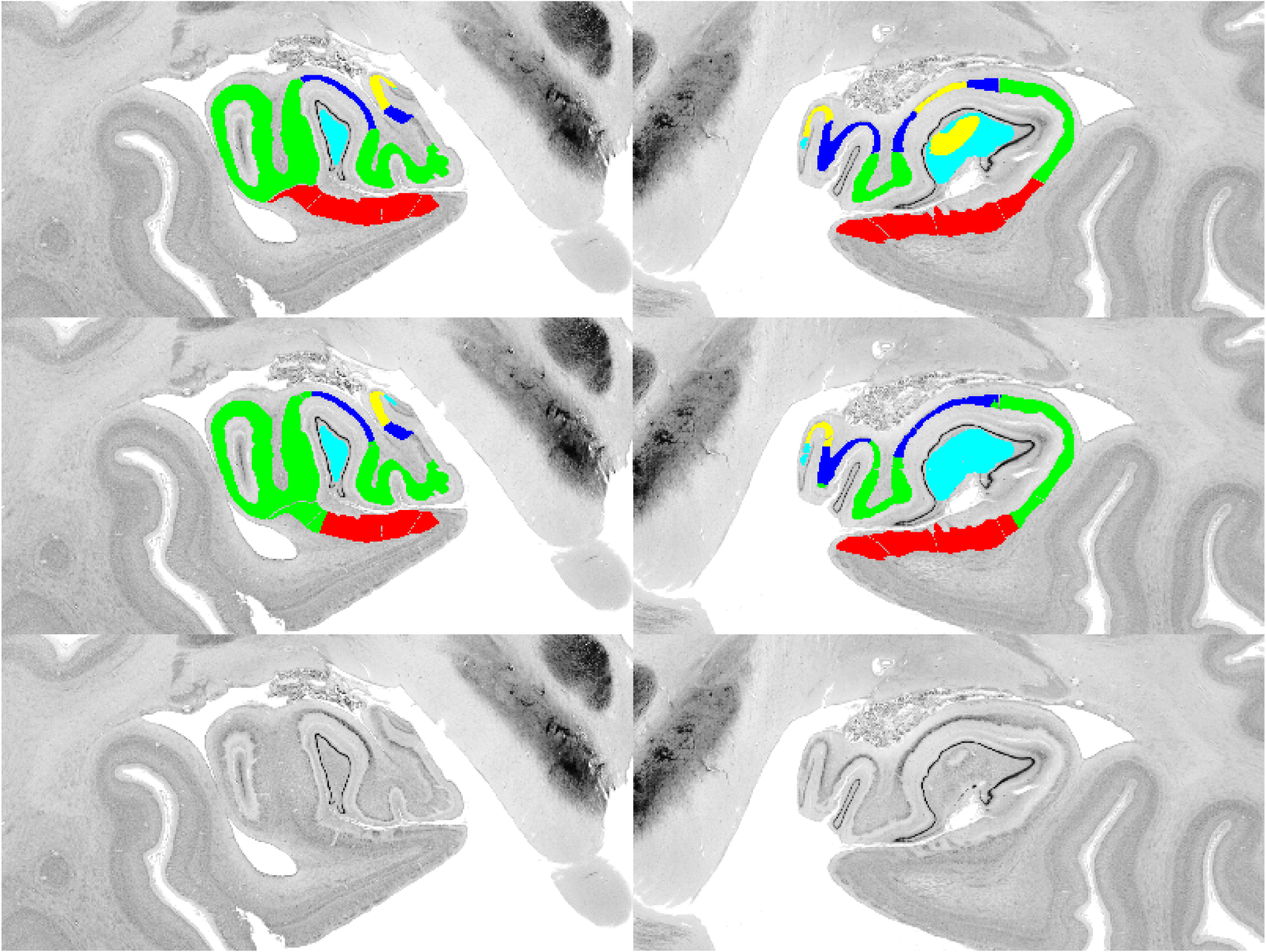

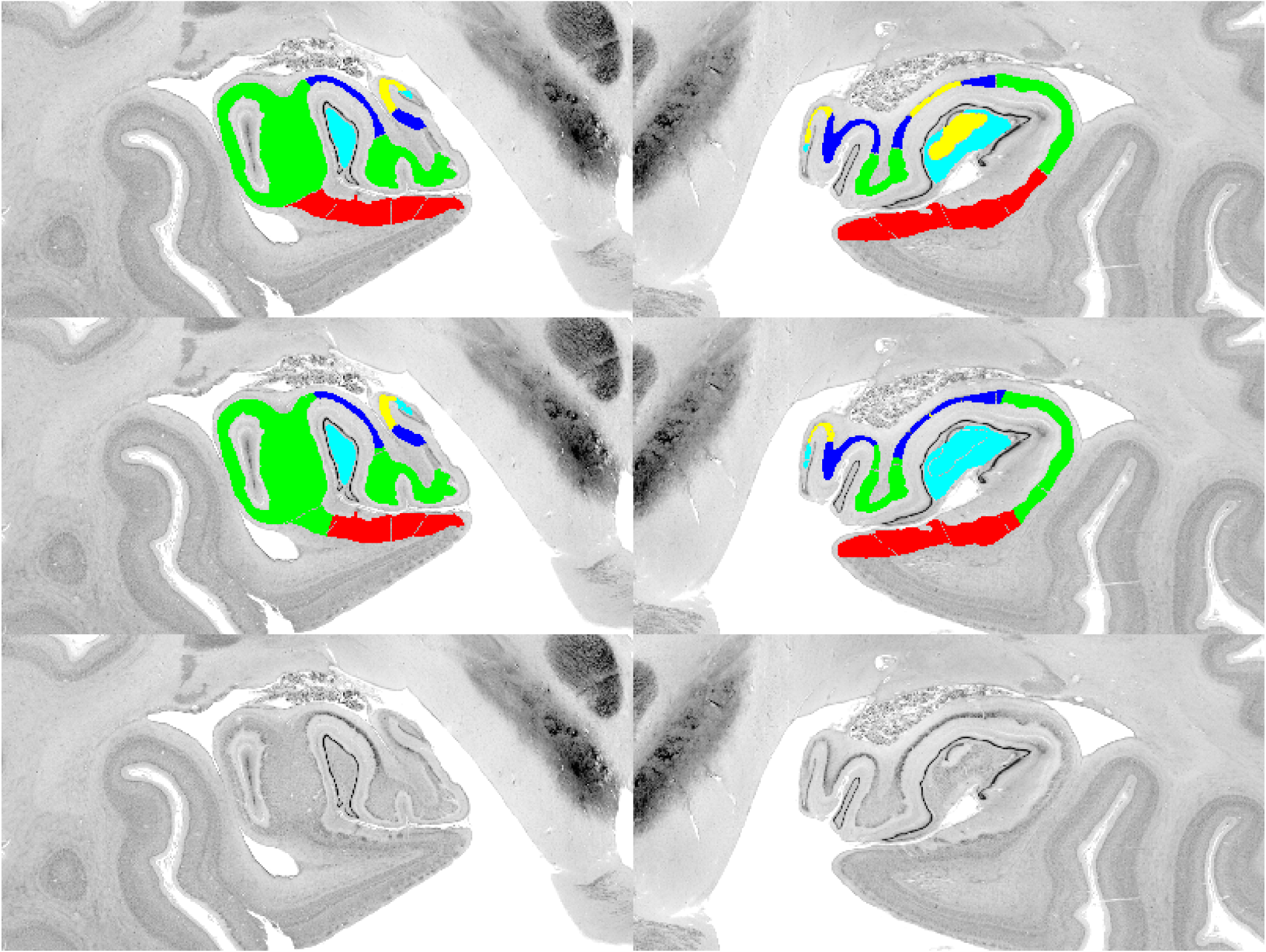

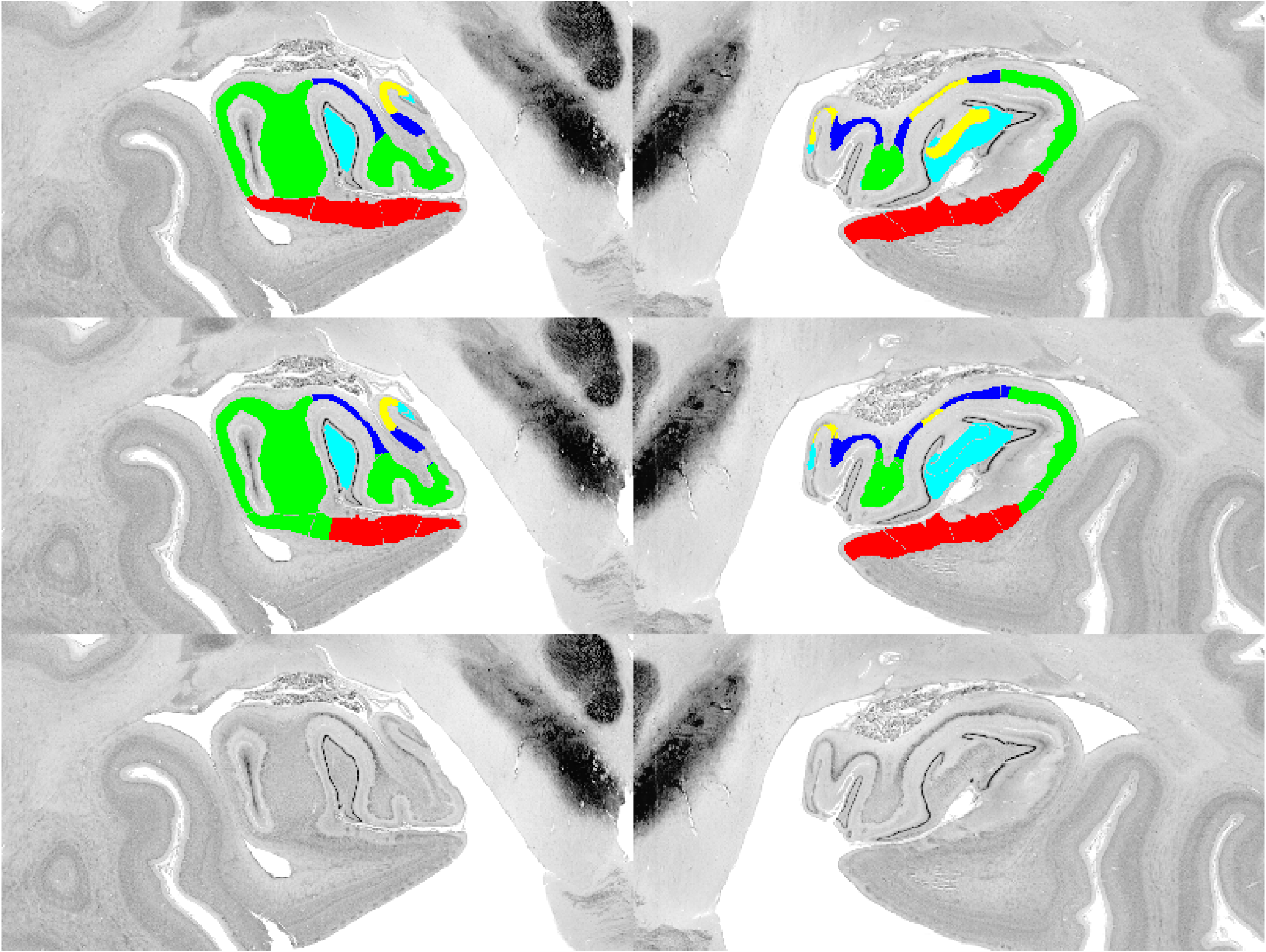

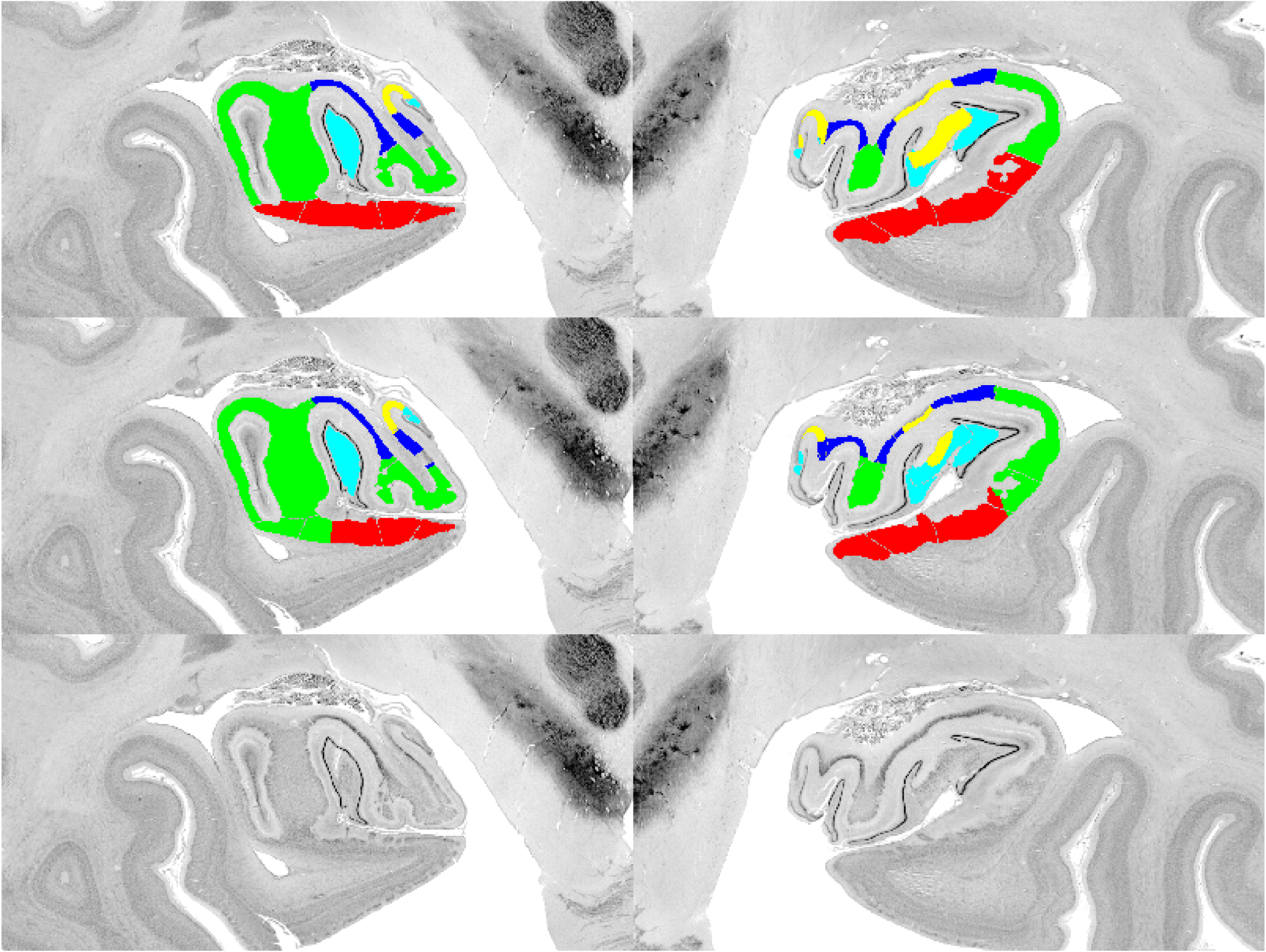

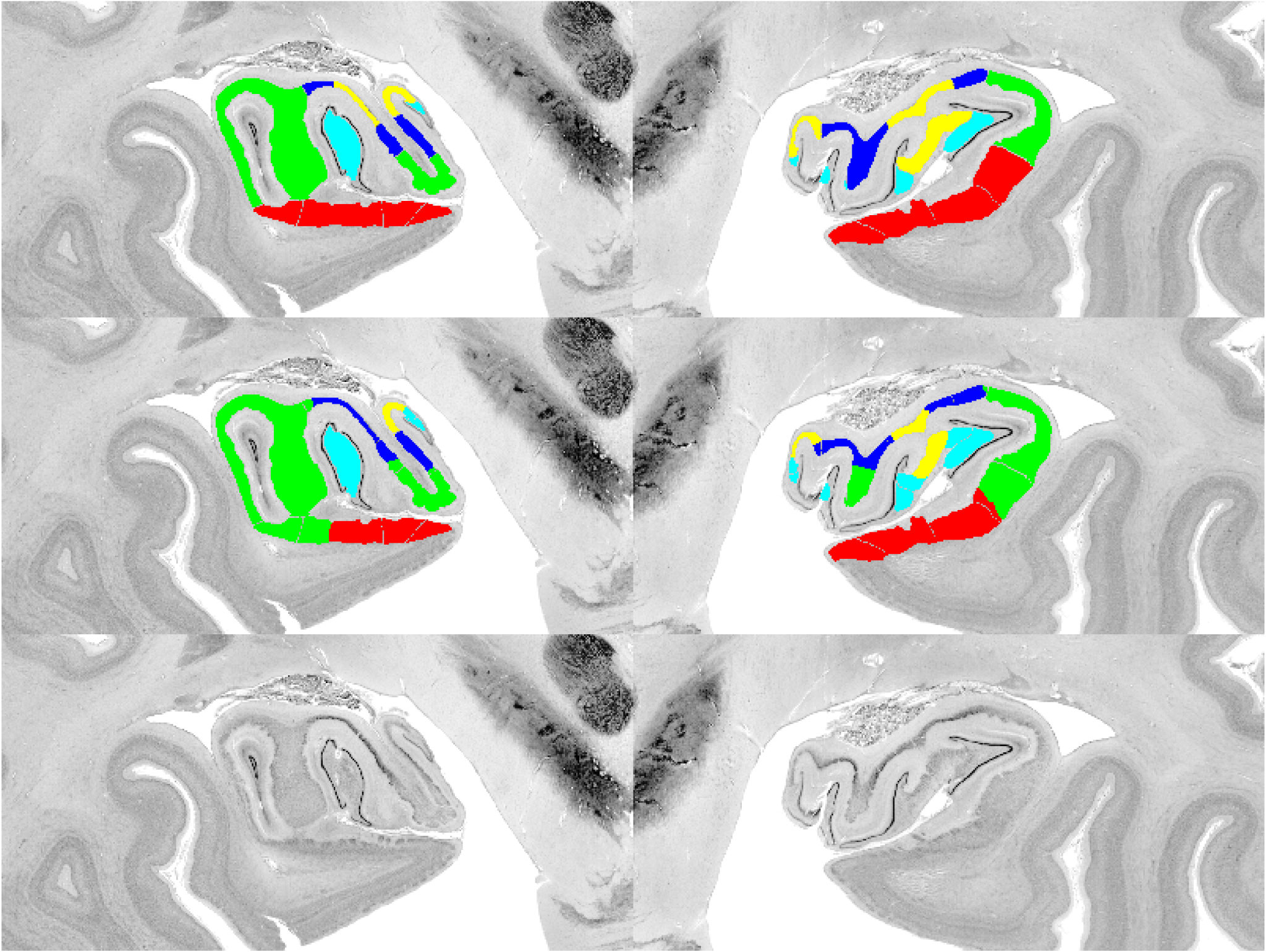

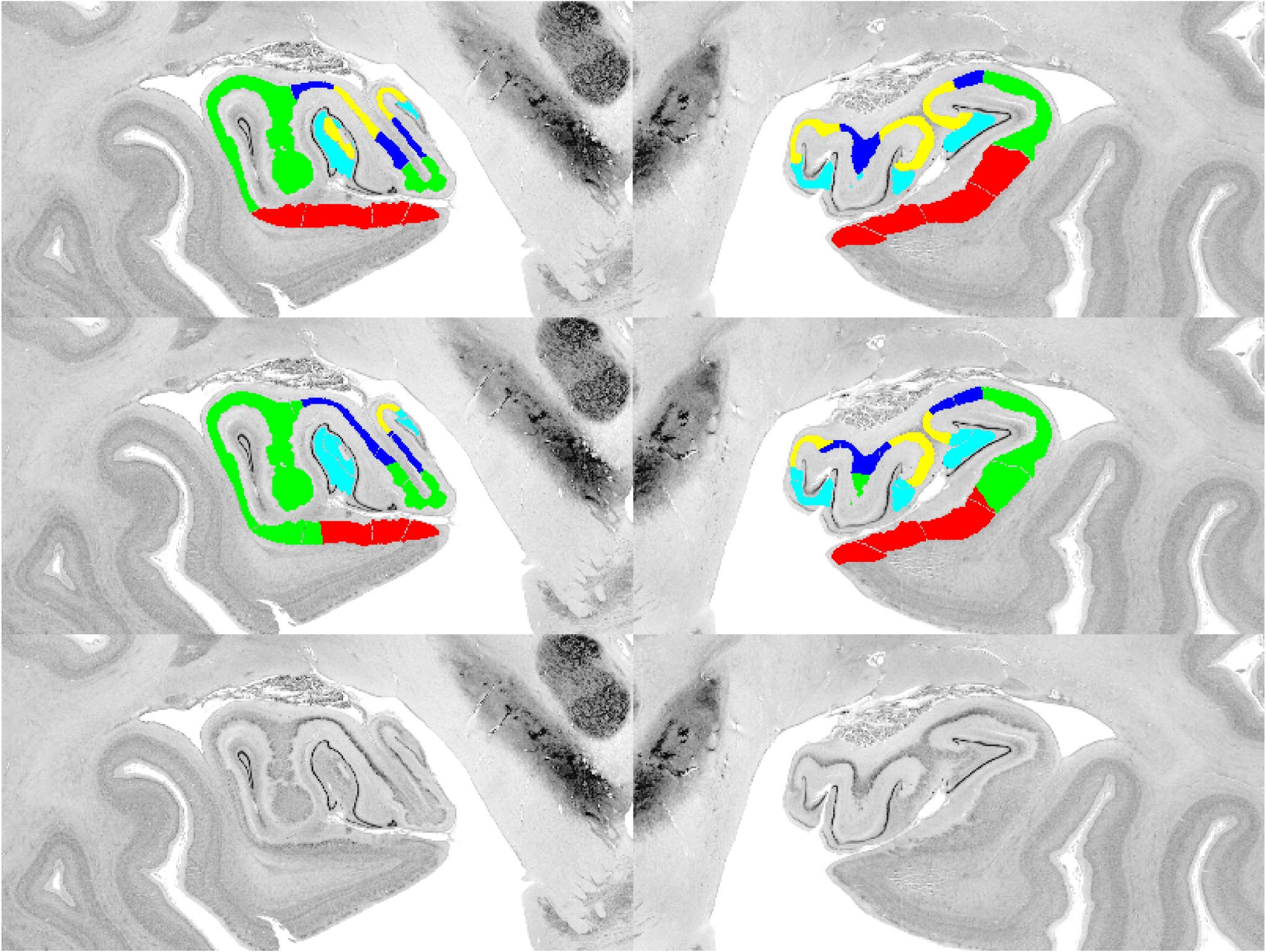

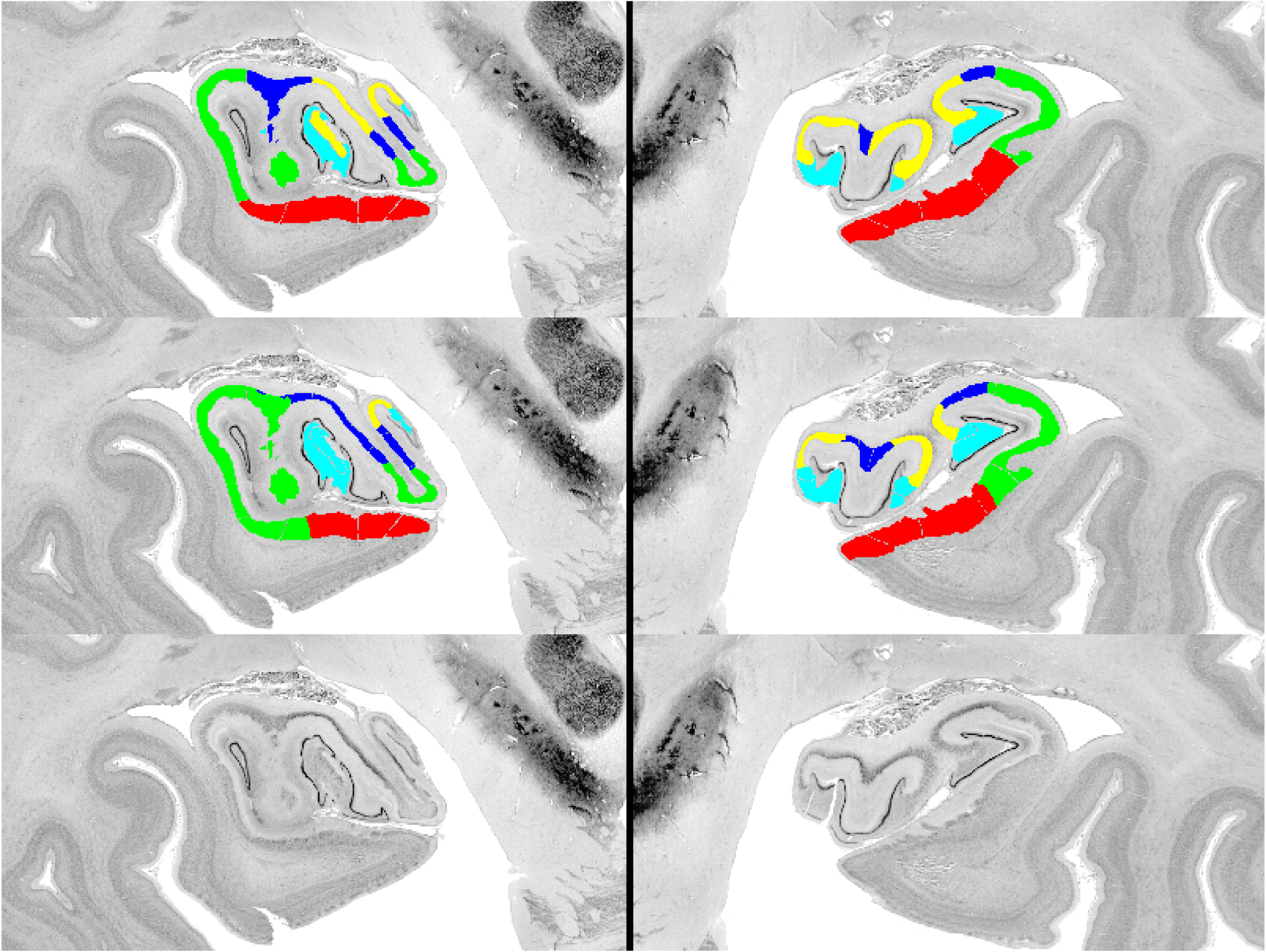

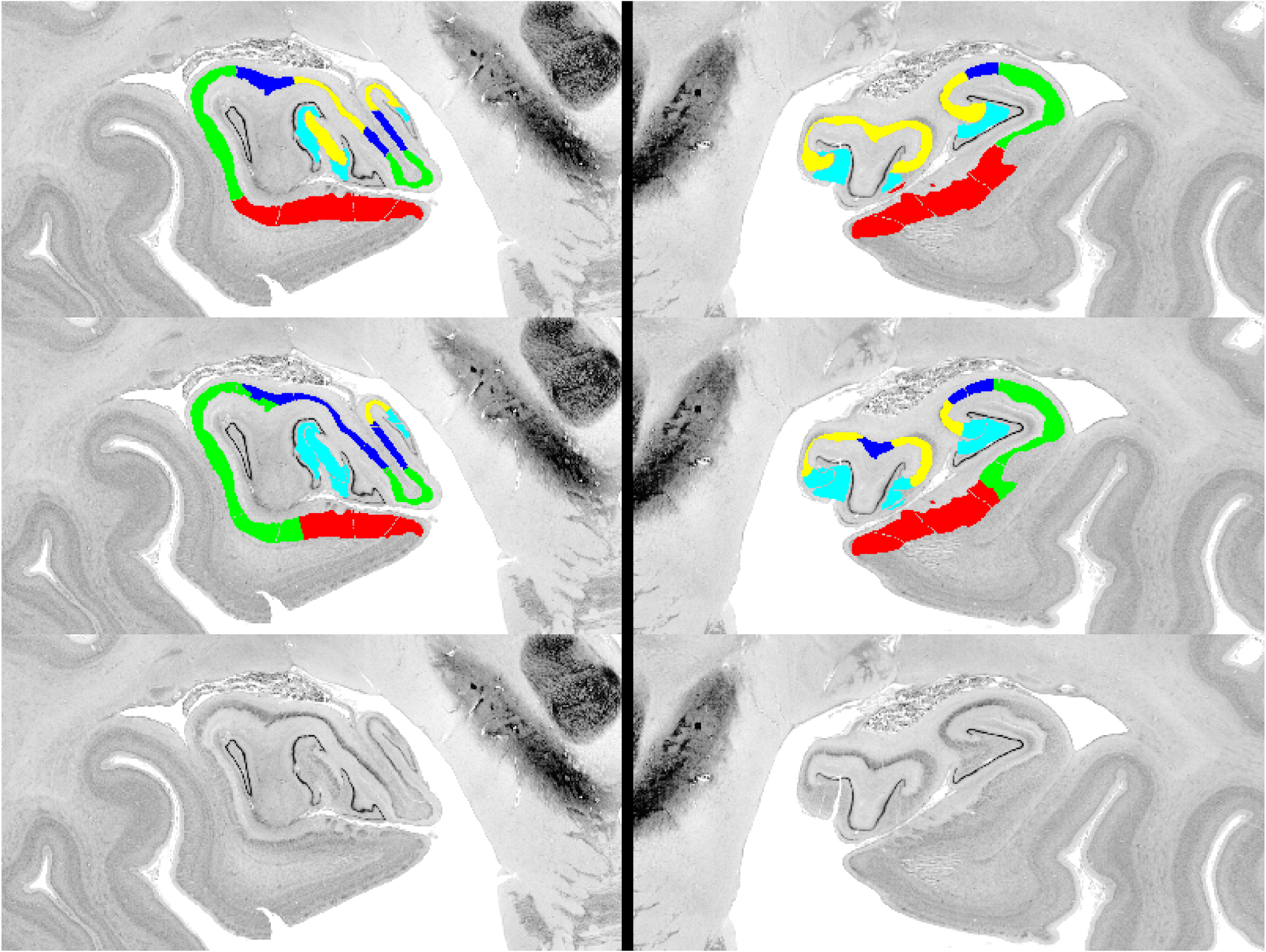

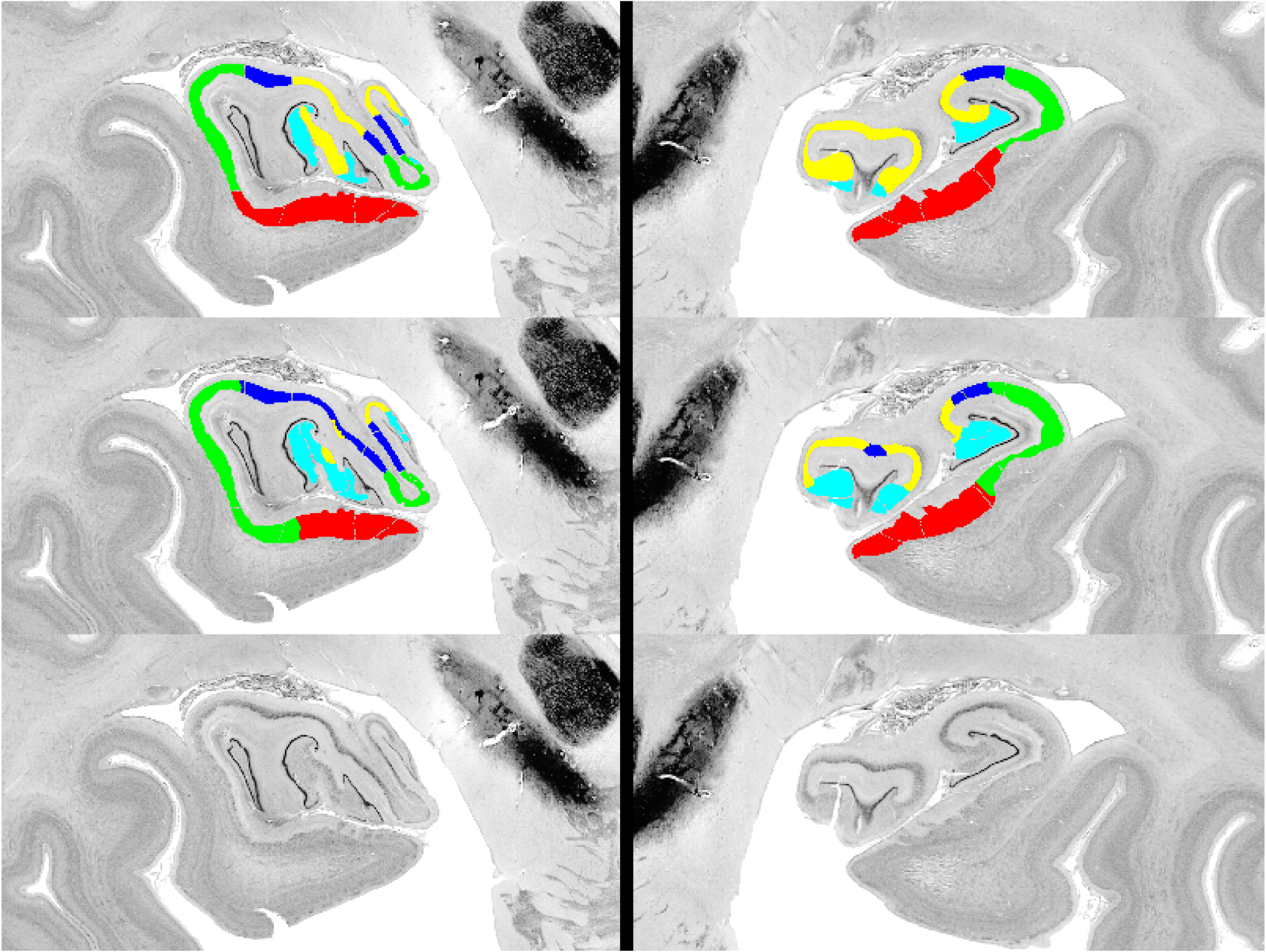

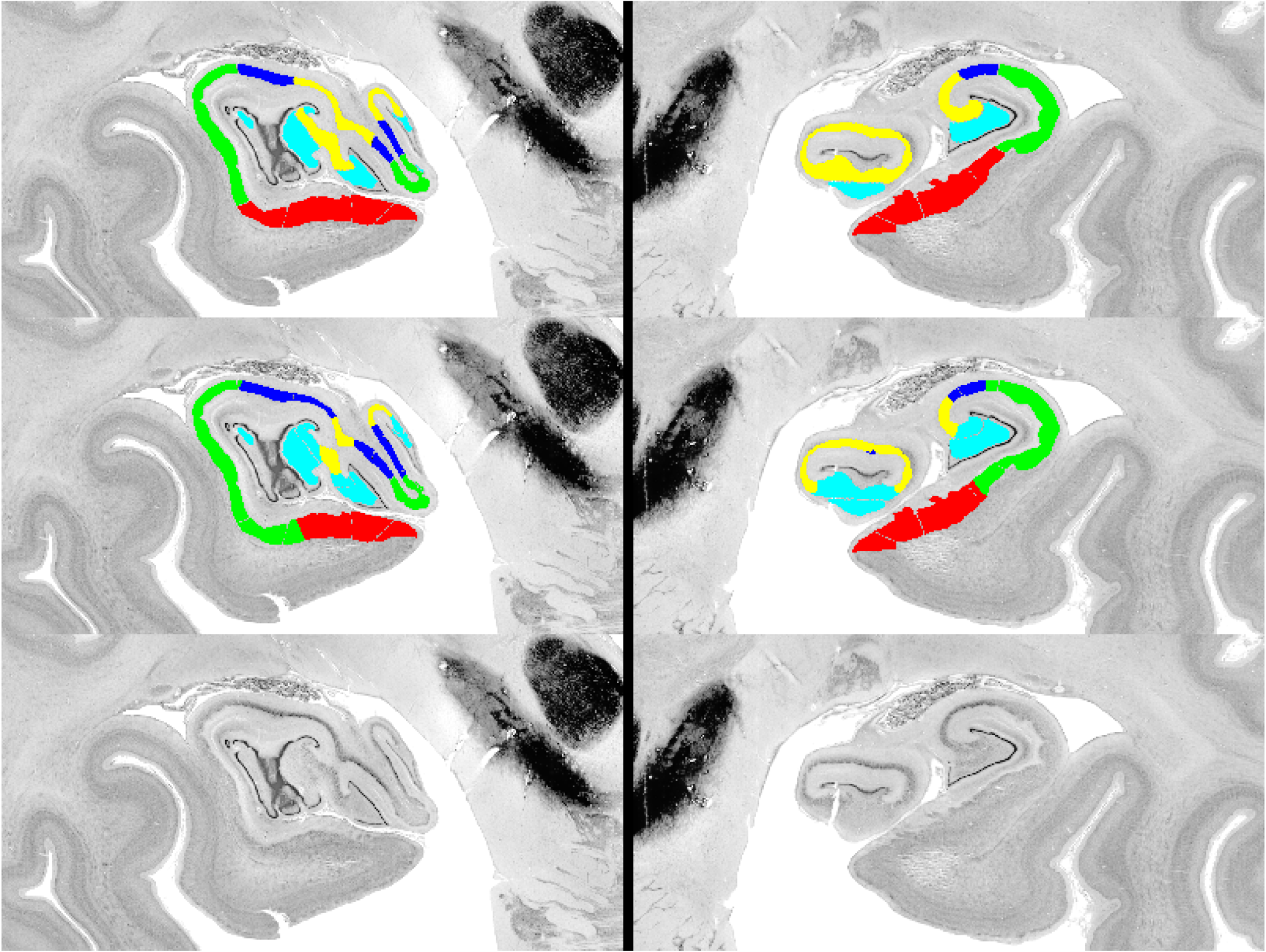

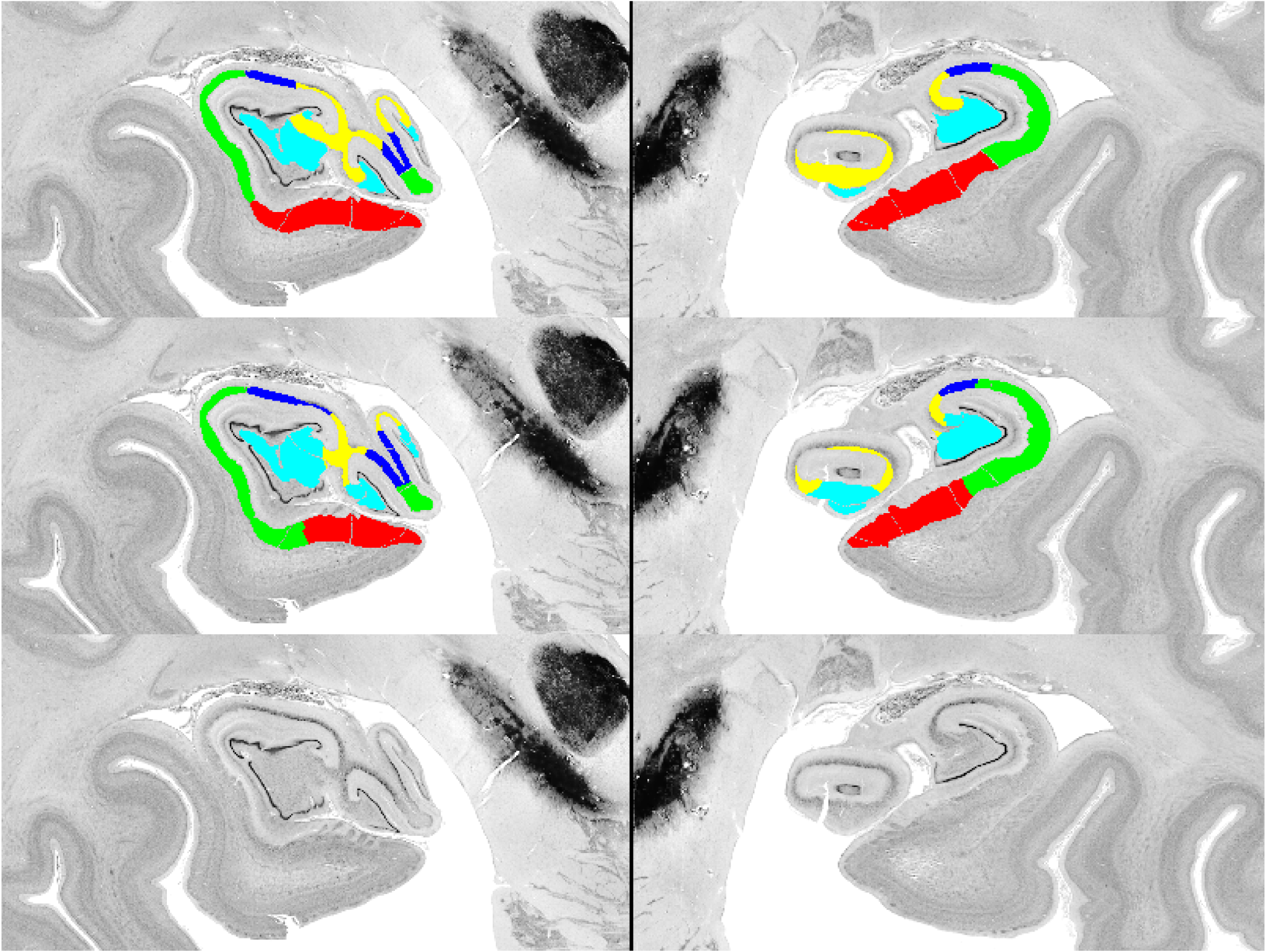

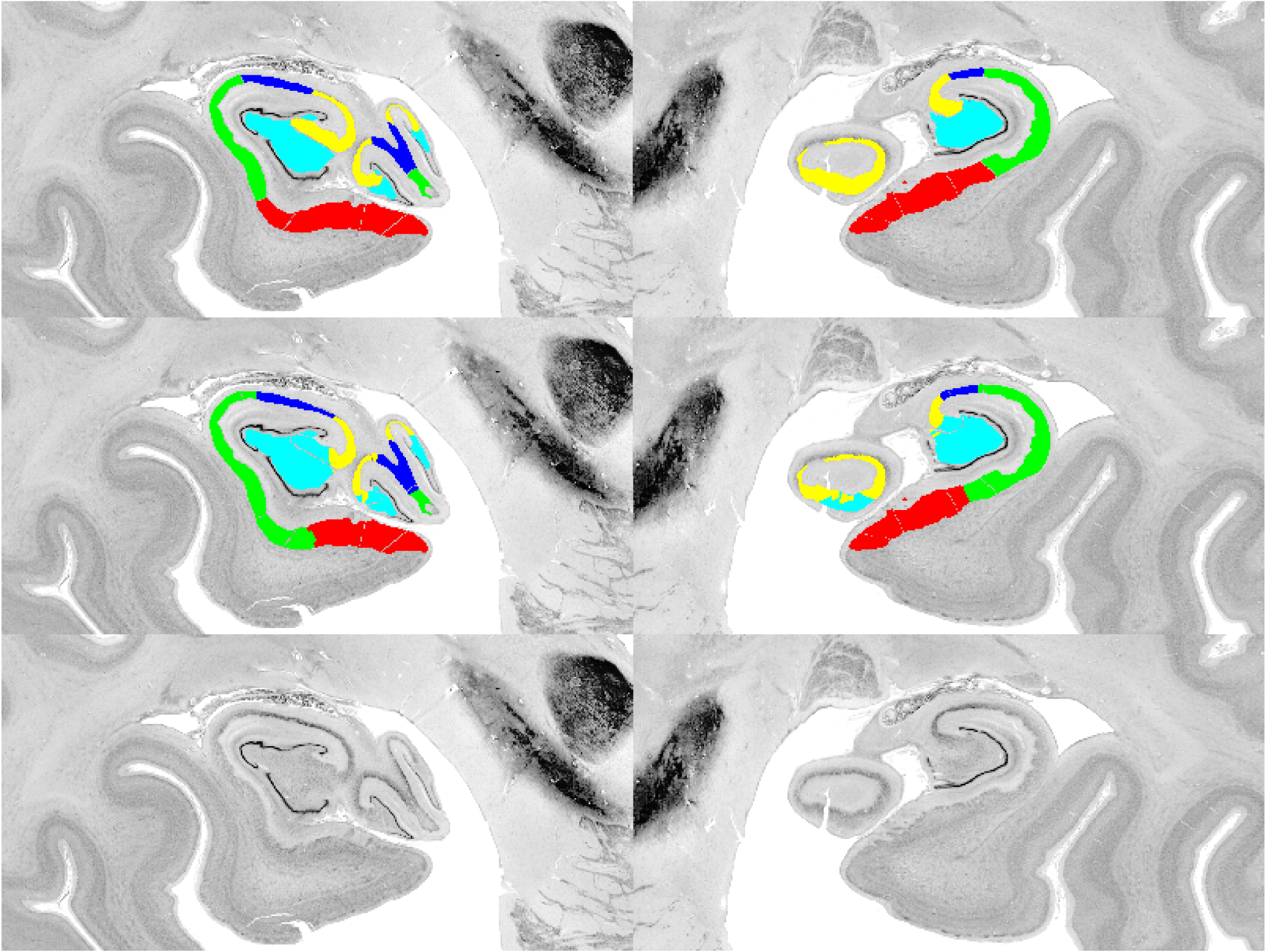

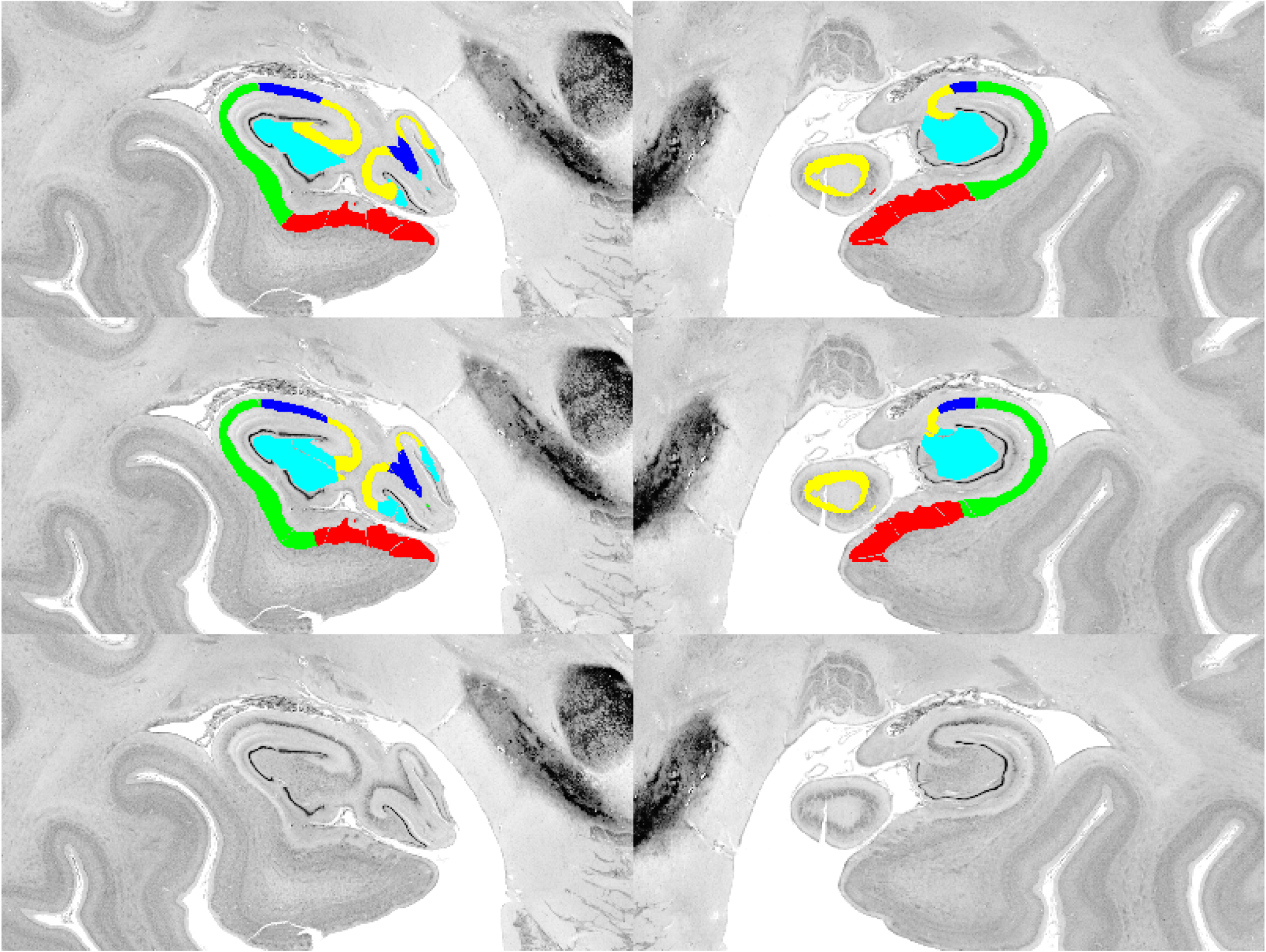

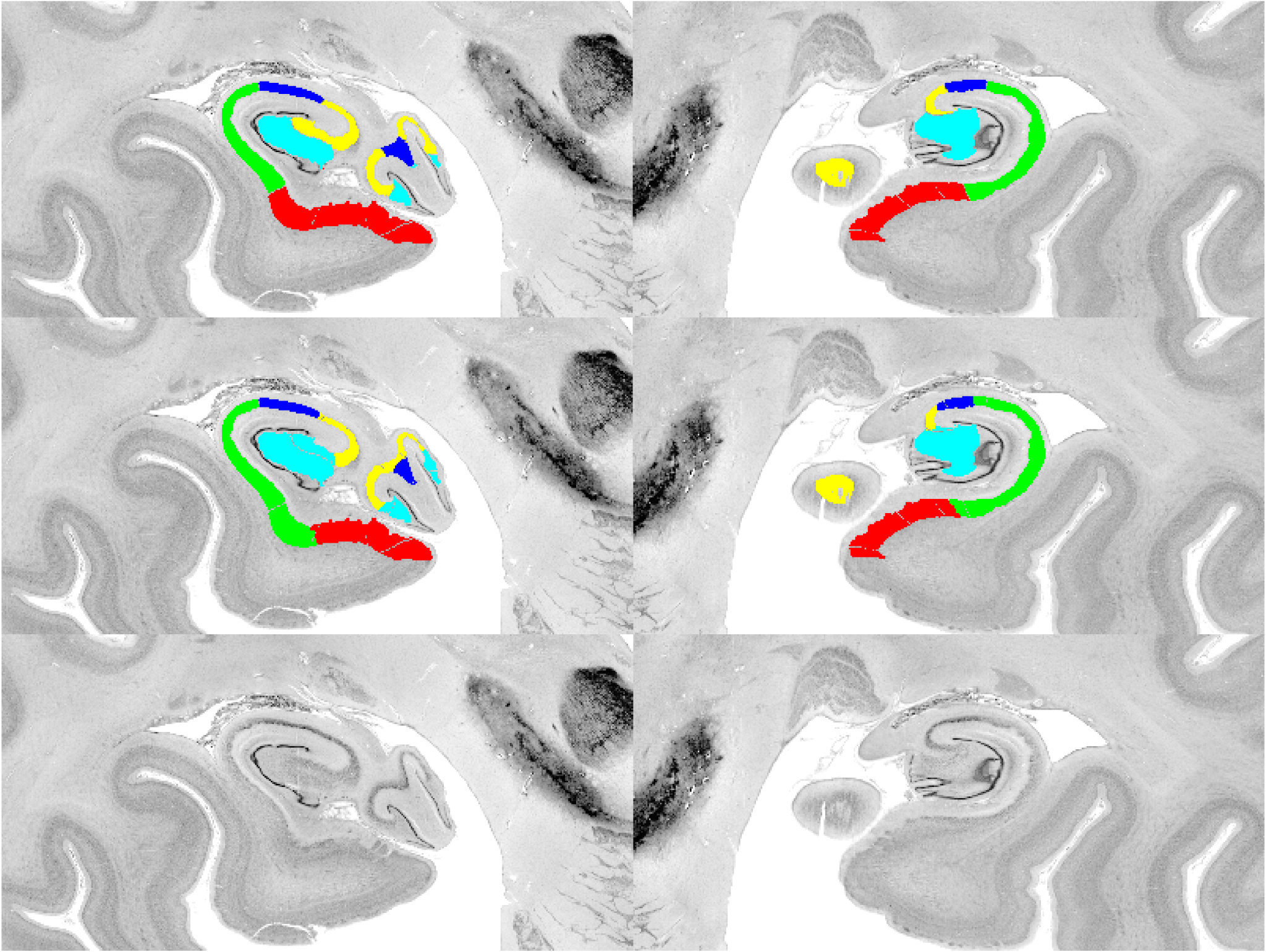

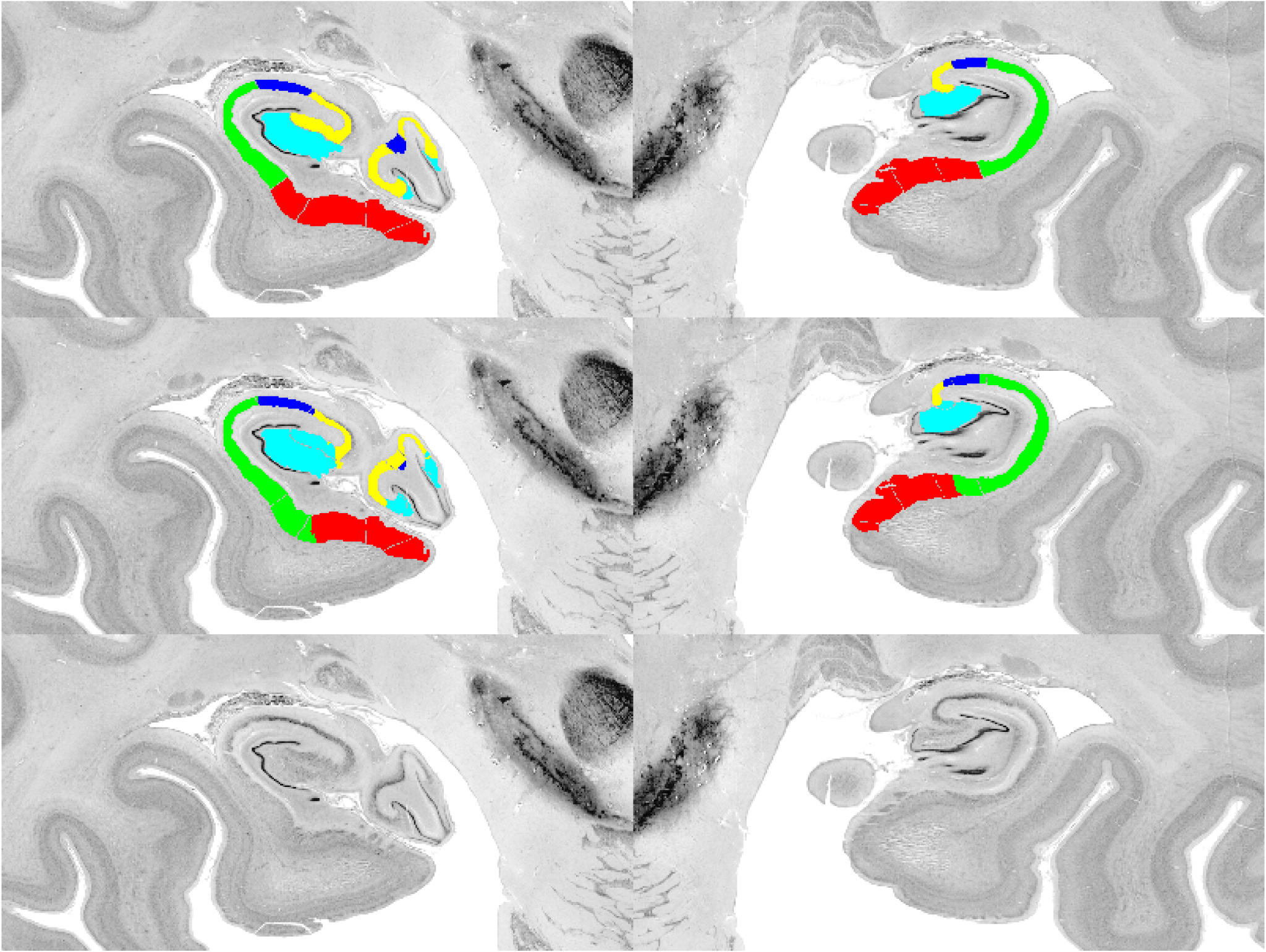

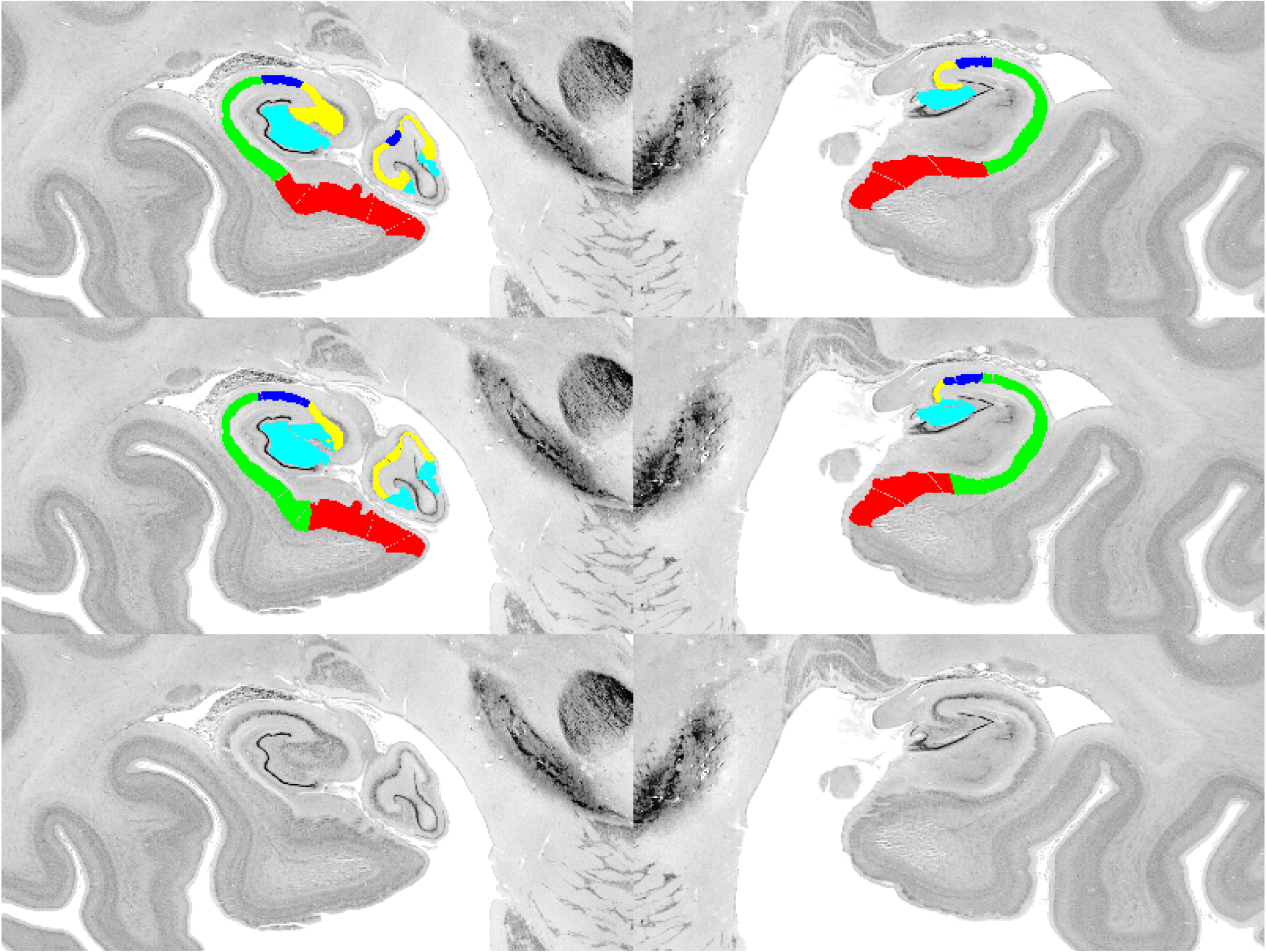

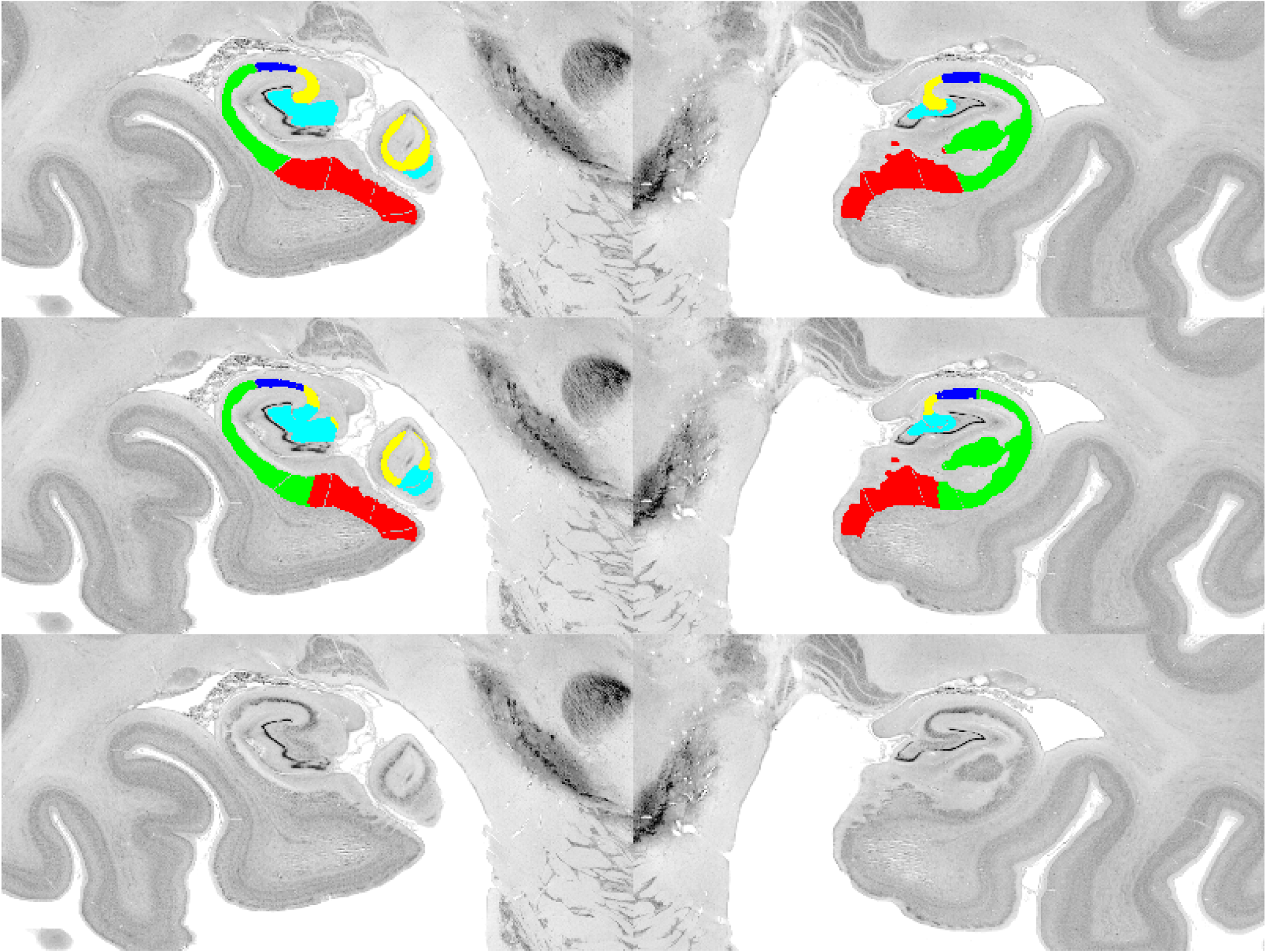

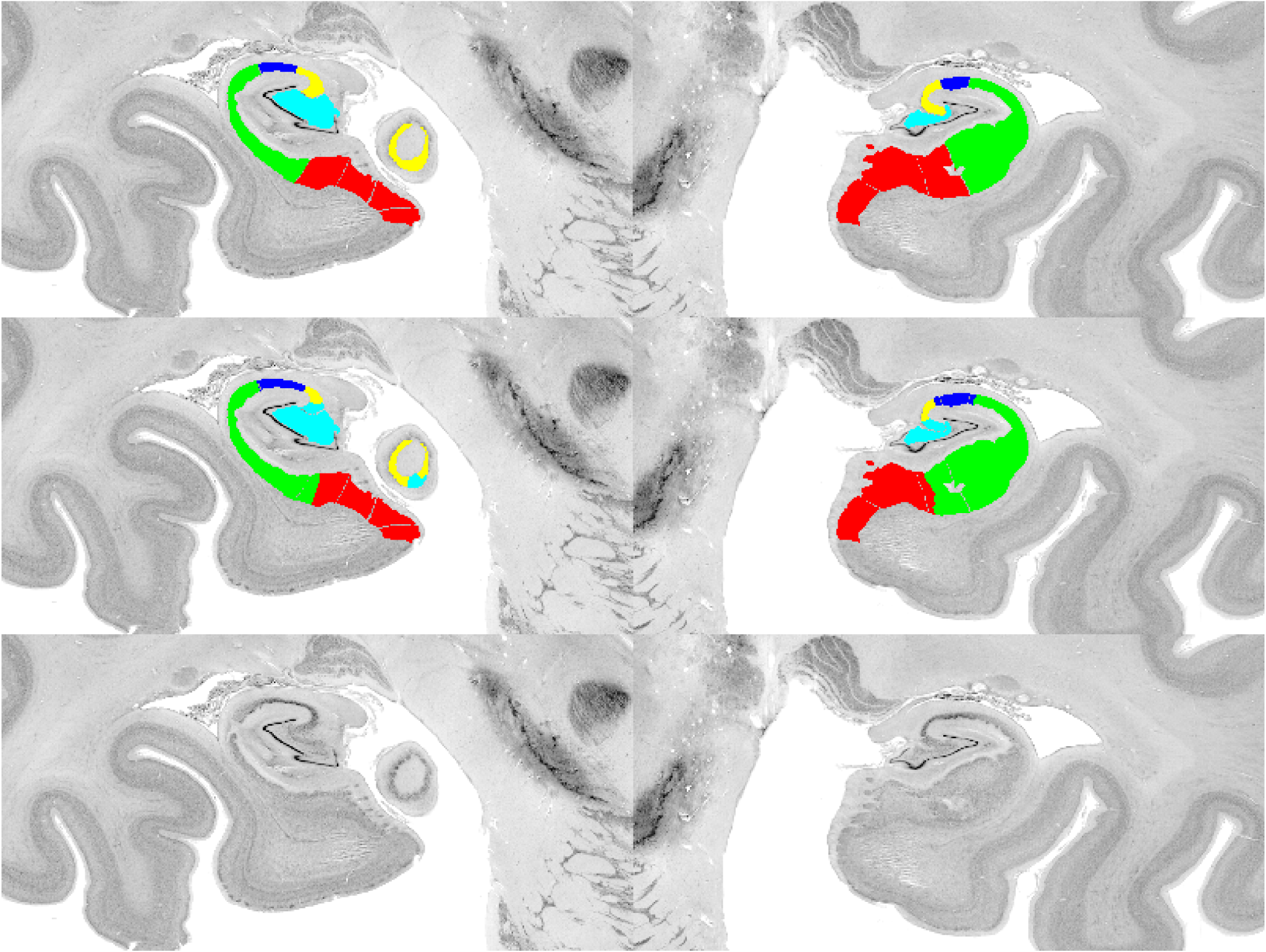

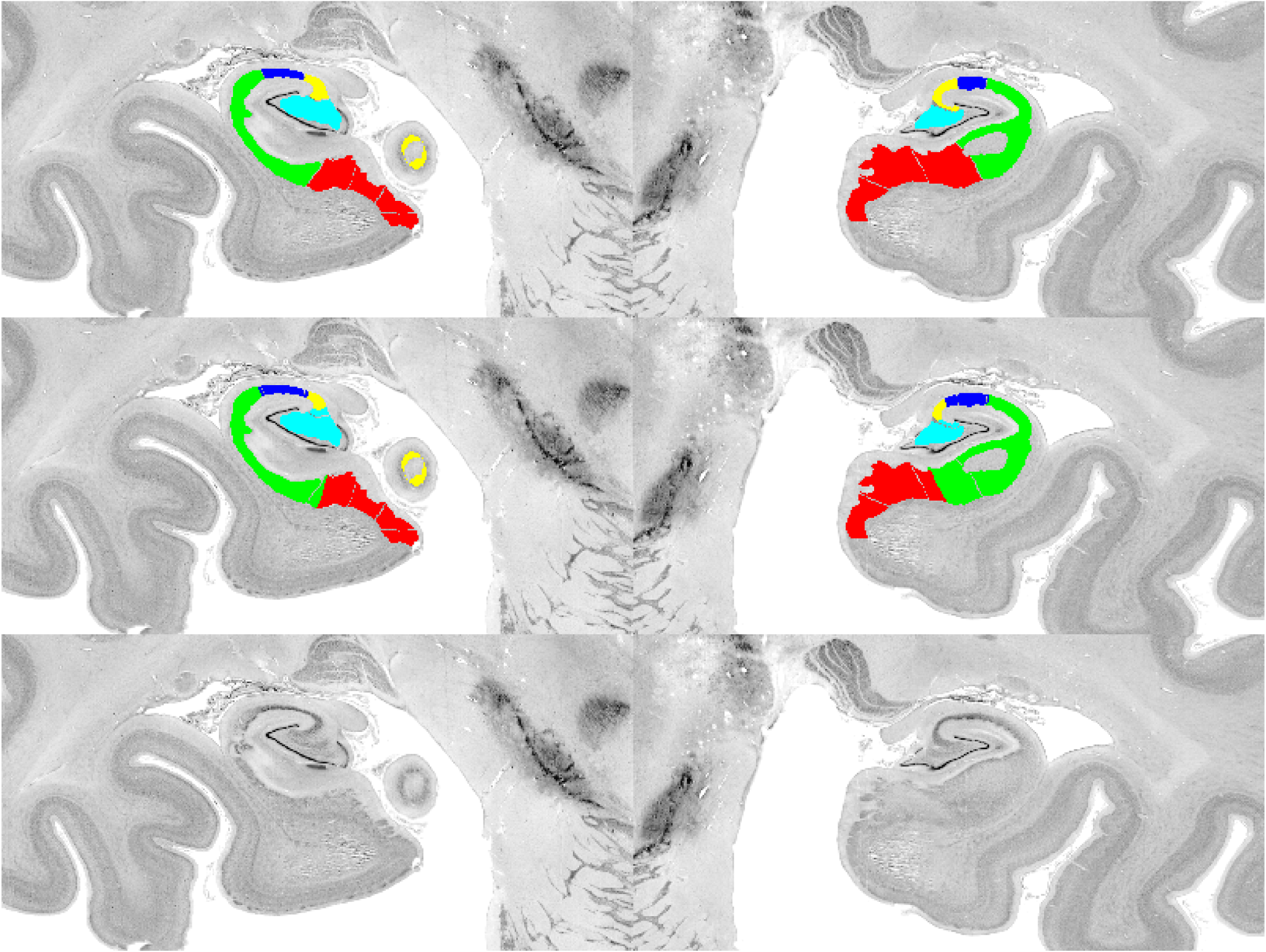

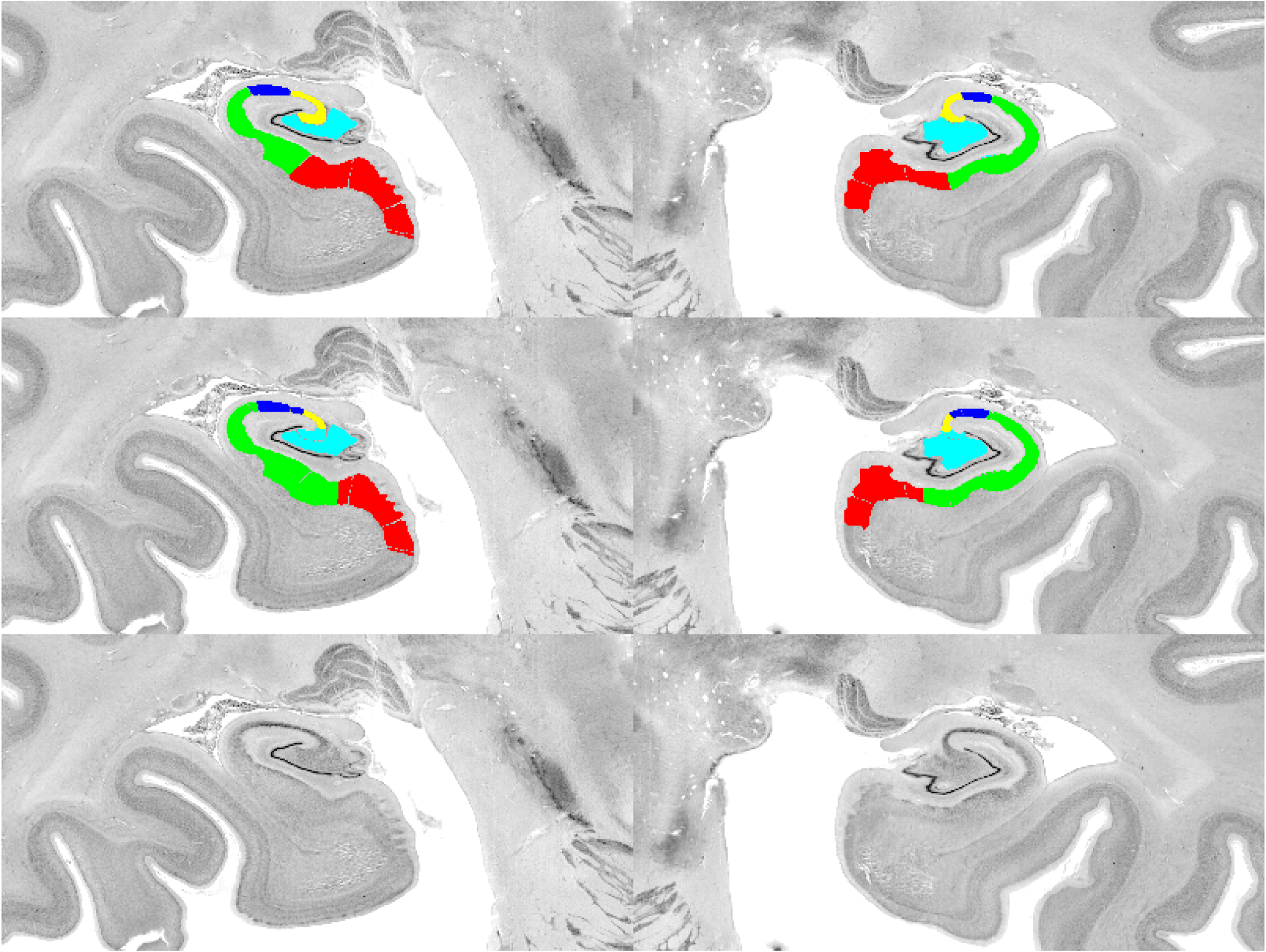

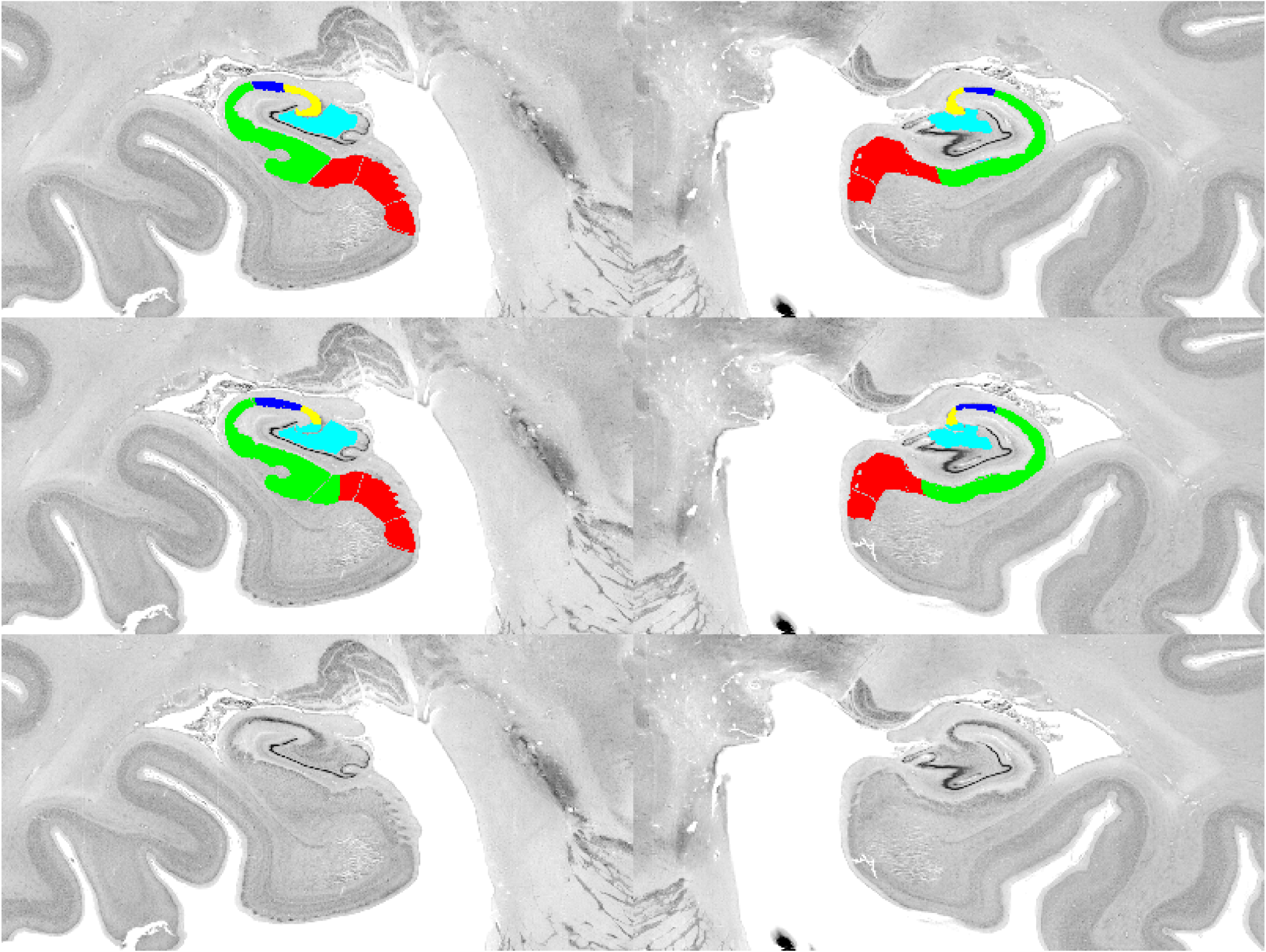

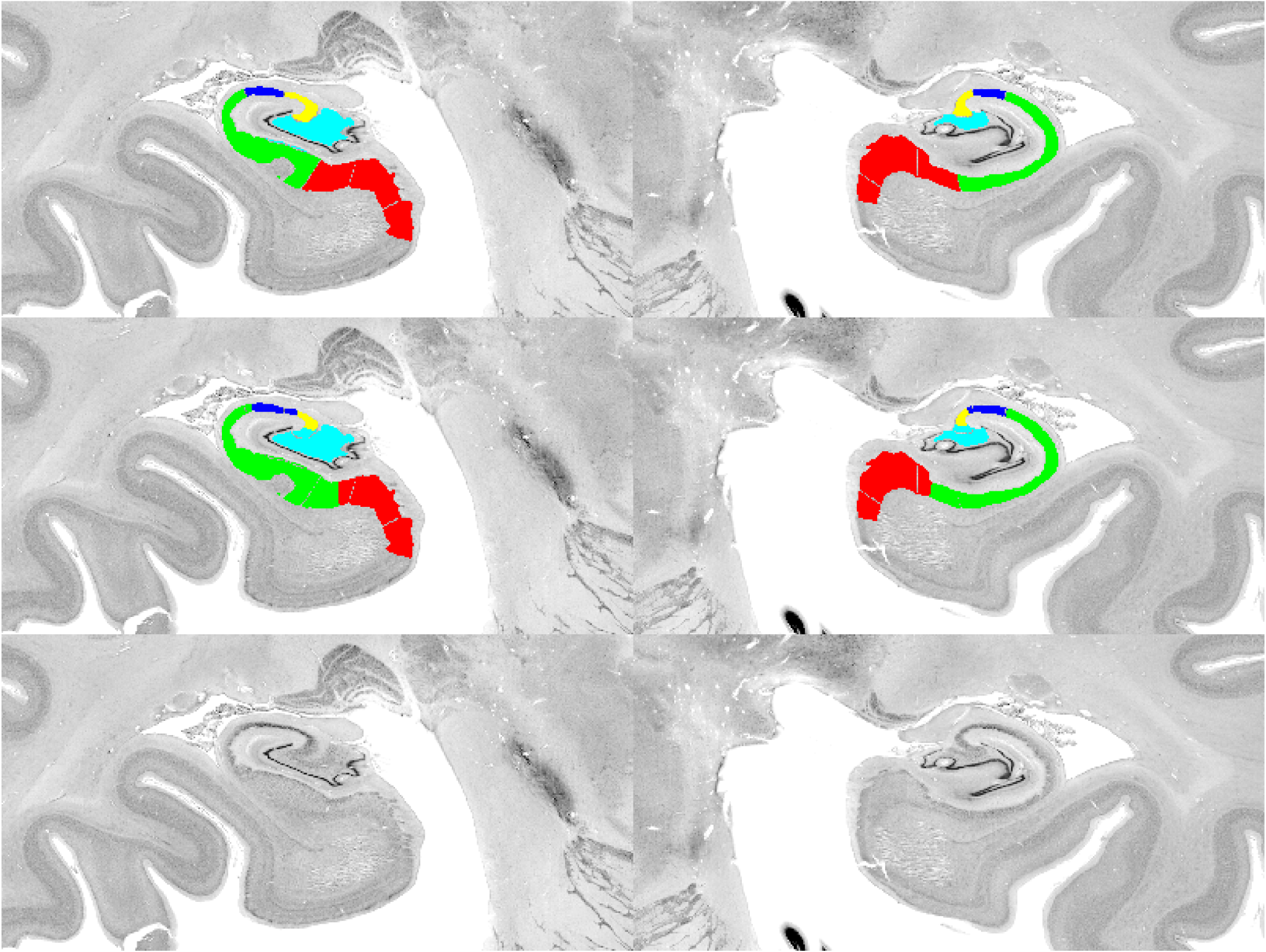

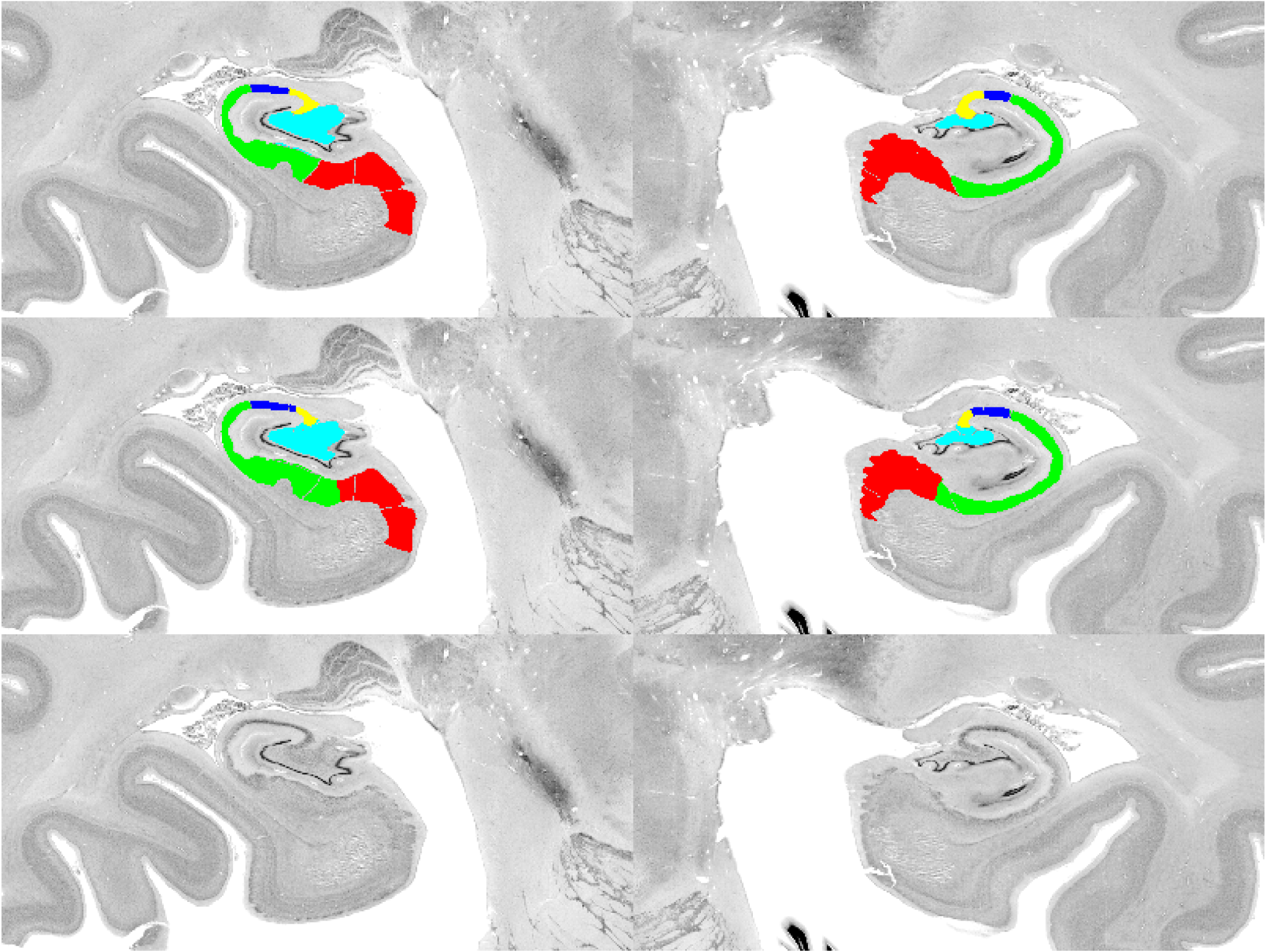

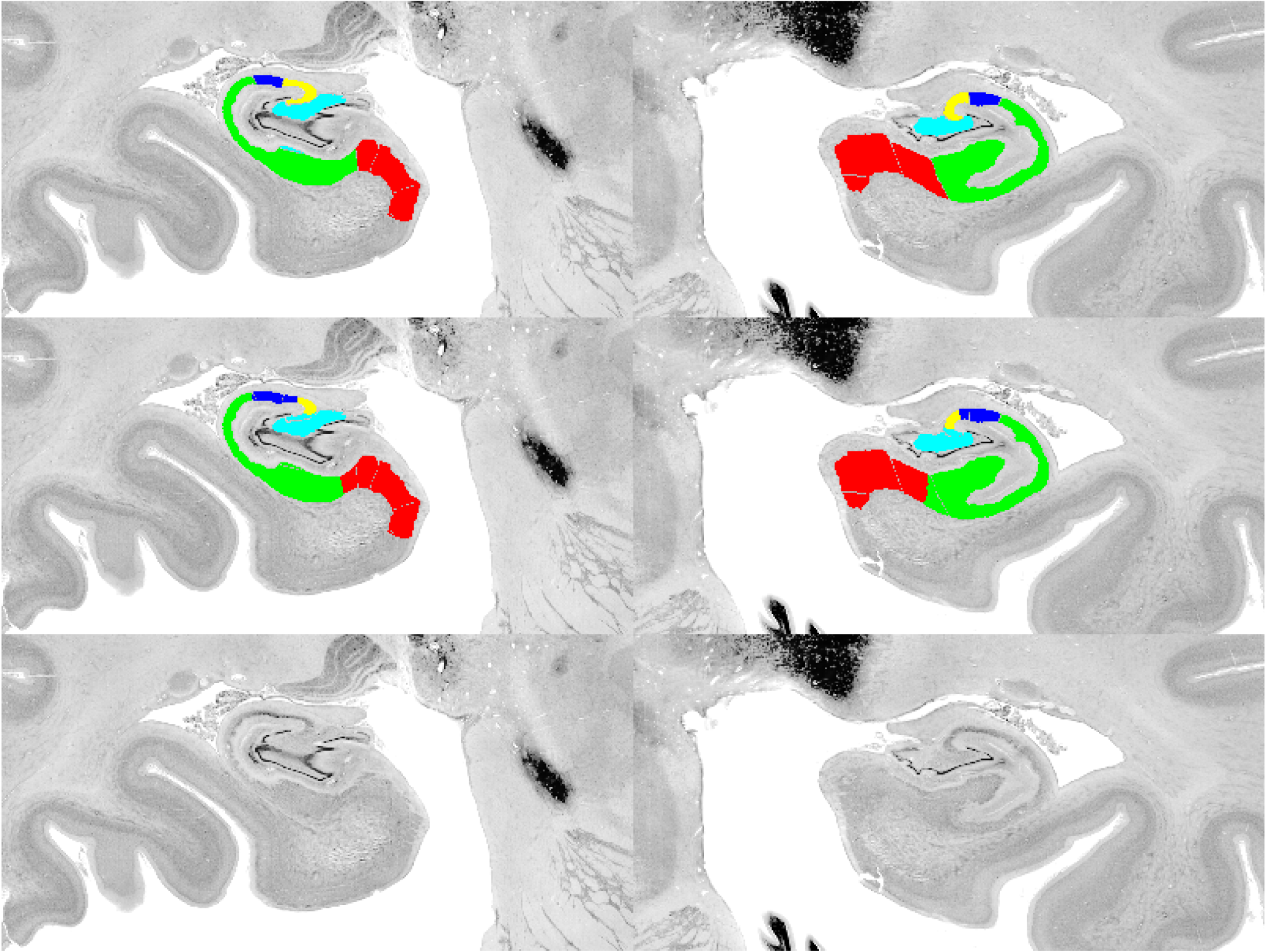

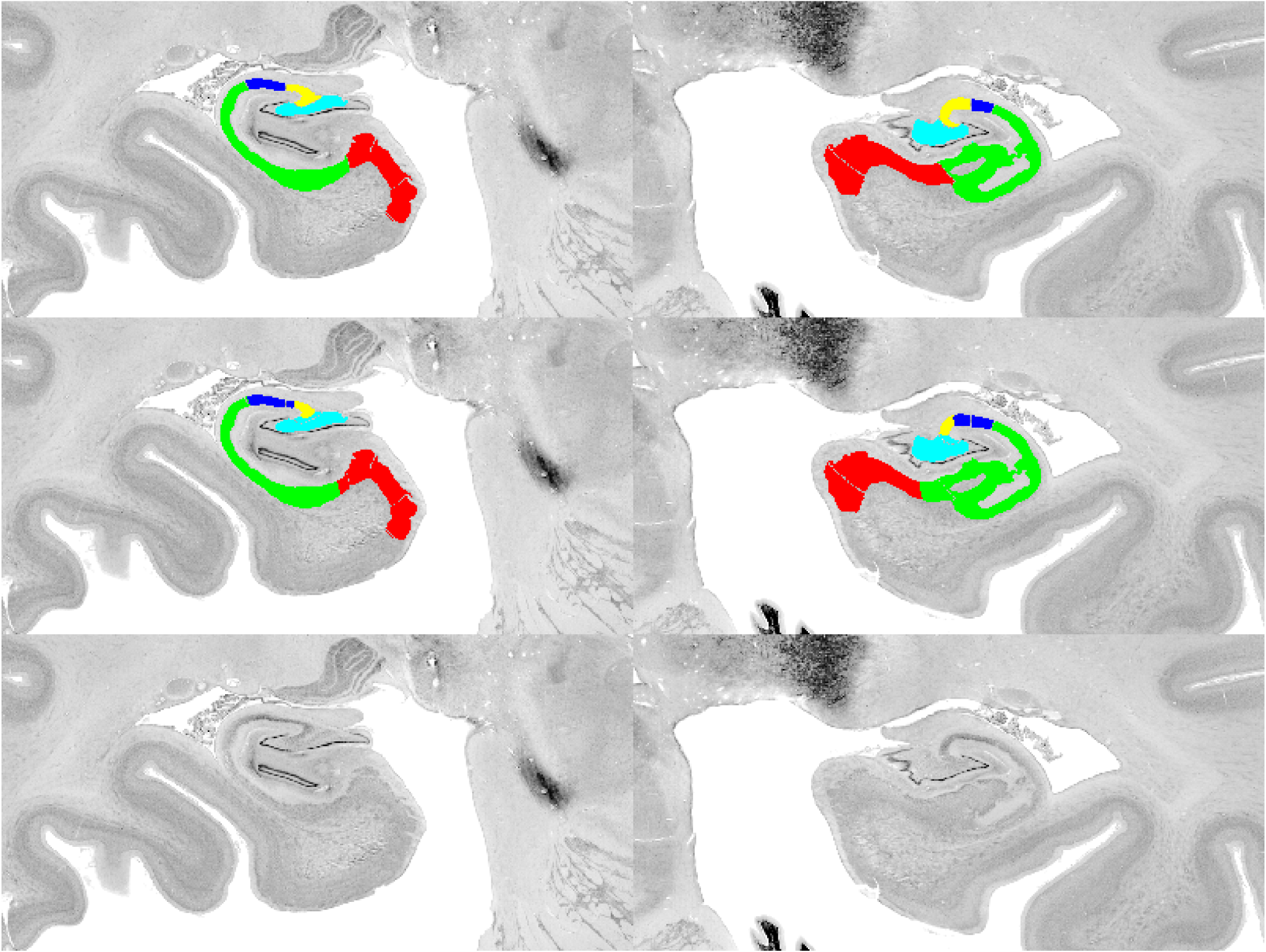

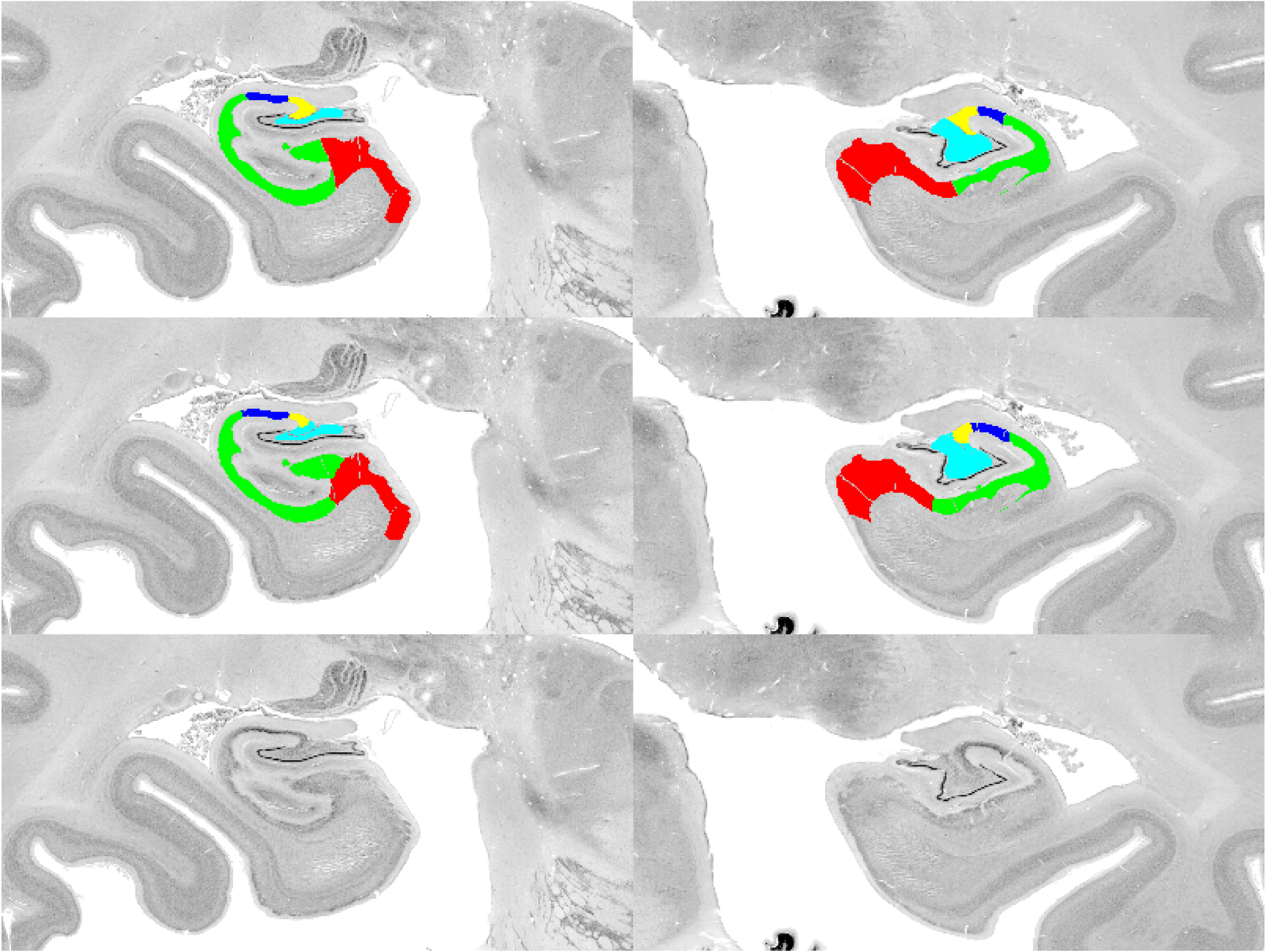

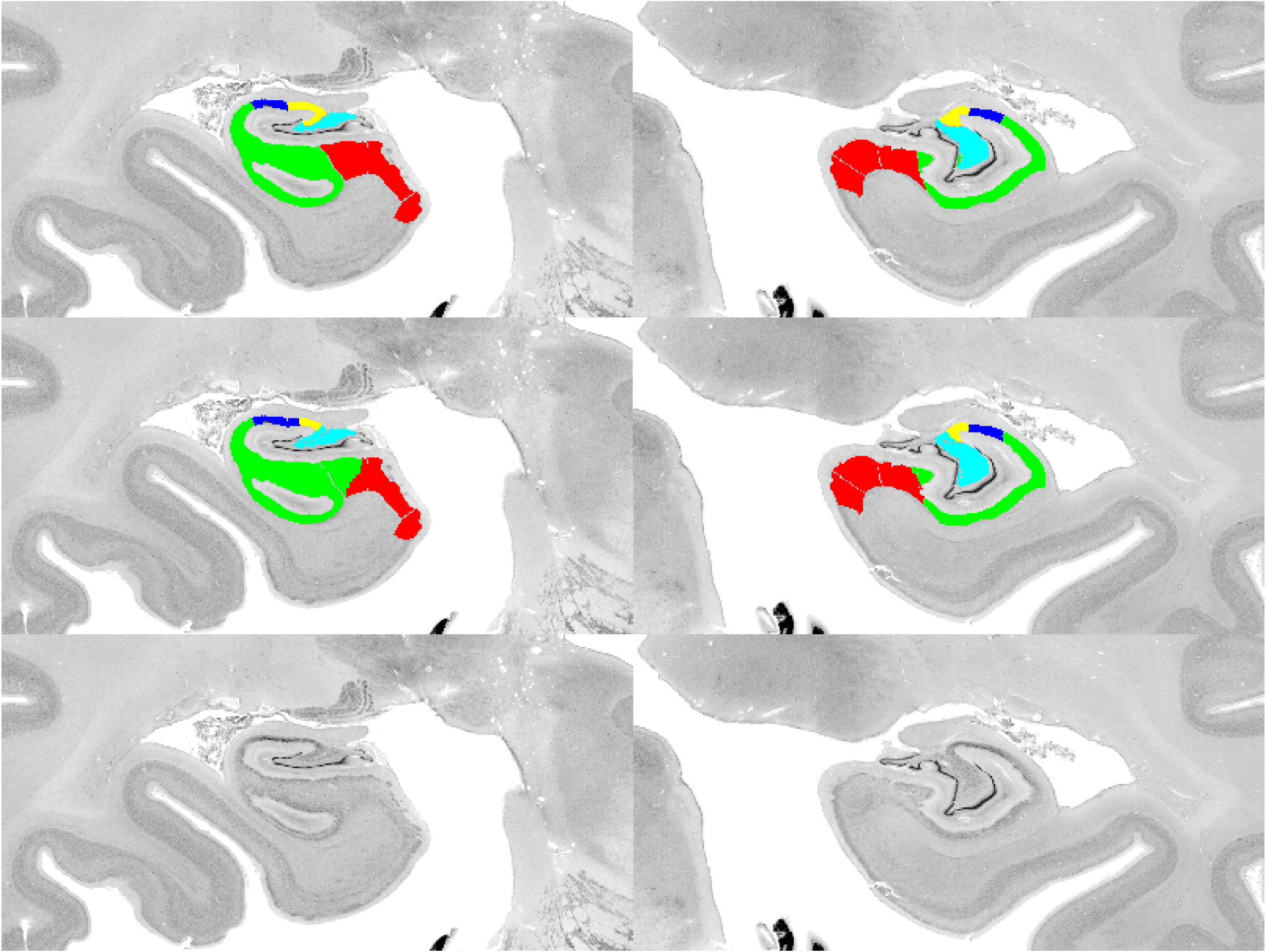

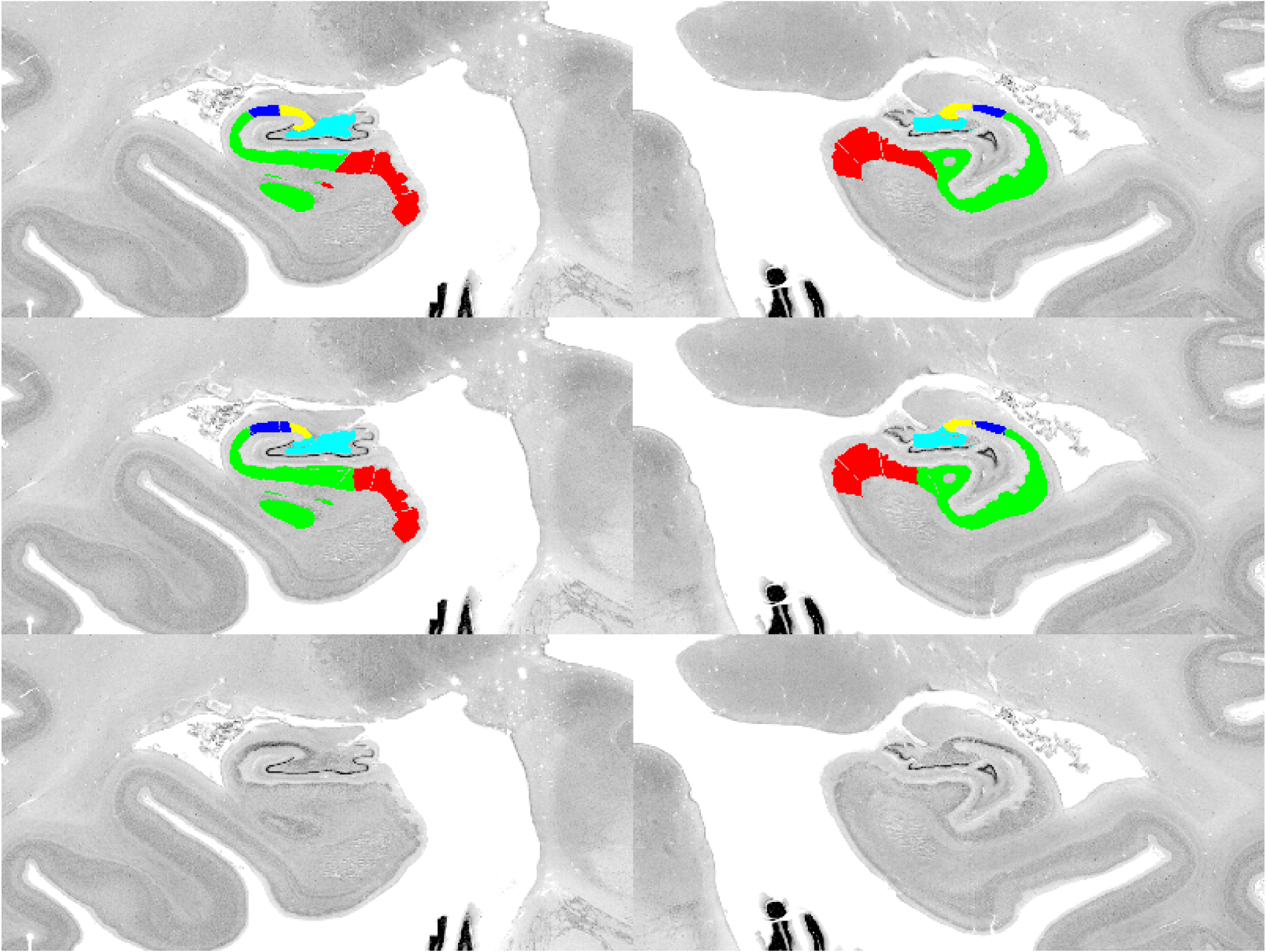

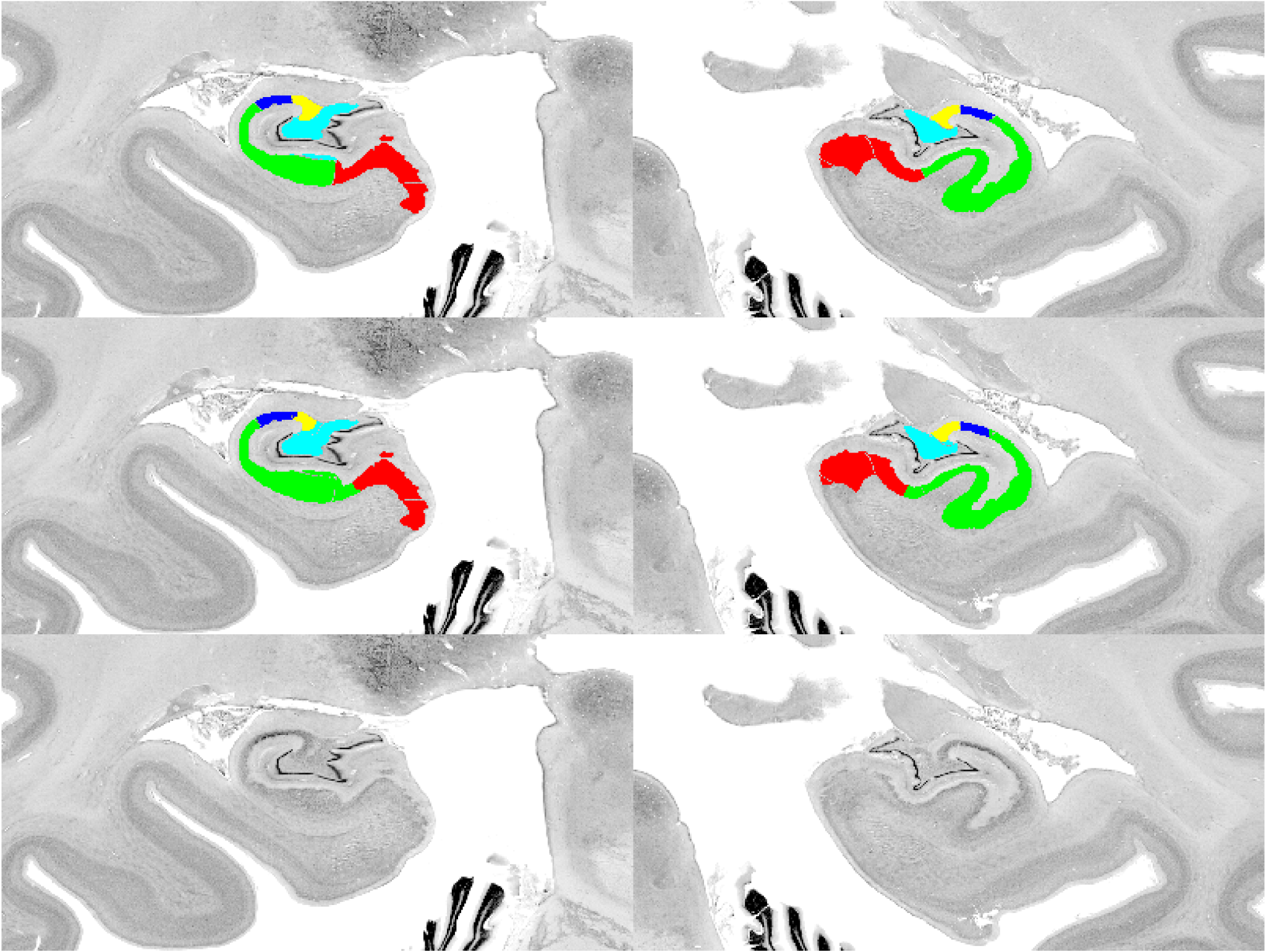

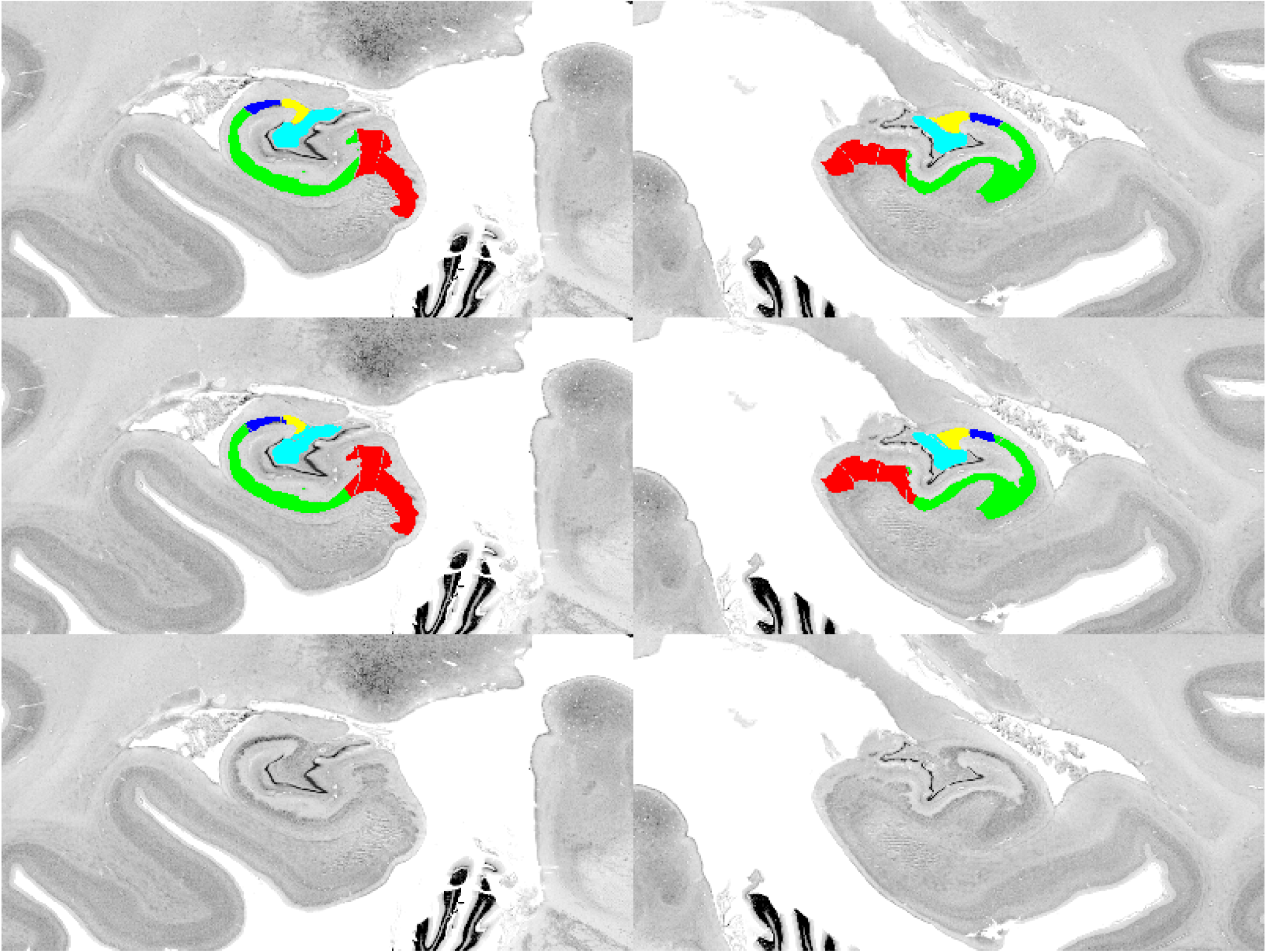

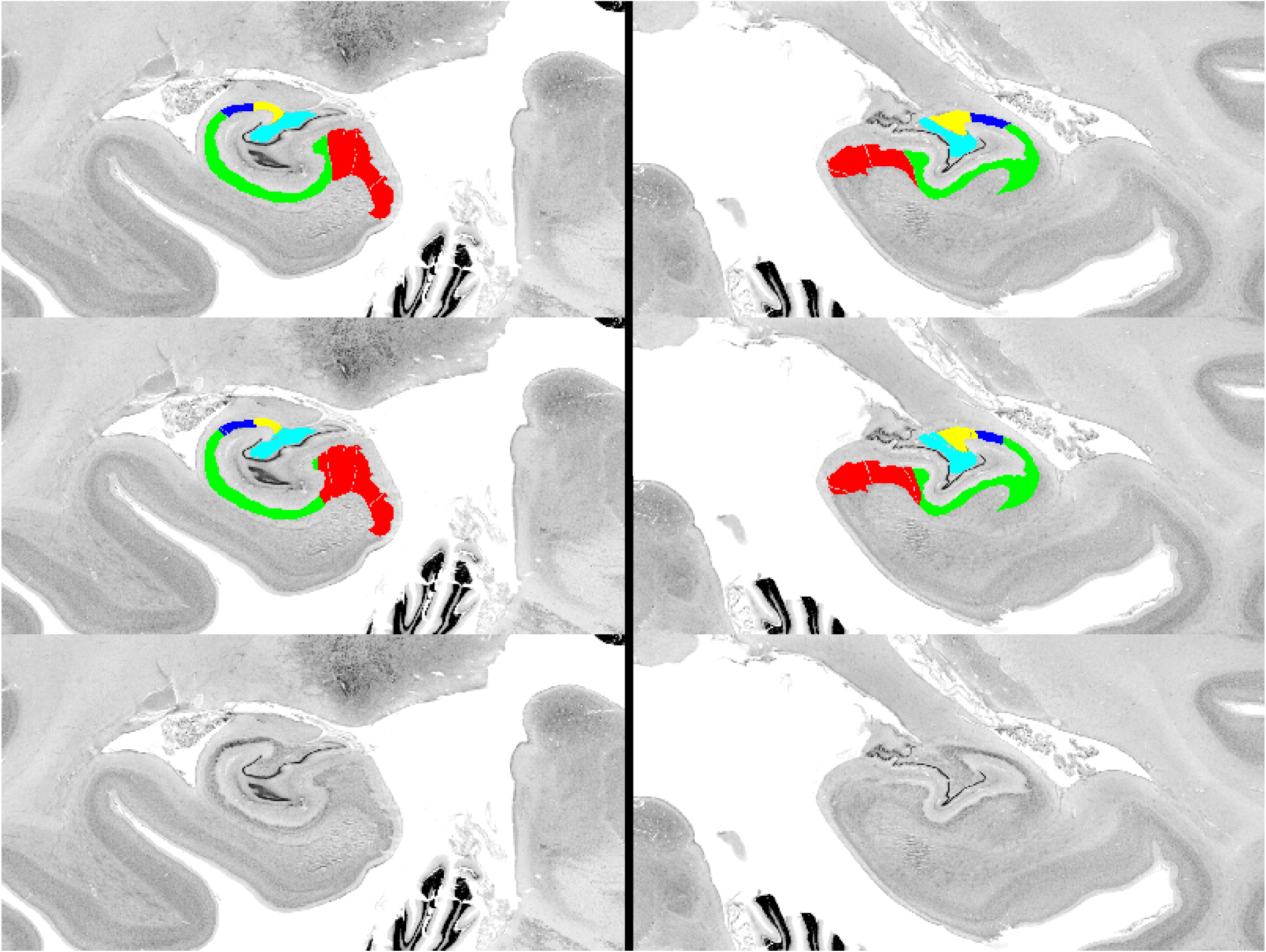

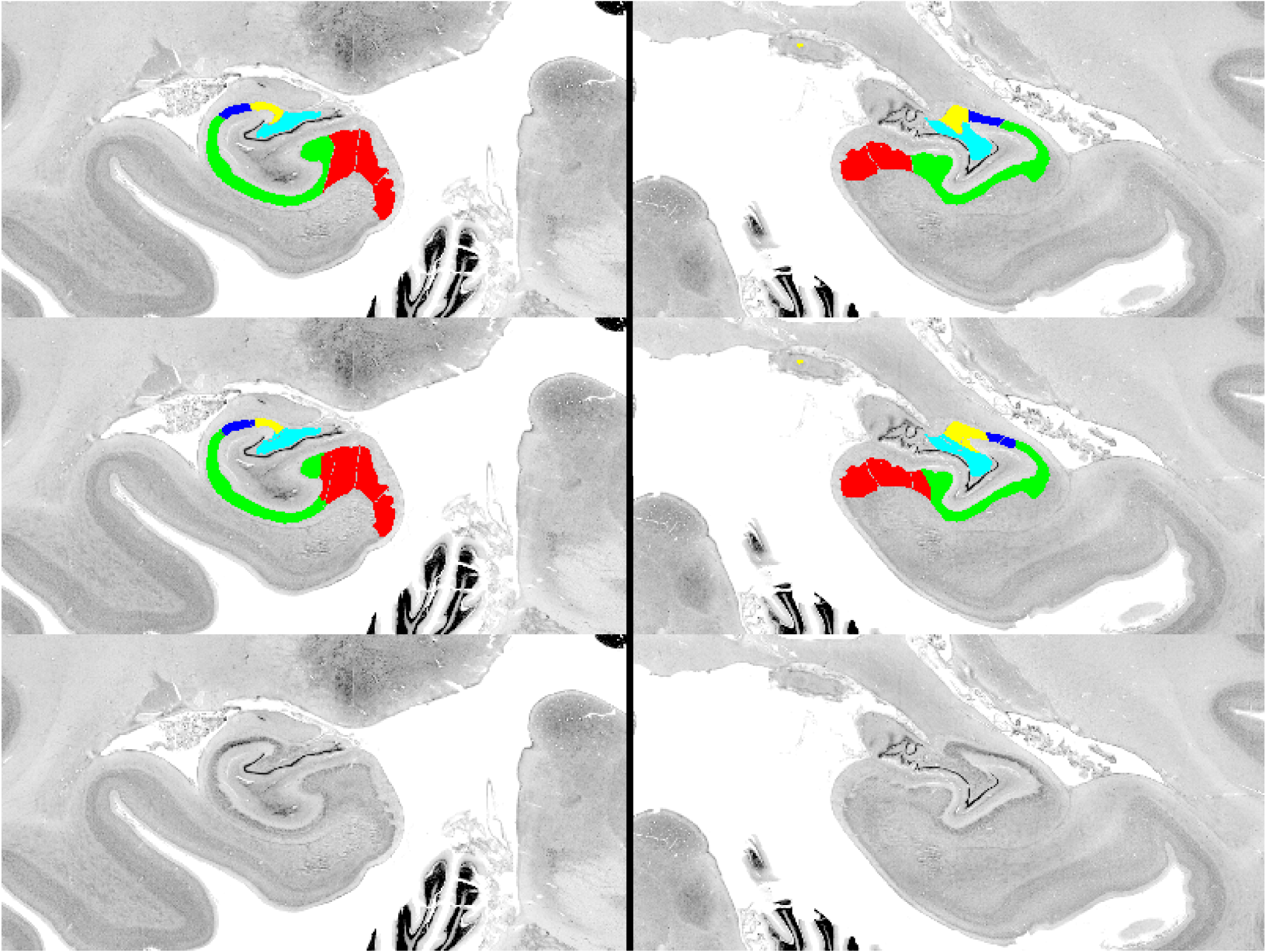

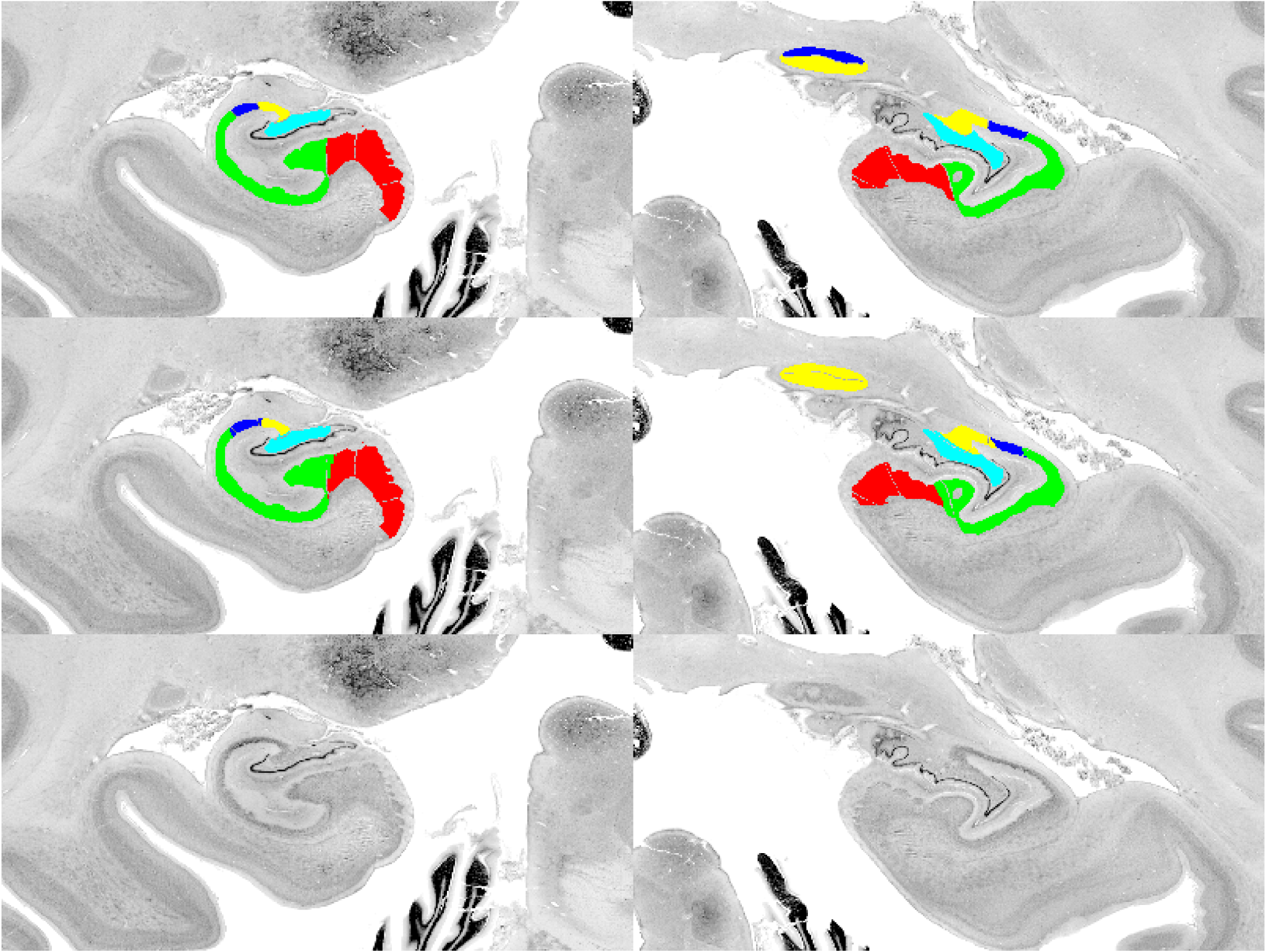

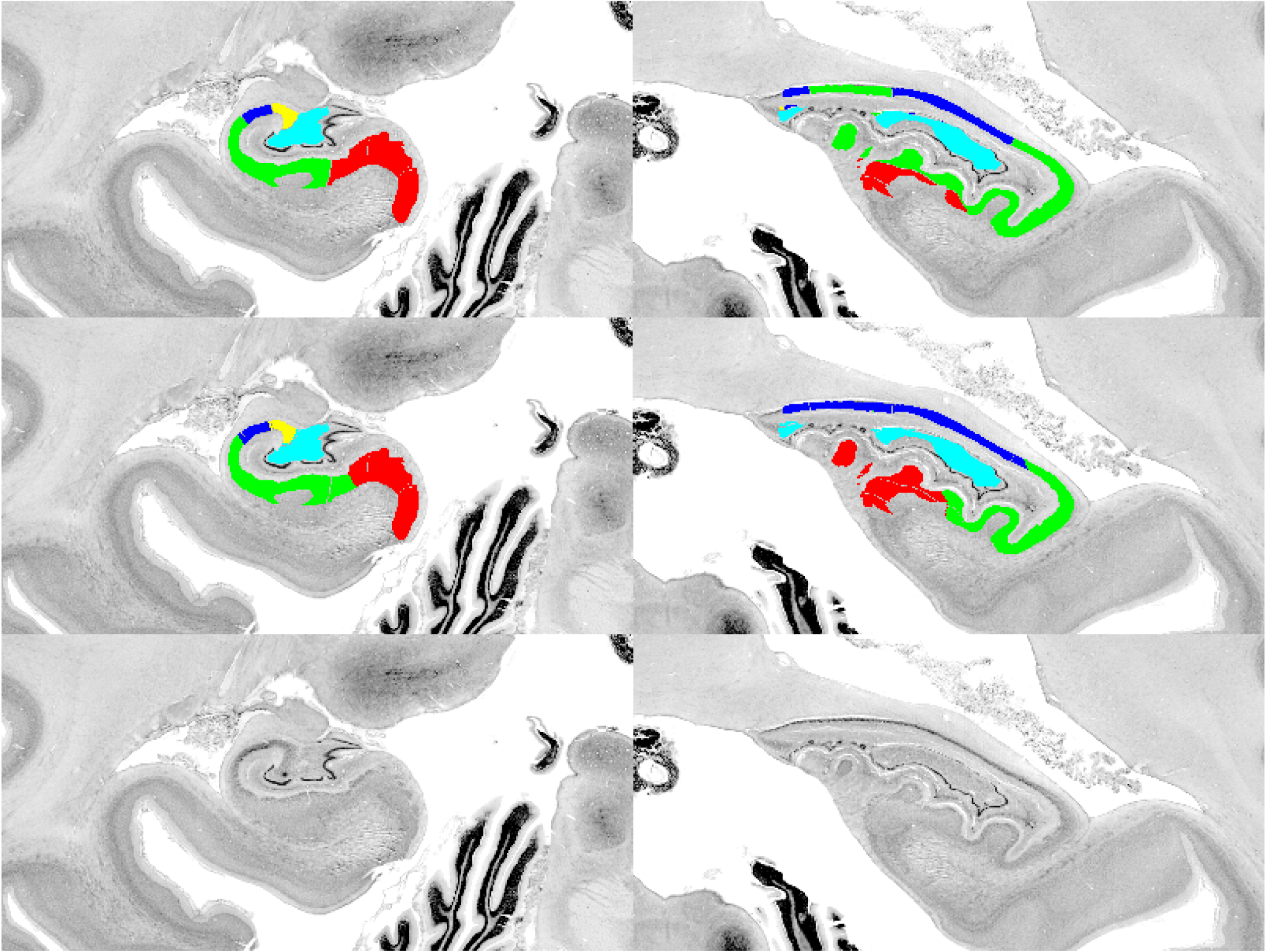

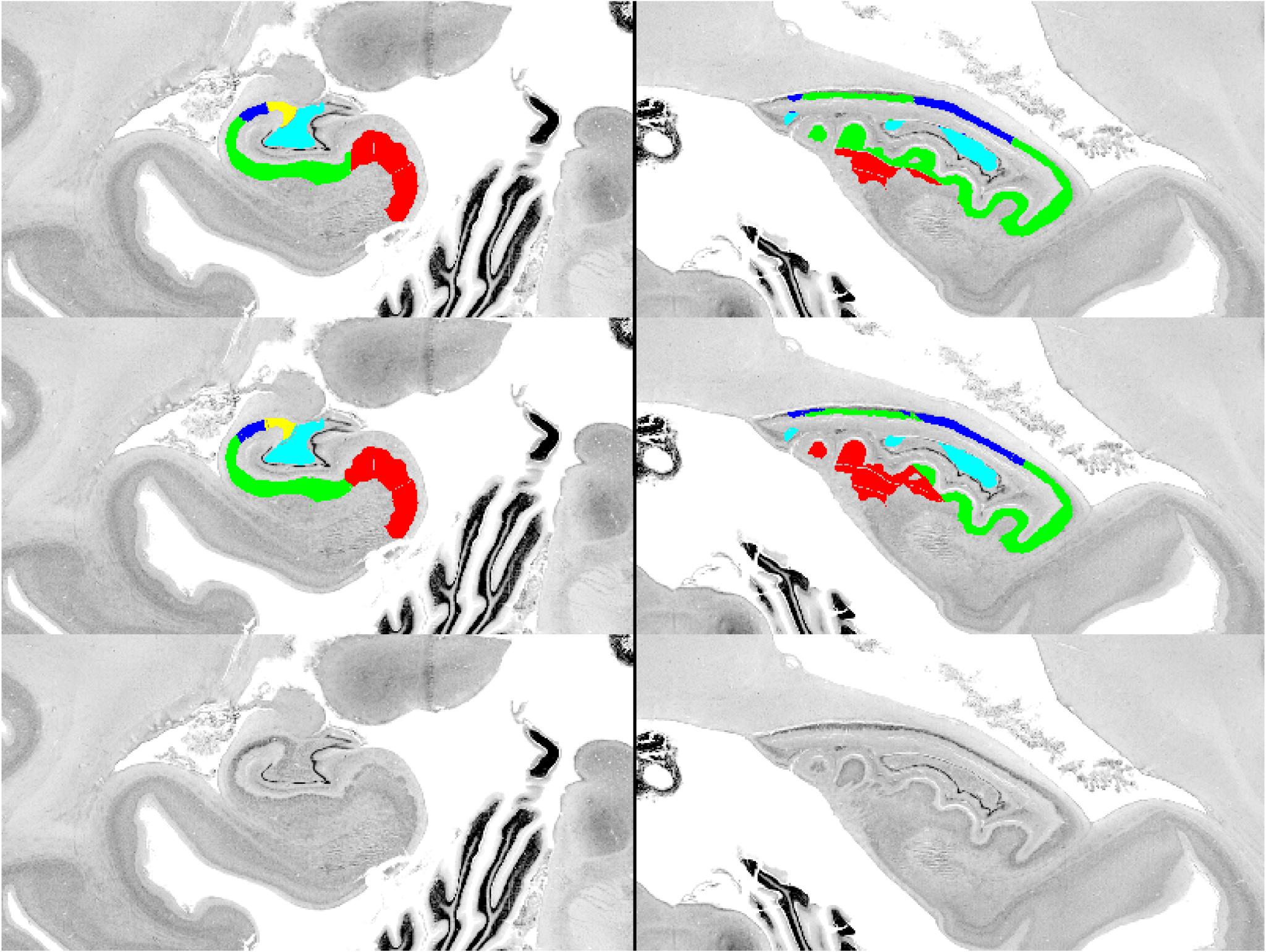

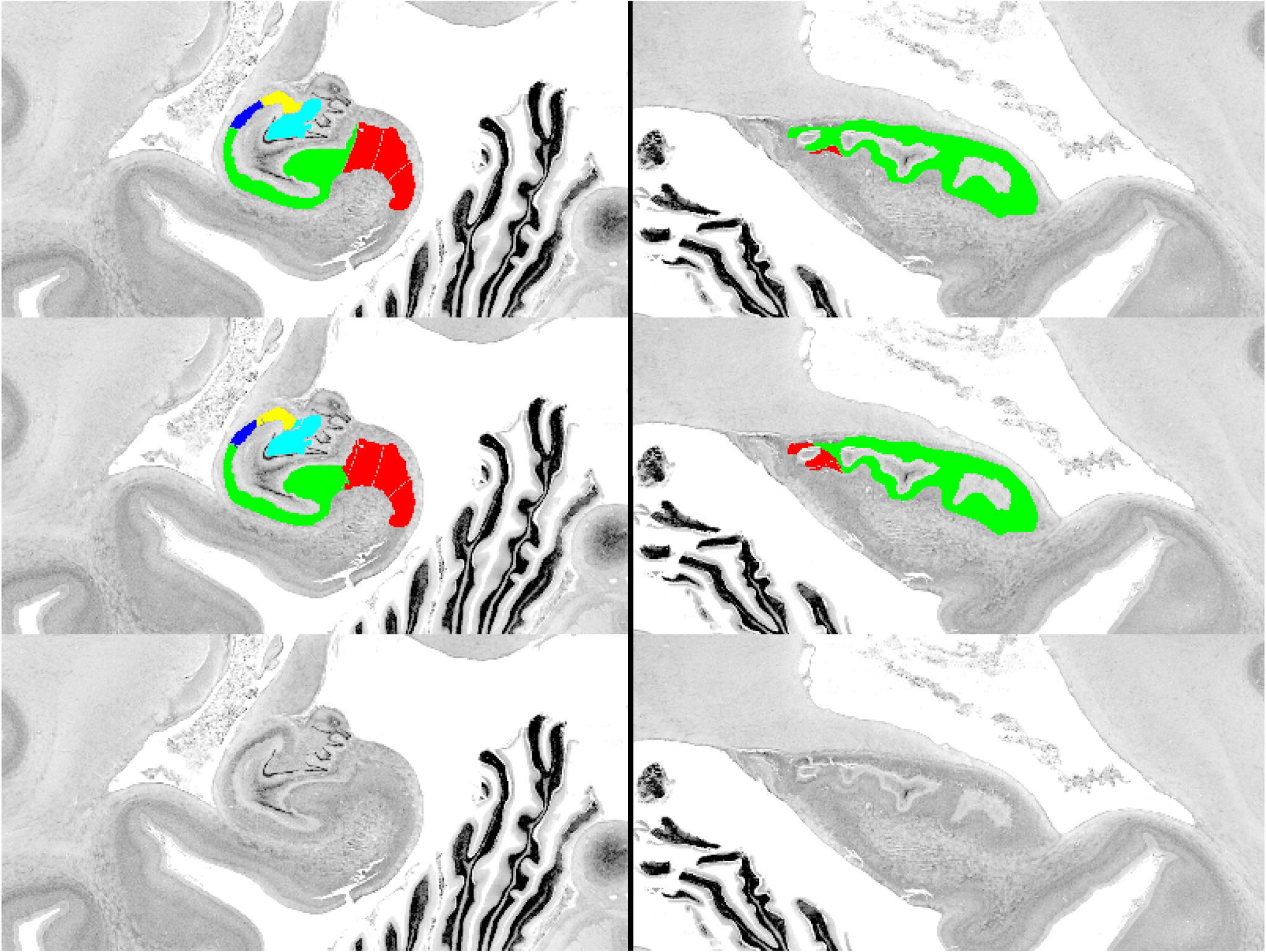

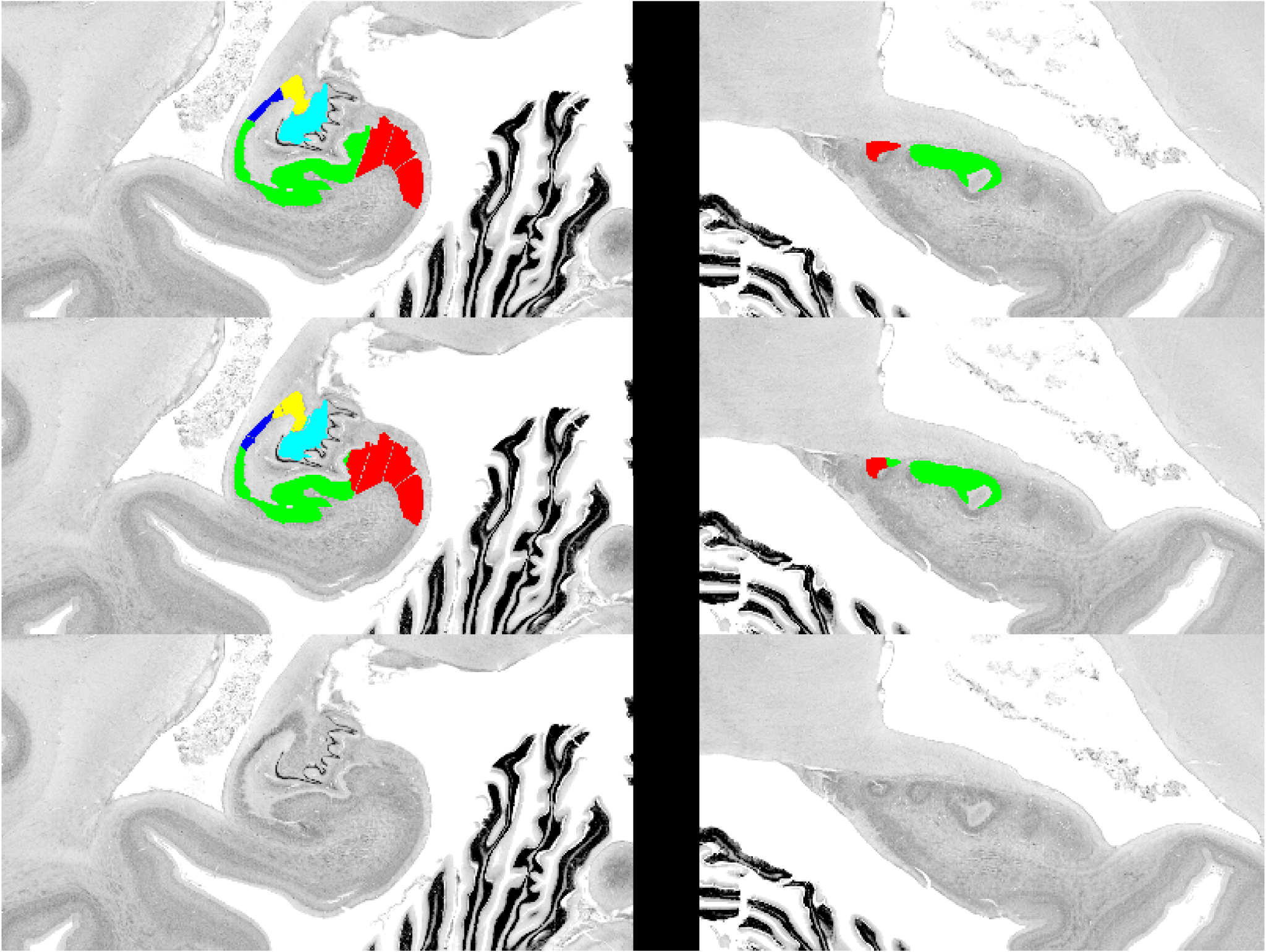

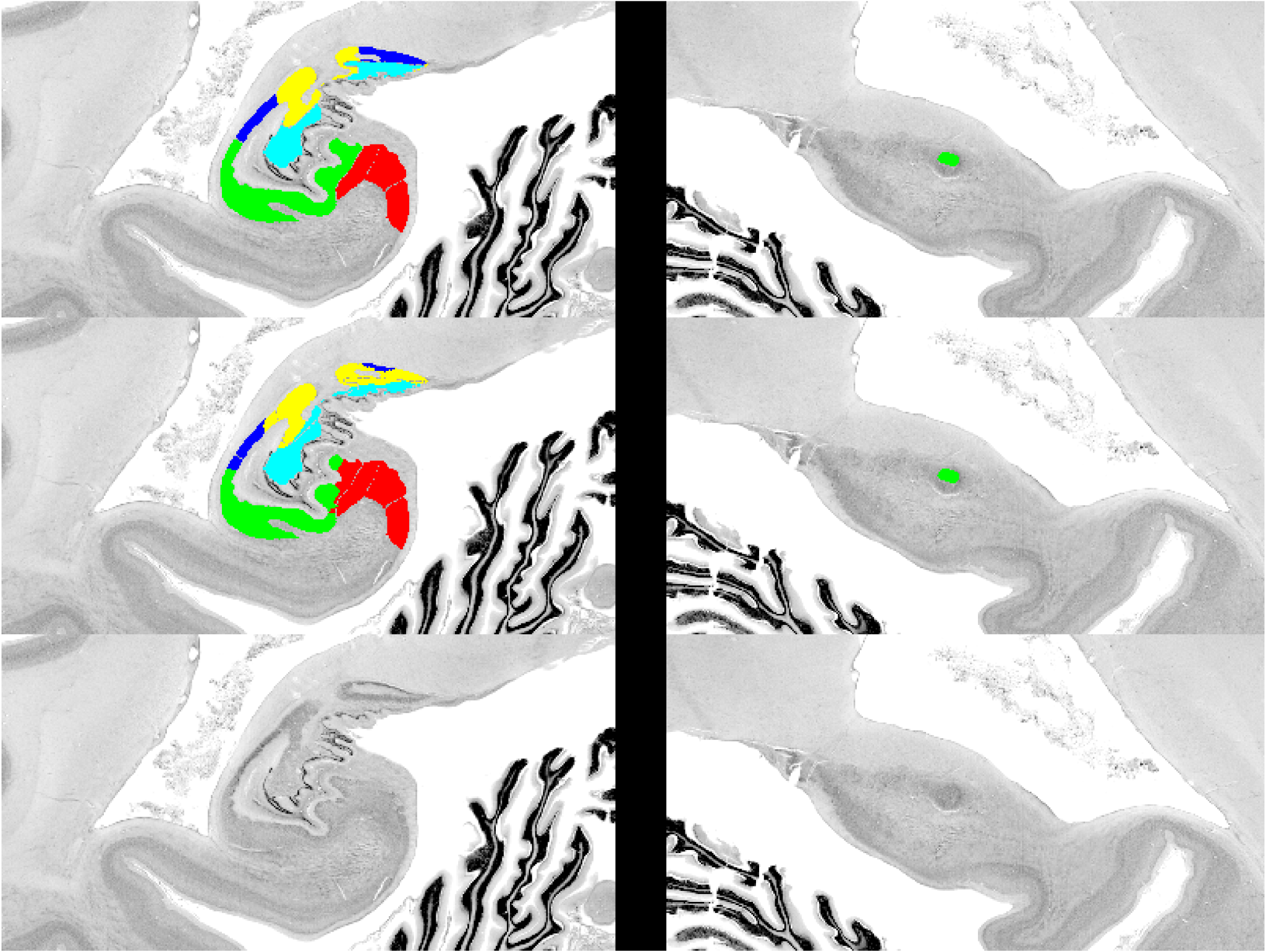
